# Neuromodulation of a peripheral nerve using fully polymeric cuff electrodes: Understanding predictability of selective stimulation

**DOI:** 10.1101/2025.10.28.685203

**Authors:** Zachary Bailey, Zachary Nairac, Estelle A. Cuttaz, Imane Ben M’Rad, Christopher A. R. Chapman, Adrien Rapeaux, Josef Goding, Aidan Roche, Timothy G. Constandinou, Rylie Green

## Abstract

Peripheral nerve stimulation (PNS) offers therapeutic benefits across numerous clinical applications but suffers from limitations in high resolution spatial selectivity, especially in mixed nerves. This study presents a fully polymeric, transverse, multipolar nerve cuff made from a conductive elastomer (CE), designed for selective activation of individual fascicles. Fabricated with laser-based manufacturing techniques, the CE nerve cuff offers mechanical conformity and high charge injection capacity. *Ex vivo* experiments on the rat sciatic nerve demonstrate reliable compound action potential recordings and fascicular selectivity (SI > 0.65). The non-metal electrodes enable microCT-aided 3D reconstruction of nerve-electrode geometries without imaging artefacts, informing anatomically accurate simulations via the ASCENT pipeline. While *in silico* simulations predict some selective fascicular activation, discrepancies were observed between predicted and experimental selectivity magnitudes and electrode positions, particularly for the sural and tibial fascicles. The model was more sensitive to neuroanatomical variation than the experimental data, indicating limitations in current perineurium and CE electrode modelling assumptions. This work validates CE-based cuffs as viable alternatives to metallic devices for selective fascicular peripheral nerve activation and highlights the potential of imaging-informed simulations to optimize nerve interface design. Future improvements in electrode and nerve tissue modelling are needed to enhance *in silico* prediction accuracy and further advance spatially selective PNS technologies.

## Introduction

Peripheral nerve stimulation (PNS) has enabled life-improving treatment in millions of patients, spanning chronic neuromuscular and neurological pain relief,^1^ enhanced neuromuscular rehabilitation,^2^ and muscle atrophy prevention following nerve injury.^3^ Most of these therapeutic interventions occur via transcutaneous electrical nerve stimulation (TENS), which is a non-invasive approach with minimal risk to the patient. However, this approach typically results in bulk stimulation of the entire mixed nerve bundle, with selectivity of the therapy being limited to placement of the electrodes on the skin at an appropriate location overlying the target tissue. Implantable systems have also been developed that stimulate peripheral nerves directly through a nerve cuff that wraps around the nerve. This direct stimulation lowers the activation threshold for inducing an action potential and decreases off target muscle stimulation, improving therapeutic outcomes and lowering the power requirements for activation, making small, implantable systems more effective and mobile compared to a TENS configuration. However, most commercial implantable devices used in the clinic are still providing bulk stimulation to the underlying peripheral nerve, resulting in unwanted side effects and suboptimal therapeutic response. This has hindered the uptake and utility of these potentially life enhancing technologies. Consequently, significant research effort over the past two decades has sought to develop more advanced devices and stimulation paradigms for improving the targeting of neural subpopulations in peripheral nerves.

Experimental clinical trials exploring more selective stimulation of subpopulations of fibers within a mixed nerve have shown the potential for tailored treatments including precise bladder control, circulatory and respiratory system regulation,^4^ movement restoration after spinal cord injury,^2^ and advanced prostheses control/feedback loops for amputees.^5^ However, the benefit of more selective stimulation must be worth the substantially increased risk of an implantable medical device for direct nerve stimulation. A critical aspect of PNS that predicts relative patient benefits in treatment is selective activation of subpopulations of fibers within a peripheral nerve. While high resolution fiber activation is often the primary objective of innovative peripheral nerve stimulation implants, without precision, there is an associated risk of off-target stimulation that could worsen patient outcomes if the mixed nerve innervates sensitive, critical organs, such as the vagus nerve innervating the respiratory diaphragm or heart.

To improve the resolution of fiber activation in PNS, a range of engineered electrode arrays have been designed to access targeted subpopulations of fibers. These approaches have sought to either penetrate the epineurium and/or the perineurium or mechanically compress the nerve into a flattened shape.^6^ While this does increase resolution, it also causes chronic inflammatory reactions and significant deformation.^7^ Therefore, these intraneural and deformation approaches have not yet achieved broad clinical uptake. Multipolar peripheral nerve cuffs offer a lower risk extraneural option that enables improved targeting of nerve subpopulations while having reduced mechanical impact on the nerve biology. Multi-ring electrodes that wrap around the entire nerve bundle enable some basic steering of charge and activation based on penetration depth of stimulation. These electrodes are proficient in targeting groups of axons based on their fiber type, not location, because of the well-defined relationship between activation thresholds and frequency of stimulation in fibers that differ by fiber diameter and degree of myelination. While fiber type specific stimulation is currently clinically useful in treating chronic pain and epileptic seizures,^8,9^ there is a need to spatially target fascicular and subfascicular groups of axons to control autonomic, motor unit, or somatosensory responses in a controlled and biologically meaningful manner. This study brings together multipolar electrode array design, *in silico* simulation, and experimental *ex vivo* stimulation across a wide range of paradigms to understand the most influential stimulation parameters in selective activation of fiber subpopulations by fiber location.

Previous studies undertaken on selective peripheral nerve stimulation have used conventional metallic-based electrodes in contact with nervous tissue. While metallic electrode arrays have known biological responses and are well tolerated in chronic implants, they present significant limitations to charge injection,^10,11^ especially when the target tissue is more distant from the electrode requiring higher currents to achieve therapeutic effects. Moreover, most metallic implants are stiff, and their rigidity affects their conformation to the neural tissue and creates gaps between the electrode and electroactive tissue, inducing current leakage. More flexible cuffs create a tighter electrical seal, leading to lower thresholds of input current for activation. However, to enable such flexibility, alternative material approaches, such as non-metallic conductors are needed to replace the metallic electrodes and tracks.^12^ Conductive elastomer (CE) electrodes, developed by Cuttaz et al.^13^ address the concerns of neural tissue damage, being significantly softer, organic and more conformal to soft tissues. Recent publications have demonstrated that CE polymer technologies can be laser micromachined into simple implantable nerve cuffs, enabling rapid development of bespoke designs for specific neuroanatomy.^13,14^ This current study adopts the same manufacturing technique to produce a transverse multipolar nerve cuff to investigate selective activation of underlying neural tissue.

While multielectrode arrays present the opportunity to steer current into nerves and selectively activate fibers from outside the epineurium,^15,16^ most studies in this field have been empirical, due to the difficulty of mapping the electrode placement to the underlying heterogeneous neuroanatomy.^17–19^ Therefore, the predictability of regions of activated fibers must be constructed by trial and error, which is a high risk and unreliable method for targeting stimulation of subpopulations of mixed nerves that innervate vital organs. An emerging approach that has assisted in understanding charge delivery and electrical field distribution within peripheral nerve interfaces is modelling and *in silico* experiments. The ASCENT (Automated Simulations to Characterize Electrical Nerve Thresholds) pipeline has recently been developed to enable a more accurate understanding of how electrodes placed on the peripheral nerves can be used to activate discrete areas of nerves or fiber types.^20^ This simulation environment is an important tool that assists in rapidly iterating both electrode design and stimulation parameters. While various studies have demonstrated validation of activation thresholds and conduction velocities using similar simulation pipelines, there has not been a highly controlled validation of fascicular activation within a simulation environment that incorporates the precise geometry of fascicles and overlying electrodes. A major limitation to modelling precise location of devices and neuroanatomy has been the lack of ability to image electrode arrays *in situ* and understand the relative location between stimulating electrode sites and the target neural tissue.

Typical approaches for mapping the nerve anatomy to the electrode placement use tissue stains, histological sectioning, and impedance tomography. All of these techniques can be time consuming and/or risk dislodging the electrode from the underlying nerve. Whole tissue imaging, such as magnetic resonance imaging (MRI), positron emission tomography (PET), and computed tomography (CT), where the electrode can be imaged in contact with the nerve has been challenging when used with metallic arrays, as the electrodes impart artefact due to interaction with the imaging modality. Recently, microCT with contrast enhancers has been used to discriminate anatomical constructs with clear delineation between nerve fascicles, sheaths and both fatty and connective tissues.^21^ When combined with CE electrode arrays, which absorb the contrast enhancer and have no associated artefact, direct mapping of the placement of an electrode cuff can be made to the underlying fascicular neuroanatomy. By pairing accurate electrode geometry and neuroanatomy with the ASCENT pipeline, a validation study of the ASCENT simulation environment for precise fascicular activation within a peripheral nerve is enabled.

This study details the design of a fully polymeric transverse multipolar electrode array and evaluates its performance in selective fascicular activation in a rat sciatic nerve. It then demonstrates the utility of microCT imaging to inform the computational model of accurate electrode-to-fascicle geometric relationships. Finally, it compares *in silico* simulated fiber activation to experimentally determined fascicular activations from an *ex vivo* rat sciatic nerve study, specifically evaluating the similarity in magnitude of fascicular selectivity indices and the stimulation parameters most responsible for selectivity.

## 2. Nerve Cuff Design and Manufacture

### Conductive Elastomer Nerve Cuff Design

The multipolar cuff designed for this study consisted of 8 rectangular electrodes (250 µm width, 2 mm length) circumferentially spaced when the cuff was wrapped around the nerve. The electrode pitch was 400 µm to fully cover the circumference of the approximately 1 mm diameter rat sciatic nerve, with specific geometries detailed in **Figure 1a**. The orientation of the cuff when placed on the rat sciatic nerve during the *ex vivo* setup is shown in **Figure 1b**. Nerve cuffs were manufactured with laser-based techniques, similar to those developed by Cuttaz et al.^14^ Briefly, a conductive elastomer (CE), being a dispersion of polystyrene sulfonate (PSS) doped poly(3,4-ethylenedioxythiophene) (PEDOT) in a polyurethane (PU) matrix was laser-cut to achieve electrodes, interconnects and bond pads. This was insulated with polydimethylsiloxane (PDMS) on both sides. The PMDS was laser ablated to achieve exposure of the active electrode sites. This final step was modified from the original fabrication protocol due to the size of the CE electrode active sites compared to previous literature. Where prior literature manually removed PDMS from relatively large electrodes, the smaller sizing of the array used herein required laser micromachining. A systematic study of the laser power and loops to achieve optimal exposure of the electroactive surface was undertaken (see Supplementary). Laser-ablated electrode performance was characterized and showed equivalence to those exposed with manual removal of the overlying PDMS.

**Figure 1.**
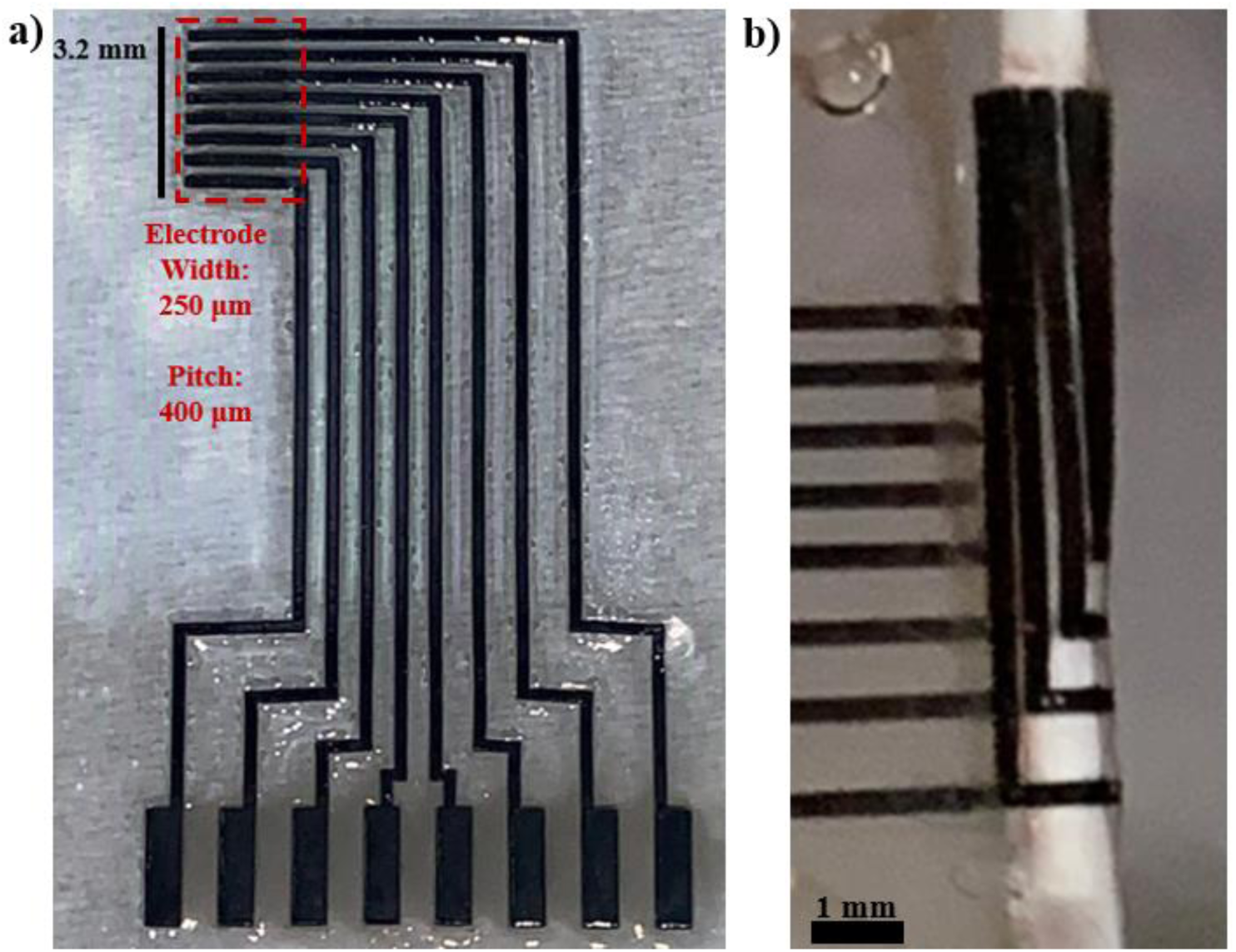
a) Multipolar nerve cuff design, 250 µm electrodes are spaced with a 400 µm pitch to cover the circumference of a 1mm rat sciatic nerve. b) Nerve cuff wrapped around an explanted rat sciatic nerve, image from one of the trials in this study.

### Electrode Array Electrochemical Performance

Electrochemical impedance spectroscopy (EIS) and cyclic voltammetry (CV) were used to assess device function and consistency in nerve cuff fabrication. The average impedance magnitude, phase shift, and voltametric response from all cuffs are shown in **Figure 2** and **Figure 3**. All 8 channels on all 5 cuffs were tested directly after manufacturing in saline and then while on the nerve during the *ex vivo* experiments, which are shown in the two curves (post-manufacturing: red and in *ex vivo*: blue). The average impedance for both conditions at 1 kHz (post-manufacturing: 10.27 ± 3.05 Ω•cm^2^, *ex vivo*: 16.61 ± 2.04 Ω•cm^2^) was within 1.41 SD, demonstrating consistent electrode performance across fabrication and testing stages. The shaded areas around each curve also indicate reliability across fabrication of each cuff. The impedance and phase measurements at 1 kHz are comparable to results from similar studies conducted by Cuttaz et al.^14^, demonstrating the scalability of laser microfabrication of CE electrode arrays. These studies also concur with prior literature across a broad range of PEDOT based electrode technologies,^22–25^ reflecting the notable lack of increase in phase shift at low frequencies which is inherent to the capacitive charge transfer of platinum electrodes. Furthermore, the electrode impedances remain low and stable when on the nerve during the *ex vivo* experiment. There is a slight increase compared to the post-manufacturing measurements, which aligns with the medium of the nerve in the chamber having a higher volume resistivity compared to the saline bath. Lastly, the average of CV hysteresis curves (**Figure 3**) across all cuffs shows consistency in shape and average charge storage capacity (CSC), with an average post-manufacturing CSC of 125.92 ± 21.80 mC/cm^2^ compared to an average *ex vivo* CSC of 104.24 ± 12.22 mC/cm^2^. These CSC measurements do not significantly differ from previously reported results^14^, and are ∼50 times larger than comparable CSC measurements from platinum electrodes. Similar to the impedance values, there is only a slight decrease in CSC values when comparing the *ex vivo* experiment to the post-manufacturing saline bath measurements, which can also be explained by the higher resistance across the surface of the electrode in the *ex vivo* nerve chamber.

**Figure 2.**
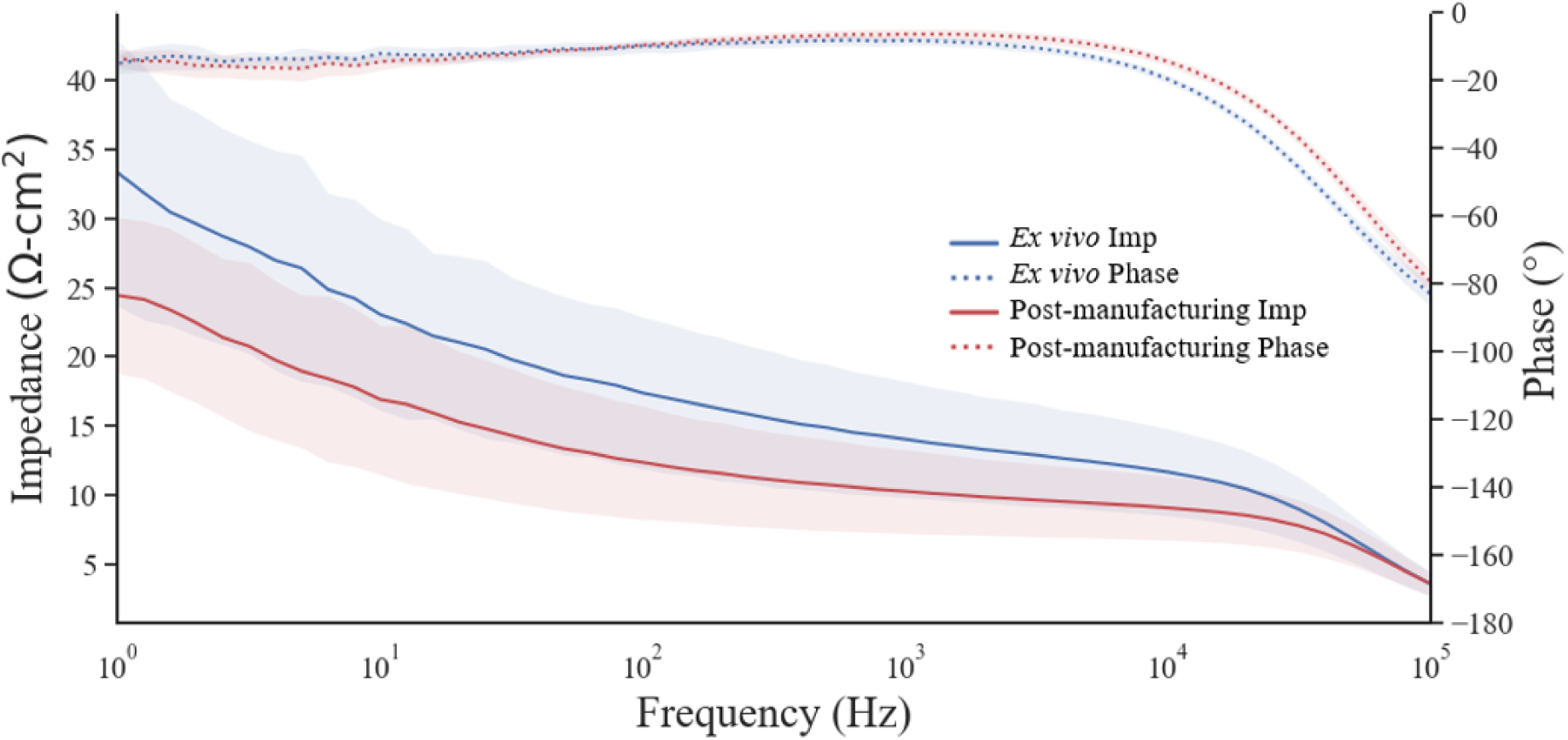
Average impedance magnitude and phase shift from an electrical impedance spectrogram (EIS) over all channels in all cuffs. Shaded areas represent 1 SD from the mean of each curve.

**Figure 3.**
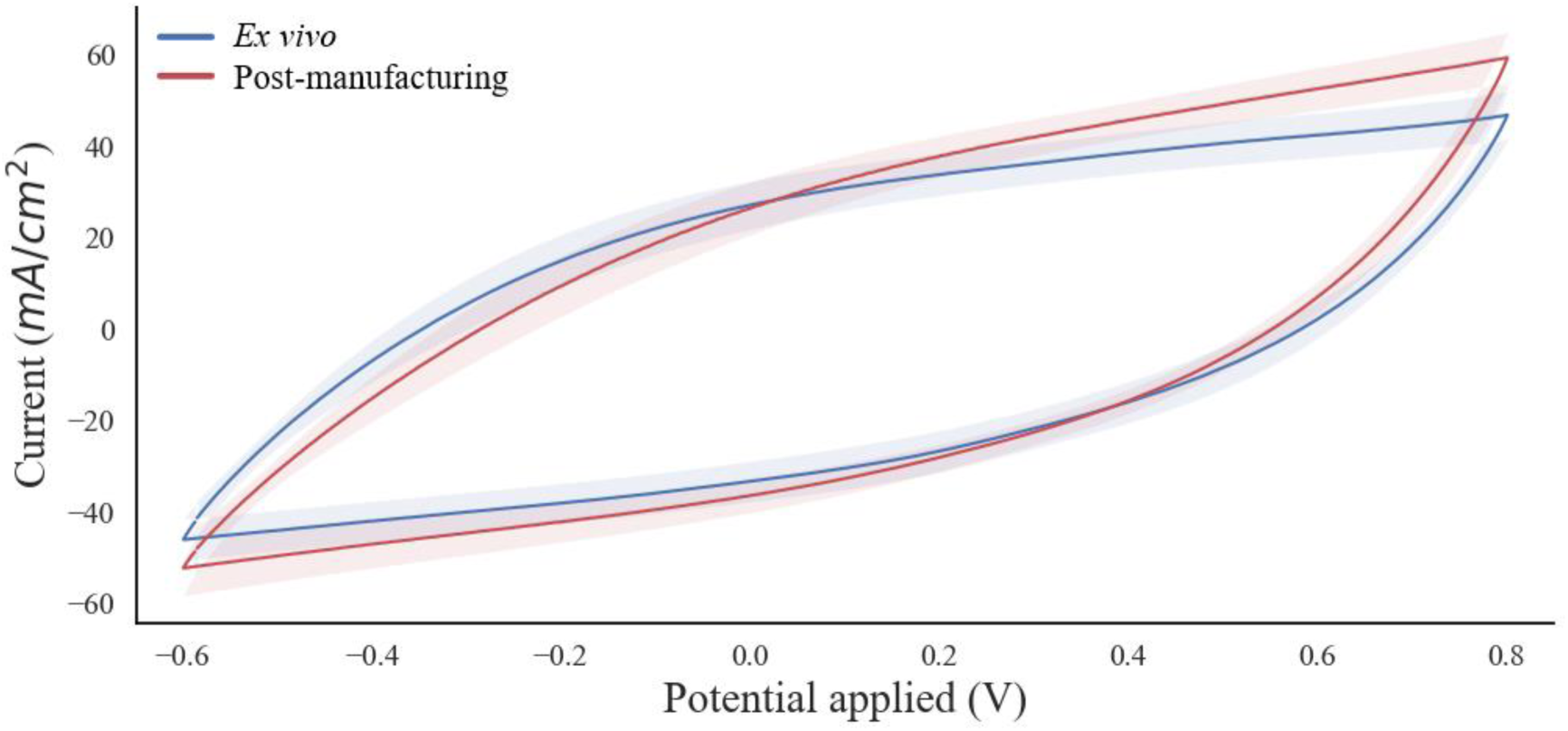
Average Cyclic Voltammetry over all channels in all cuffs. Shaded areas represent 1 SD from the mean of each curve.

## 3. Ex Vivo Experiments

### Experimental Setup

To assess the capabilities of spatially selective stimulation, nerve cuffs were wrapped around excised rat sciatic nerve and placed in a bath of oxygenated Krebs-Henseleit buffer solution. Stimulation of the nerve occurred on one side of the chamber within the buffer solution, while recordings were taken from each of the three trifurcated fascicles in a second chamber connected by a via sized for the rat sciatic nerve. The second chamber was filled with mineral oil to reduce noise and crosstalk between the bipolar recording hook electrodes on each fascicle. The nerve chamber can be seen in **Figure 4**.

**Figure 4.**
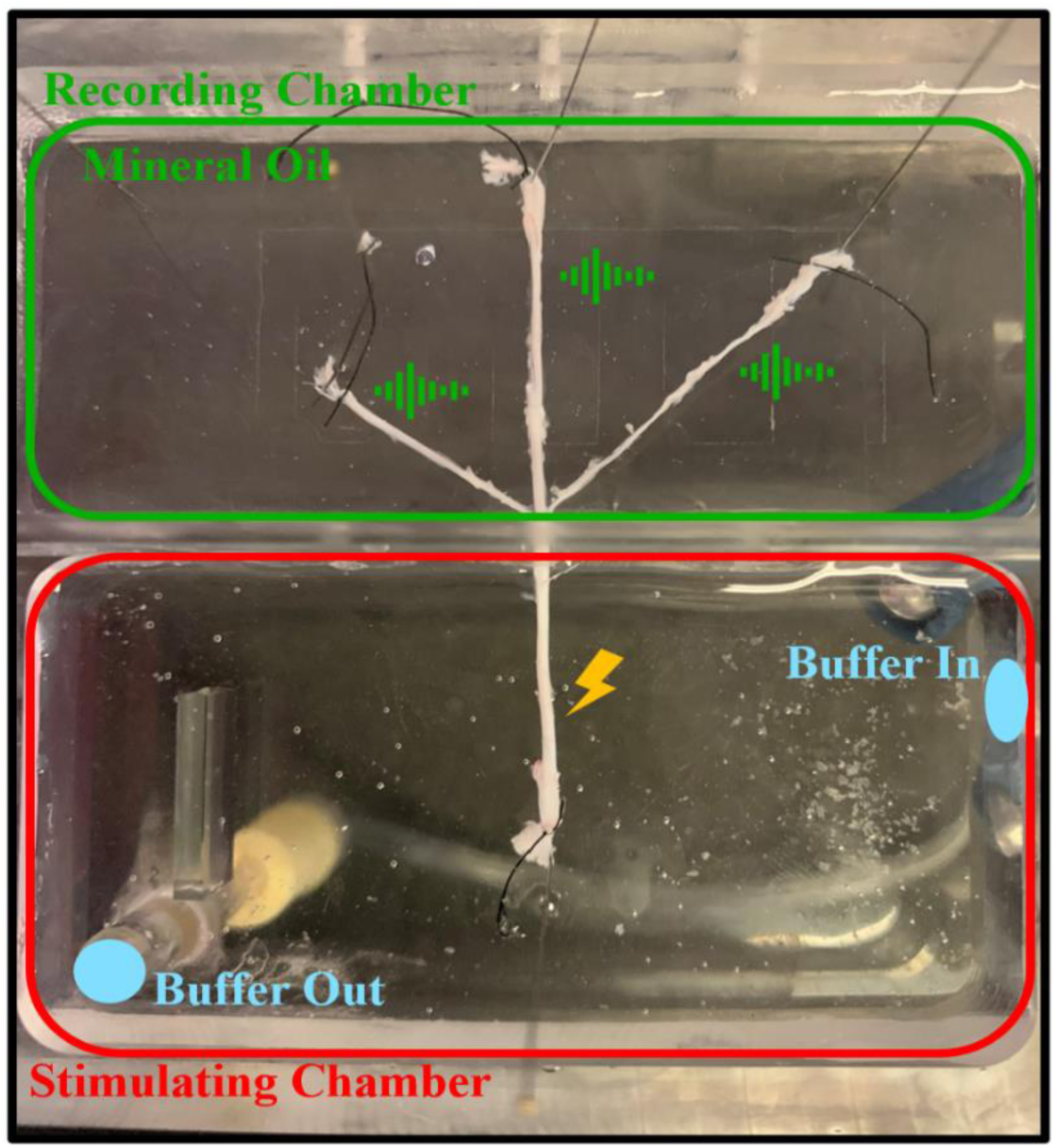
Excised and cleaned branching sciatic nerve placed within acrylic nerve chamber, in both the stimulating (red) and recording (green) chambers. The nerve proceeds through a feedthrough hole in the separating wall. Oxygenated Krebs-Henseleit buffer is cycled in and out of the stimulating chamber (blue). Nerve is stimulated (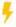) with a nerve cuff (not shown) and neural responses are recorded (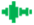) with 3 bipolar silver hook electrodes (not shown).

### Compound Nerve Action Potential (CNAP) Recordings by Fascicle

A systematic study spanning a broad range of stimulation protocols was investigated to ascertain the capacity for evoking fascicular level selectivity in the *ex vivo* sciatic nerve. **Figure 5** depicts the full range of the stimulation parameter space explored. Each unique combination of electrode configuration (bipolar or tripolar) and waveform shape (symmetric and asymmetric) were included in the study. The study was limited to this specific set of stimulation paradigms as the study was time-limited based on the viability of the nerve in the *ex vivo* chamber. The nearest neighbor electrode configurations were selected as they are known to isolate the current to a smaller portion of the nerve compared to a transverse stimulation pattern.^26^ Symmetric and asymmetric waveforms were used because of their common use in clinical settings. Most of the stimulation parameter space resulted in non-selective stimulation, either due to subthreshold stimulation for any fascicle, or suprathreshold for multiple fascicles. A non-selective response can be seen in **Figure 6a-b**, in which evoked CNAPs were measured in all three fascicles in response to the same stimulation. While the stimulation artefacts align in time, the neural responses were staggered based on the relative lengths of the fascicles that were harvested from the body, as the hook electrodes were attached 2 mm from the end of each fascicle. The selectivity indices (SI)^27^ for each fascicle are displayed below the voltage traces. These are calculated using **Equation 1**, which subtracts the average off target normalized activation of the other two fascicles from the normalized activation of the fascicle in question to obtain an SI for each unique set of stimulation parameters.

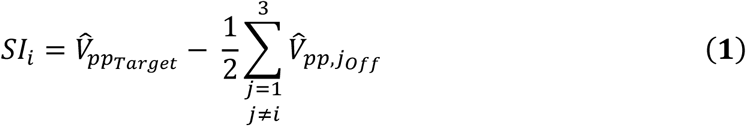

**Figure 5.**
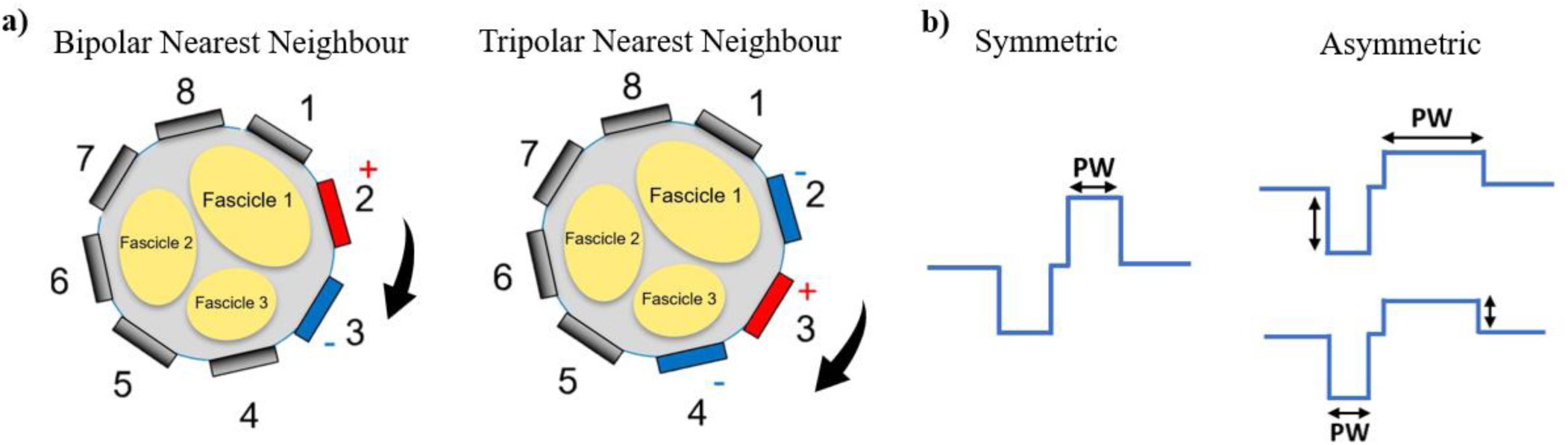
Stimulation parameter range for *ex vivo* experiments. a) Both bipolar and tripolar nearest neighbour (NN) electrodes were used for stimulation and the pairs or triplets were cycled around the nerve. b) Both symmetric and asymmetric biphasic waveforms were used in the bipolar and tripolar electrode configurations. For the asymmetric waveforms, in one instance the cathodic pulse was held at a constant pulse width while the anodic pulse was allowed to vary in pulse width, and in the other instance, vice versa. All biphasic waveforms were charge-balanced. For all unique combinations of electrode configuration and waveform, an amplitude sweep (0-1600 µA) and pulse width sweep (0-300 µs) was performed.

**Figure 6.**
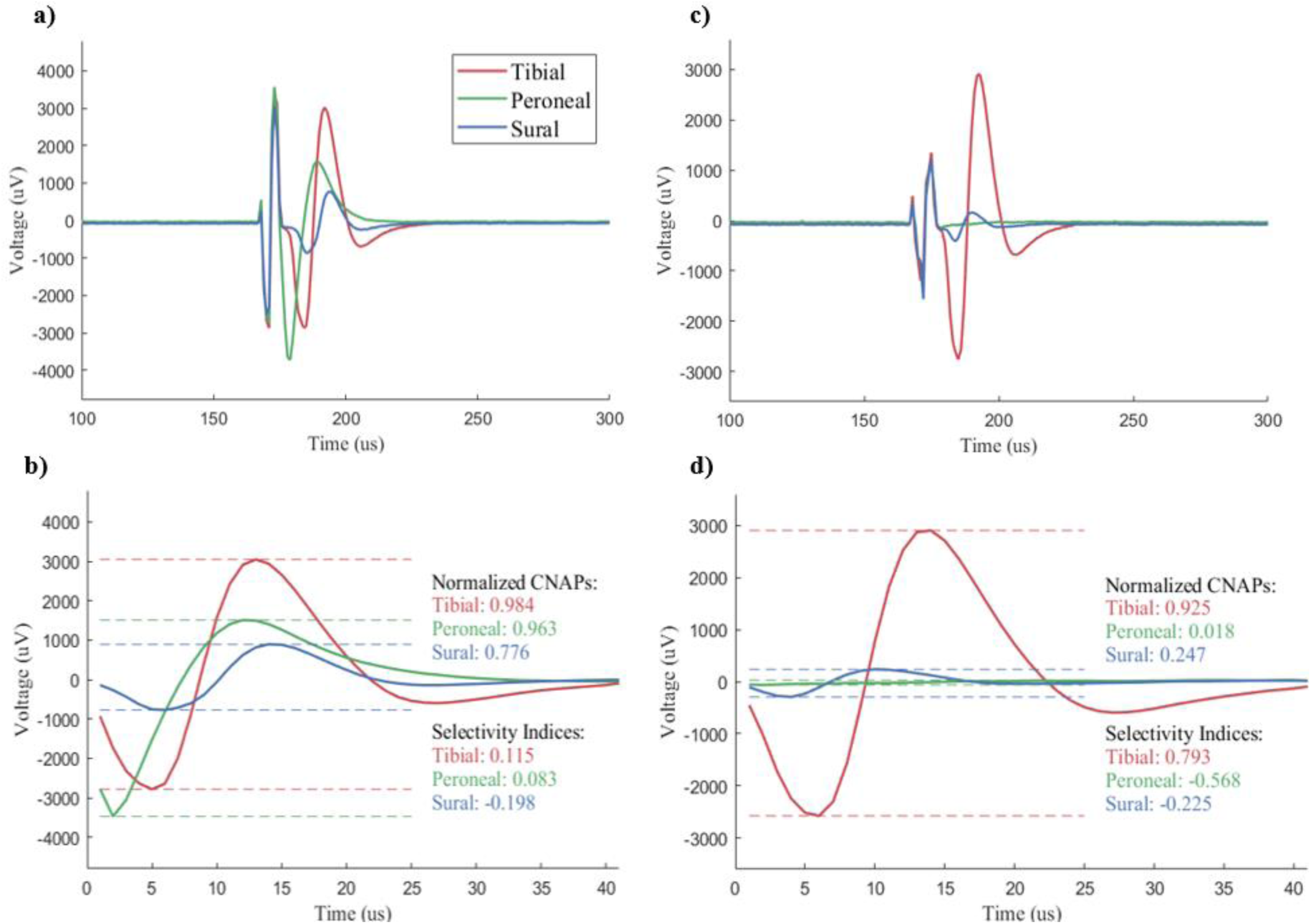
a) A voltage trace of non-selective neural recording of CNAPs for all three fascicles with the stimulation artefact included. b) Clipped voltage trace of non-selective neural recording of CNAPs for all three fascicles with the artefact removed. Selectivity indices near 0 because all fascicles are activated. c) A tibial-selective neural recording of CNAPs for all three fascicles with the stimulation artefact included. d) Clipped tibial-selective neural recording of CNAPs for all three fascicles with the artefact removed. Tibial selectivity is comparatively higher as the other two fascicles did not activate as much.

The non-selective response had SI values near zero or negative values, which indicate poor targeting of any fascicle. In opposition, the selective response (**Figure 6c-d**) was observed as an isolated neural response and that was corroborated by the high SI calculated. It is worth noting that the axis scales of each response have been kept the same to demonstrate that not only was the tibial nerve being selectively targeted in this example, but it was nearly at maximum recruitment as well. To best visualize how the fascicular selectivity changed with respect to the change in stimulation parameter space, a multiple contact heatmap was produced that represents the selective activation of each fascicle based on each unique stimulation parameter combination. The parameter space is defined by the active electrode and nearest neighbor (NN pair or triplet), the amplitude and pulse width. On each electrode NN pair/triplet a randomized pulse width sweep (0-300 µs) was performed, and a randomized amplitude sweep (0-1600 µA) was performed at each pulse width. **Figure 7** represents the symmetric waveform with bipolar stimulation for nerve trial #1. Heatmaps for all nerves and all configurations are available in the Supplementary Material. In this figure, “Electrode 1” represents the bipolar stimulation using electrode 1 as the source electrode and electrode 2 as the sink electrode. This convention was maintained as all NN pairs were utilized. For quick visualization of selectivity, each pixel in the heatmap was assigned a set of three values representing the fascicular activations (tibial, peroneal, sural). Each of those normalized activations was assigned the colors red, green, and blue, respectively, to create a color heatmap based on the RGB system. A black square represents [0, 0, 0] which means no activation and therefore no selectivity. A white square represents [1, 1, 1] which means high activation of all, so still no selectivity. Selectivity was visualized in the colored heatmap by a pure red, green, or blue color. While both a light red and a dark red pixel would represent only tibial activation, the dark red pixel would represent a higher value of selectivity as opacity of the color increases from 0 to 1. Therefore, there was high selectivity for the tibial fascicle in electrode 6, peroneal fascicle in electrodes 2, 3, and 5, and sural fascicle in electrode 7. Using the microCT image of the nerve in the middle of the heatmaps, it was easy to observe the positive relationship between the proximity of the fascicle to an electrode and the higher selectivity values for that particular fascicle when that electrode is being used as the source electrode. This is consistent with similar studies using metallic multipolar cuff arrays.^16,28–30^ These observations include the inversely proportional relationship between pulse width and amplitude that is present in a strength-duration curve commonly used in peripheral nerve stimulation literature.^31–34^ This trend aligns with the known understanding of evoked action potentials depending on charge injection (the product of amplitude and pulse width) and not directly on either individual parameter.^11^ The minimum amount of current required for activation, the rheobase, can also be observed across all electrodes. It typically lies around 400 uA at a pulse width of 100 uA for these electrodes. At 40 nC of injected current, this is in the general range of threshold charge injection for similar nerve cuff experiments^35,36^ and within the range of safety limitations for PEDOT-based electrodes.^37^

**Figure 7.**
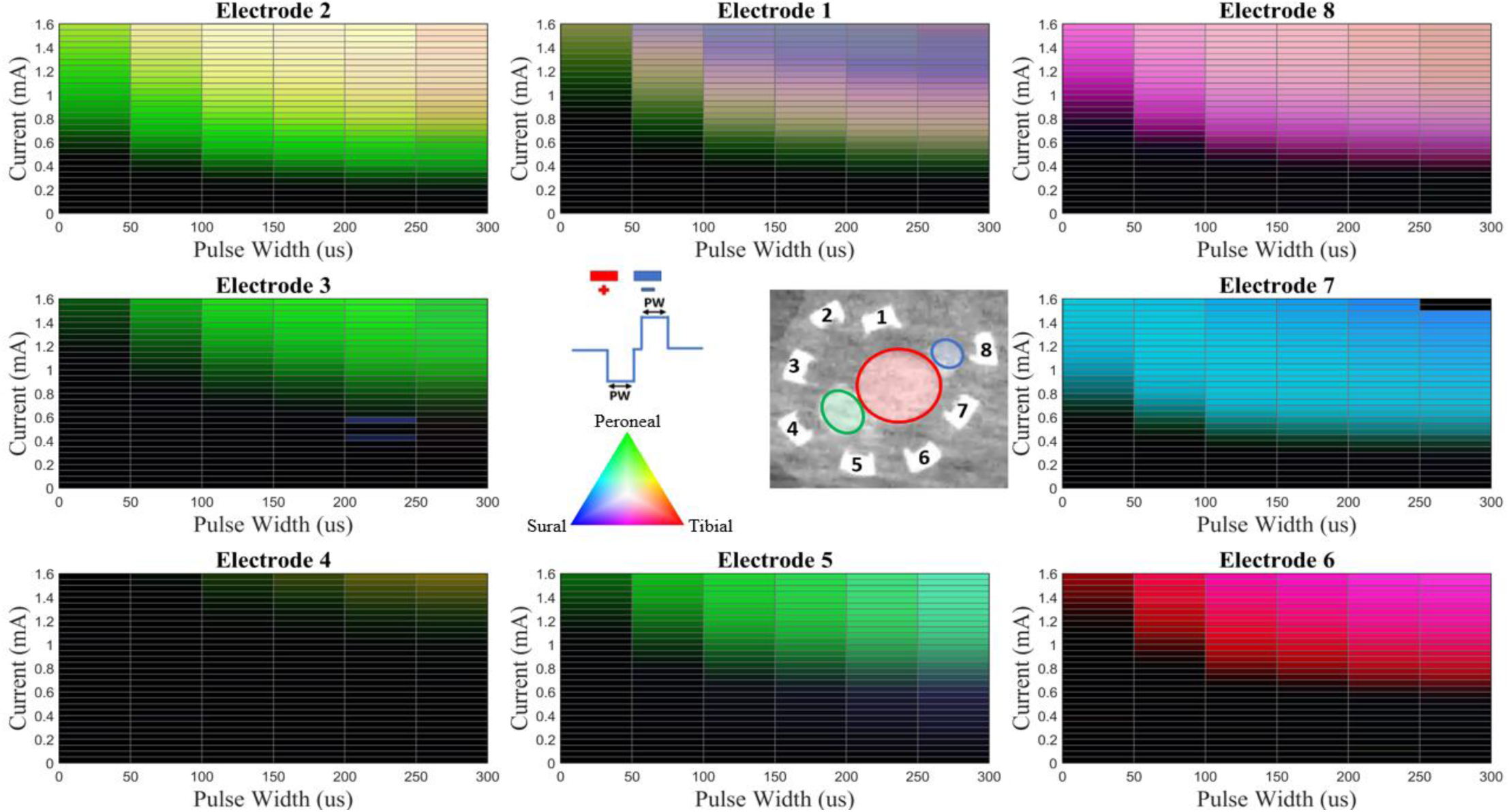
Multi-contact selectivity heatmaps of *ex vivo* stimulation. Each contact represents the stimulation delivered using that contact as the source and the anti-clockwise nearest neighbor as the sink. Tibial selectivity is represented by red, peroneal by green, and sural by blue. The microCT image in the middle shows the relationship between the overlying electrodes and the underlying fascicles, which are color-coded appropriately. These selectivity results are only from one nerve (#2) and one waveform configuration (bipolar symmetric). The color triangle indicates the activation color for each fascicle and the blended colors of multiple fascicular activations.

### Ex Vivo Selectivity Trends

To evaluate trends across nerve trials and waveform configurations, the top 5 values of selectivity (with a lower threshold of SI > 0.25) were selected from each of the 5 unique combinations of electrode configurations and waveform types. This was done for each fascicle in each nerve trial and then averaged across nerve trials (**Figure 8**). Across the nerve trials, each fascicle was selectively activated equally by each of the waveforms. While configuration 4 (tripolar) first appears to provide higher sural selectivity, a repeated-measures ANOVA test reveals the lack of a significant difference between any of the configurations for a particular fascicular selectivity (p = 0.2298). Post-hoc paired tests revealed an uncorrected significant difference between configuration 1 and 4 for the sural fascicle specifically (p = 0.0448), but after accounting for the other configurations, there was not enough evidence to suggest a fascicular preference for a particular waveform or number of poles. Perhaps with more repeats, a relationship would become more pronounced, as the tripolar, asymmetric waveform profile is known to localize the current distribution to the highest degree of any of these configurations.^15^ While it is safe to say that on average, each fascicle can be selectively targeted at an SI of at least 0.6, agnostic of the underlying neuroanatomy, the large standard deviations in **Figure 8** indicate that there are notable differences in the selectivity values achieved across different nerve trials. To understand the influence on the neuroanatomic diversity on the selectivity outcomes, the nerves were imaged with the nerve cuffs in place. MicroCT has been used to image and map the neuroanatomy of fascicles in peripheral nerves that have been stained with a contrast enhancer.^21^ This study is the first to incorporate a fully intact multicontact cuff electrode array, enabled by conductive polymer material in the array that does not produce metallic artefacts in microCT scans. Following the *ex vivo* stimulation and recording sessions for each nerve trial, the nerve and cuff were immediately embedded in alginate set with a 6% Lugol’s iodine solution. The alginate block was immersed in Lugol’s solution (containing iodine) to provide a differential tissue contrast stain and subsequently scanned using a microCT. Because the conductive elastomer also absorbs the iodine contrast stain, the nerve cuff electrodes are easily visualised, without causing artefacts such as those typically associated with metals in peripheral nerve interfaces. The anatomical variability across nerve trials can be explicitly seen in **Figure 9**, in which the three primary fascicles of the sciatic nerve have been highlighted in each trial (red: tibial, green: peroneal, blue: sural). To more directly observe the differences in selectivity due to anatomical and electrode placement differences, the top selectivity for each configuration within a nerve has been averaged together, and nerve trials are then directly compared (**Figure 10**). When the three fascicular selectivity indices are then averaged together for each nerve and then compared across trials, there is not significant difference between the averaged fascicular selectivity indices across nerve trials. However, a repeated measures ANOVA test when keeping fascicle selectivity indices separate reveal a significant difference in the selectivity values of the tibial fascicle across nerve trials. Post-hoc paired tests confirmed that both trials 1 and 4 significantly differ from nerve trial 3 (p<0.01) and from nerve trial 5 (p<0.05). The significantly lower selectivity in nerve trial 4 in the tibial fascicle may be explained by the positioning of the tibial fascicle, in between the two electrodes that were relatively far apart. The microCT also reveals that in this particular trial the cuff did not fully close around the nerve. This positioning of the electrodes relative to the fascicle occurred as a function of random neuroanatomy and uninformed surgical placement of the cuffs on nerves. Although fascicles are sometimes visible through the epineurium, in this study, no specific orientation of the cuff closure to particular fascicles were chosen a priori.

**Figure 8.**
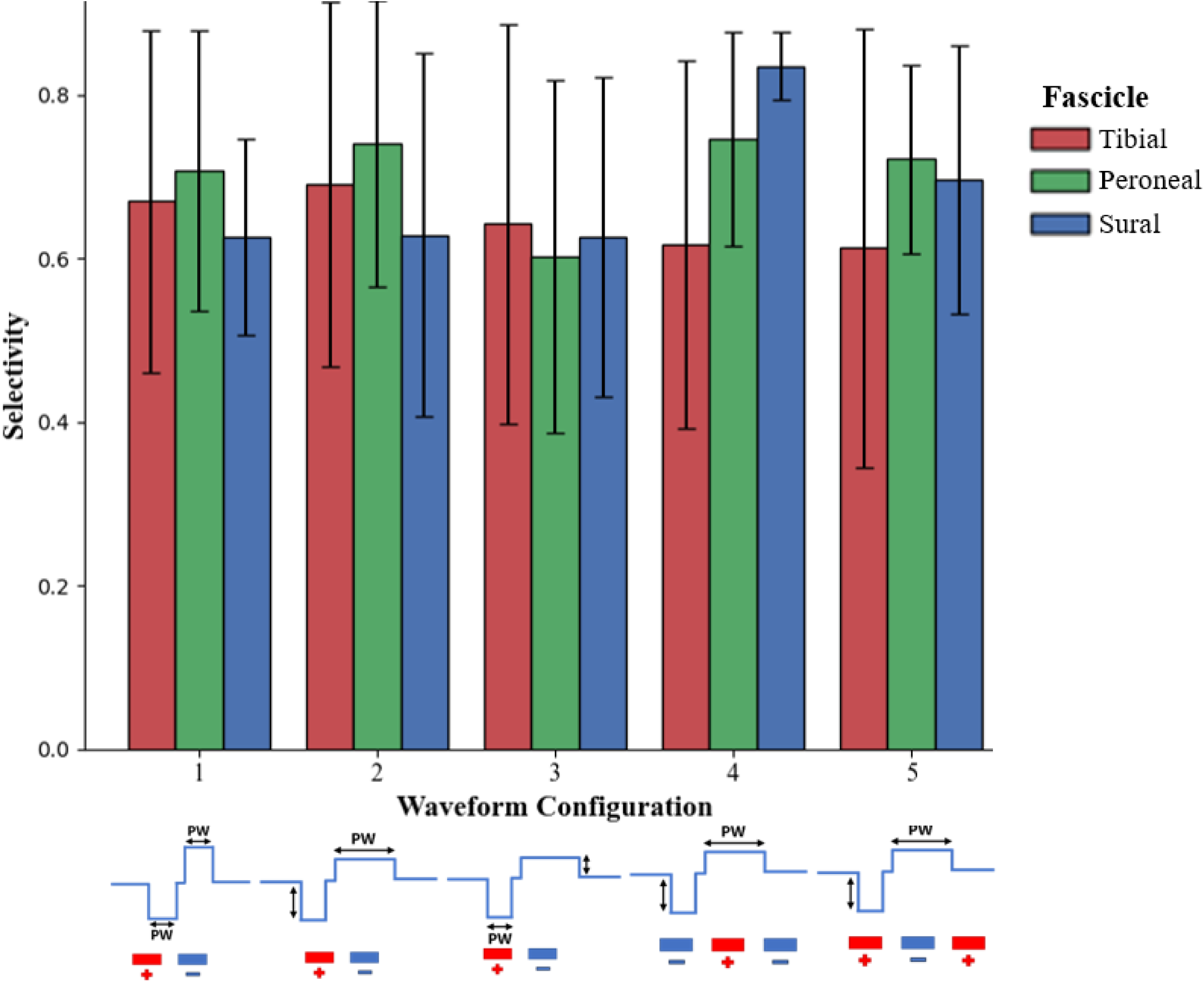
Top 5 selectivity indices in each of these stimulation configurations from each fascicle on each nerve. Error bars represent 1 SD from the mean of the 5 animals selectivity indices in each configuration.

**Figure 9.**
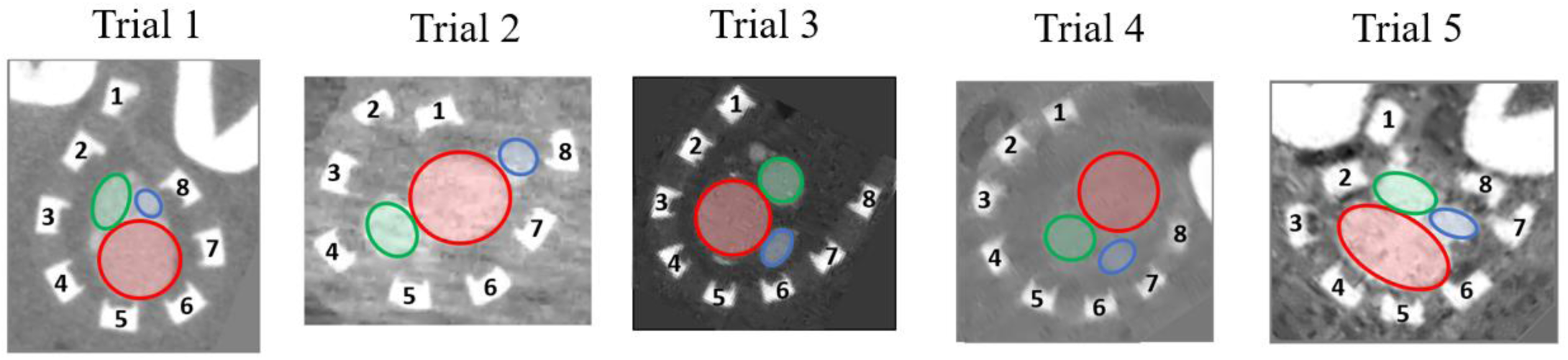
MicroCT images of the *ex vivo* nerve trials captured 48 hours after the stimulation session after the tissue had been fixated and stained in alginate set with 6% Lugol’s solution.

**Figure 10.**
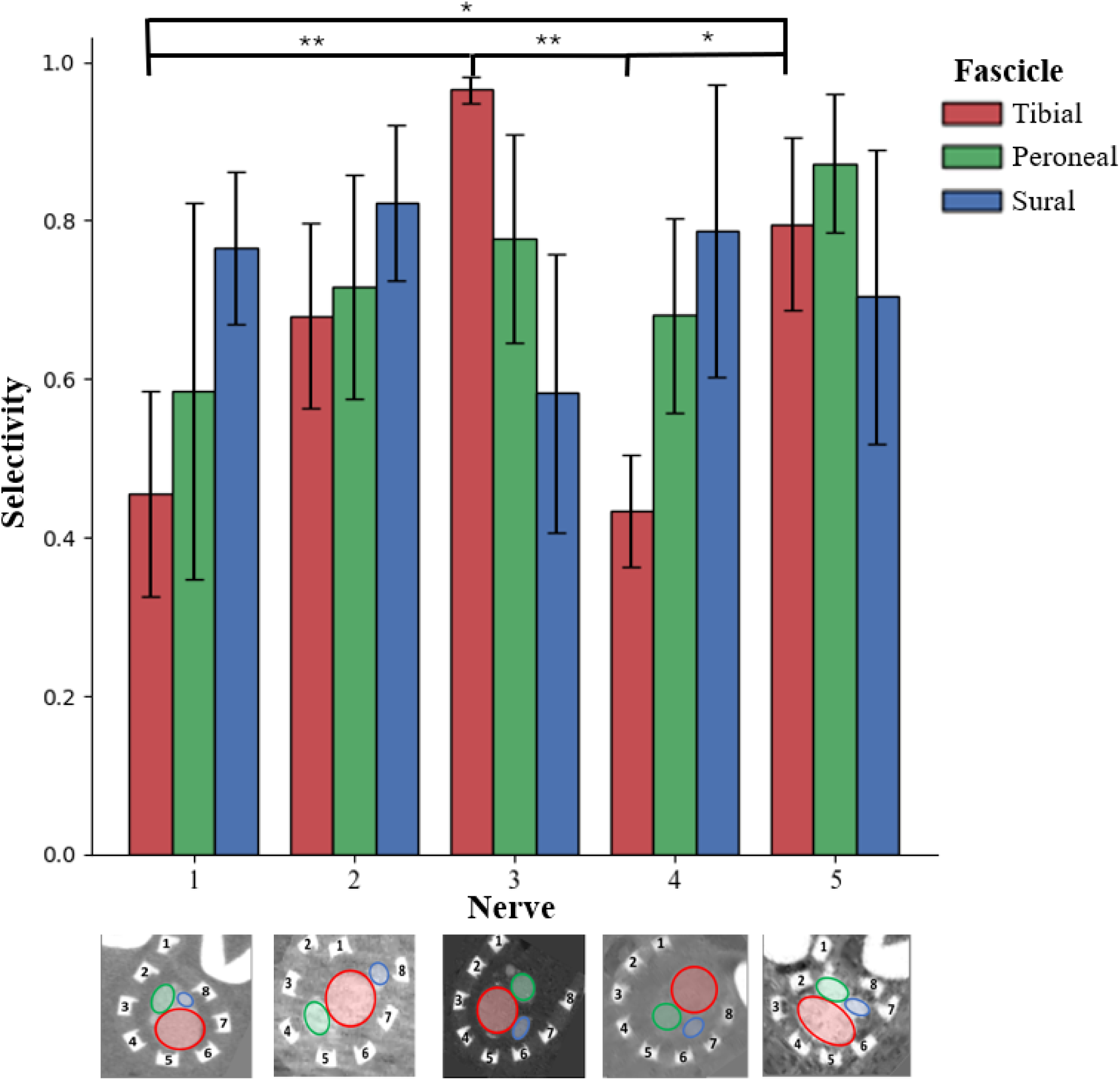
Top selectivity per configuration in each *ex vivo* nerve averaged together. The nerves with color-coded fascicles are underlying the x-axis. Error bars represent 1 SD. * (p < 0.05), ** (p < 0.01).

## 4. Simulation

### Creation of Model Geometry from MicroCT

Referencing the microCT images seen in **Figure 9**, the electrode-to-fascicle geometric relationships were captured and recreated within the ASCENT pipeline to model *ex vivo* setup most accurately. A custom elliptical part primitive was designed to appropriately generate the shape of the PDMS insulation around each nerve. **Figure 11b** shows the computation models rendered in COMSOL (v5.6) for each of the nerve trials.

**Figure 11.**
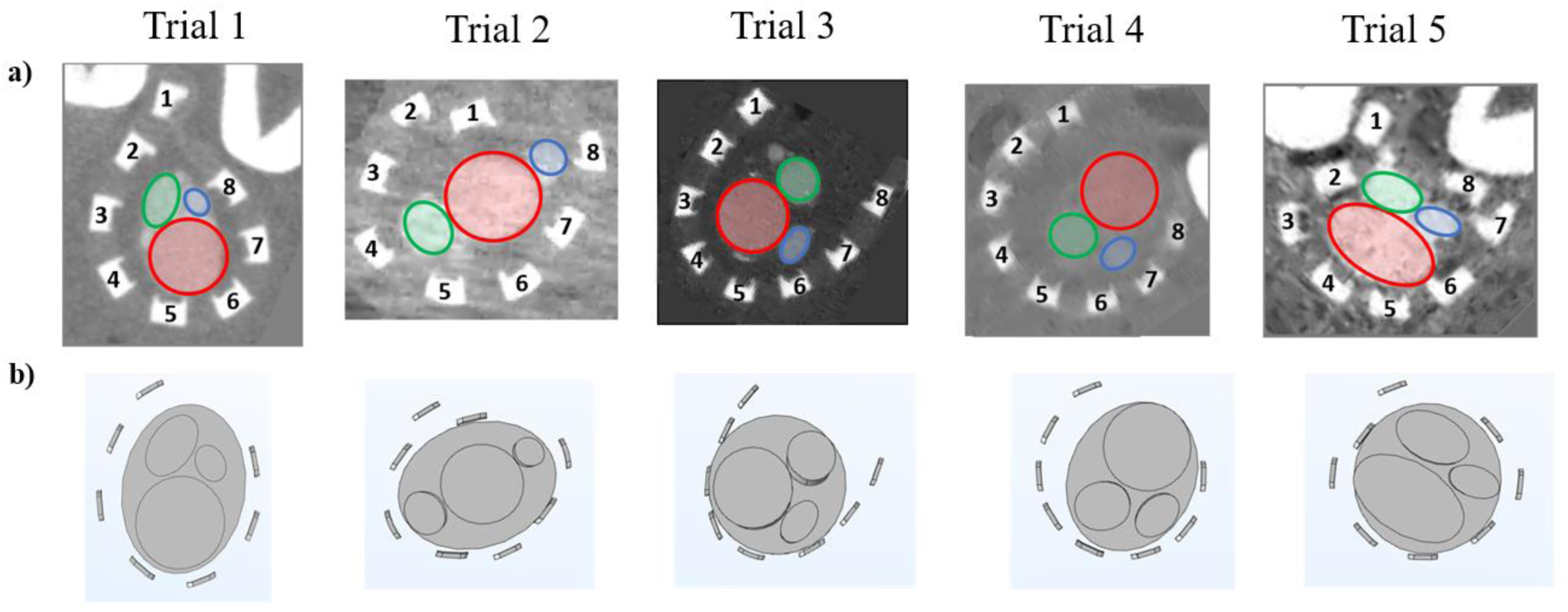
a) MicroCT images of the *ex vivo* nerve trials captured 48 hours after the stimulation session after the tissue had been fixated and stained in alginate set with 6% Lugol’s solution. b) COMSOL generated meshes of the reconstructions of each of the nerve trials within the ASCENT pipeline.

### Comparison of Simulated and Ex Vivo Selective Stimulation

Once the nerve models were prepared and all model cuffs were made, the same stimulation parameters were applied to the *in silico* models within the ASCENT pipeline. To measure the magnitude of fascicular activation, the number of fibers activated in a particular fascicular was divided by the total number of fibers in that fascicle to give a normalized activation value. This value was then used to calculate selectivity using the same equation as the *ex vivo* selectivity calculations **(Equation 1)**. Heatmaps similar to **Figure 7** were created for all nerves and waveform configurations, mapping the theoretical selectivity for each simulated stimulation. The simulated selectivity of same nerve (Trial 2) and waveform configuration (even, bipolar) from **Figure 7** is shown in **Figure 12**. While there are clear similarities between the specific electrodes that evoke selective activation in specific fascicles, there are few key differences: the large drop in activation thresholds in the simulation compared to the *ex vivo*, the overprediction of the sural fascicle activation (blue), and the lack of an isolated tibial activation (red). The selectivity results between the *ex vivo* and the simulation were investigated across all nerves and waveform configurations, comparing the magnitudes of selectivity and the electrode positions that evoked the highest selectivity.

**Figure 12.**
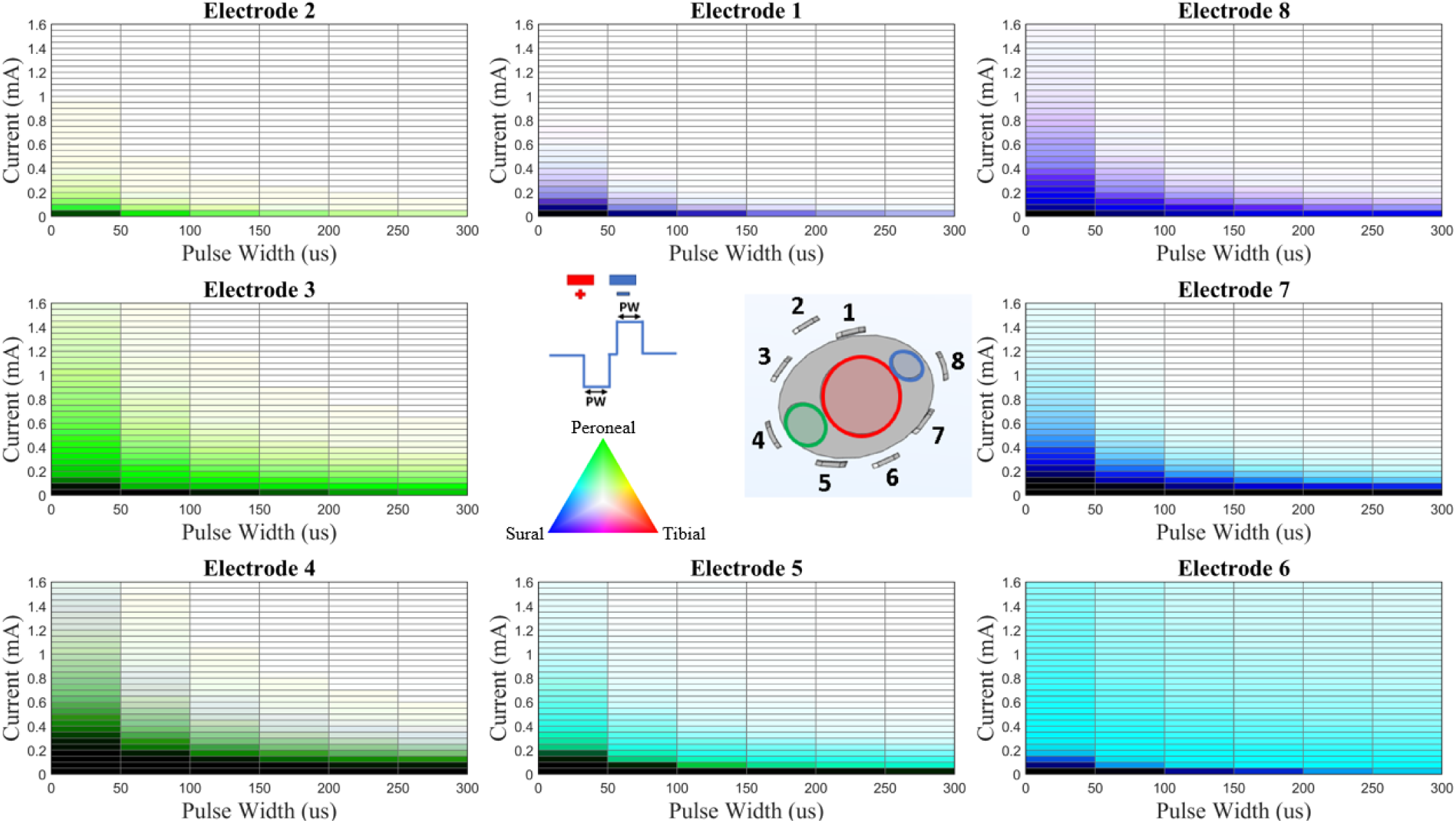
Multi-electrode selectivity heatmaps of simulated stimulation. Each electrode represents the stimulation delivered using that electrode as the source and the anti-clockwise nearest neighbor as the sink. Tibial selectivity is represented by red, peroneal by green, and sural by blue. The COMSOL image in the middle shows the relationship between the overlying electrodes and the underlying fascicles, which are color-coded appropriately. These selectivity results are only from one nerve (#2) and one waveform configuration (bipolar symmetric). The color triangle indicates the activation color for each fascicle and the blended colors of multiple fascicular activations.

### Selectivity Magnitude Comparison

When the top 5 selectivity indices from each nerve were averaged together for each waveform configuration (**Figure 13**), the overprediction of the sural selectivity seen in **Figure 12** became apparent across all nerves (low standard deviation) and all configurations. The only significant differences in average selectivity indices when isolating the fascicles was between the *ex vivo* and simulated sural fascicle selectivity (p<0.03 for all). While not significantly different across the nerves for any configuration, there was a trend of lower predicted selectivity of the tibial fascicle across most of the configurations. This limited tibial selectivity can be observed in **Figure 12** as well with the lack of red pixels in the expected electrode heatmaps. However, the lowest threshold tibial activation is still observed between electrodes 8 and 1, where the purple color in the pixel is a combination of the tibial (red) and sural (blue) activations. While this is not the same electrode location as the highest selectivity for the tibial fascicle in the *ex vivo* experiments (electrode 6), electrode 1 is still in closer proximity to the tibial fascicle than most other electrodes, due to the unique alignment of the fascicles in trial 2. Finally, similar to the *ex vivo* data, the standard deviations of the tibial and peroneal selectivity indices are very large, indicating a large effect of the neuroanatomical variability on the selectivity achievable in each nerve. In fact, the simulation selectivity has larger standard deviations for the peroneal (+120%) and tibial (+35%) fascicles, which indicate the simulation selectivity results are more sensitive to differences in neuroanatomy than the *ex vivo* results. Only the sural fascicle had a low standard deviation in the simulation data, but primarily because it achieved nearly 100% selectivity in all configurations, agnostic of neuroanatomy.

**Figure 13.**
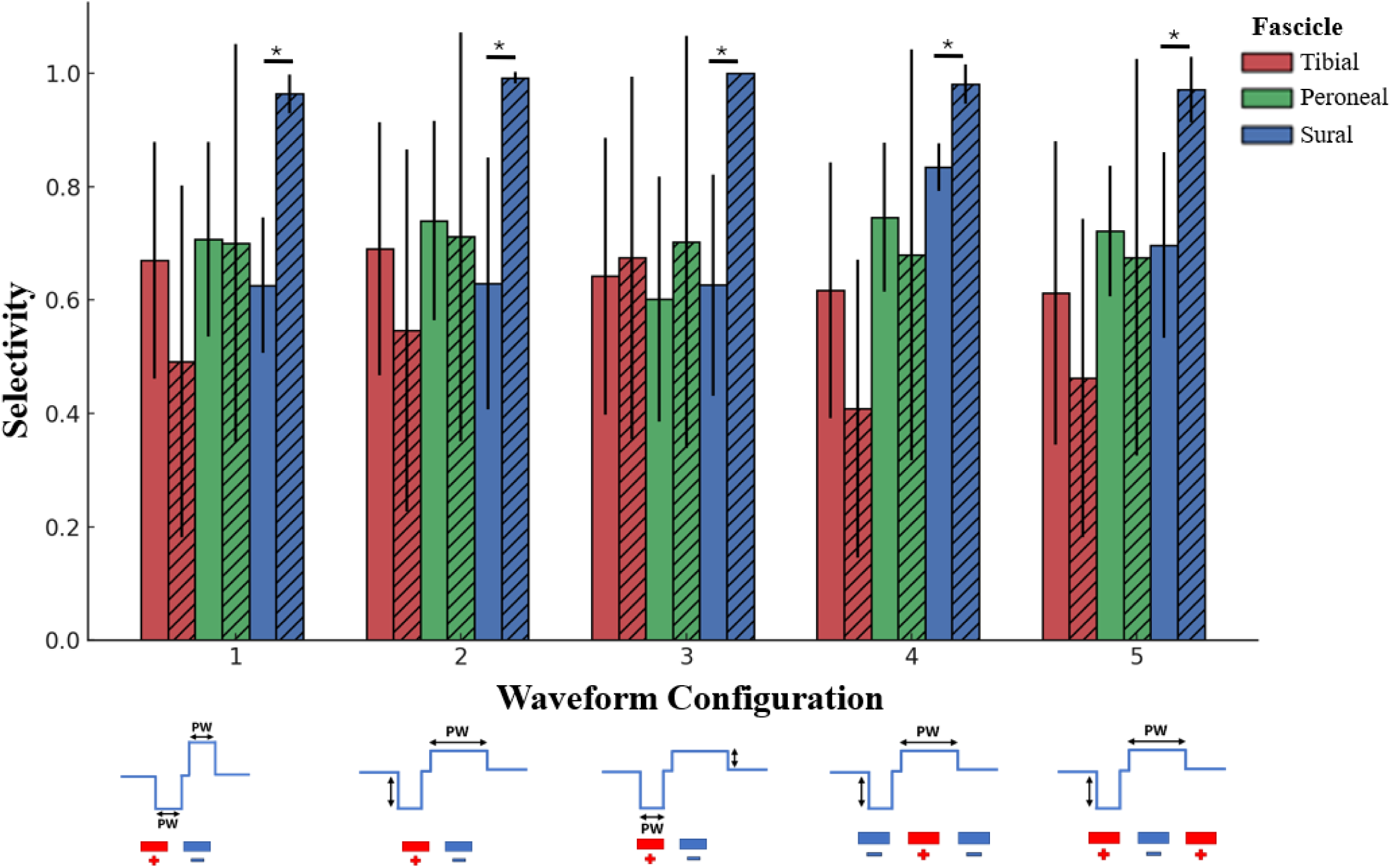
Comparison of *ex vivo* (solid) and simulation (striped) selectivity results. Top 5 selectivity indices in each of these stimulation configurations from each fascicle on each nerve. Error bars represent 1 SD from the mean of the 5 animals selectivity indices in each configuration. * (p < 0.05)

To investigate this neuroanatomic dependence further, the top selectivity from each configuration was averaged for each nerve trial and compared in **Figure 14**. The overprediction of the sural nerve was still present in every nerve trial, each with a statistically significant difference (p<0.05). There was much more variation in average selectivity for the tibial and peroneal fascicles across different nerves. The selectivity indices of the peroneal fascicle were generally quite high and similar between the *ex vivo* and simulated results, except for trial 3. The stark difference in the trial 3 peroneal fascicle was partially explained by the location of the fascicles in relation to the overlying electrodes. The peroneal fascicle in trial 3 was on the most exposed side of the nerve bundle where the nerve cuff did not fully wrap around and conform to the nerve bundle. While this non-ideal cuff fit led the simulation to predict very little selective activation, the *ex vivo* experiments resulted in one of the highest average selectivity indices for the peroneal fascicle, indicating again that the model is more sensitive to non-ideal nerve-cuff geometries. The lowest simulated tibial fascicle selectivity was trial 2, where the tibial fascicle had a unique relationship to the other two fascicles. Rather than having a triangular shape with one fascicle at each vertex, the trial 2 nerve had a linear arrangement of the fascicles along the midline of the nerve. This arrangement limited the direct exposure of the tibial fascicle to isolated electrodes. While trials 1 and 4 match quite well in their fascicular trends, the other 3 trials differ significantly in their tibial and/or peroneal predicted and experimental data for selectivity magnitude, even without accounting for the similarity between the location of the electrodes that provided those highest selectivity indices.

**Figure 14.**
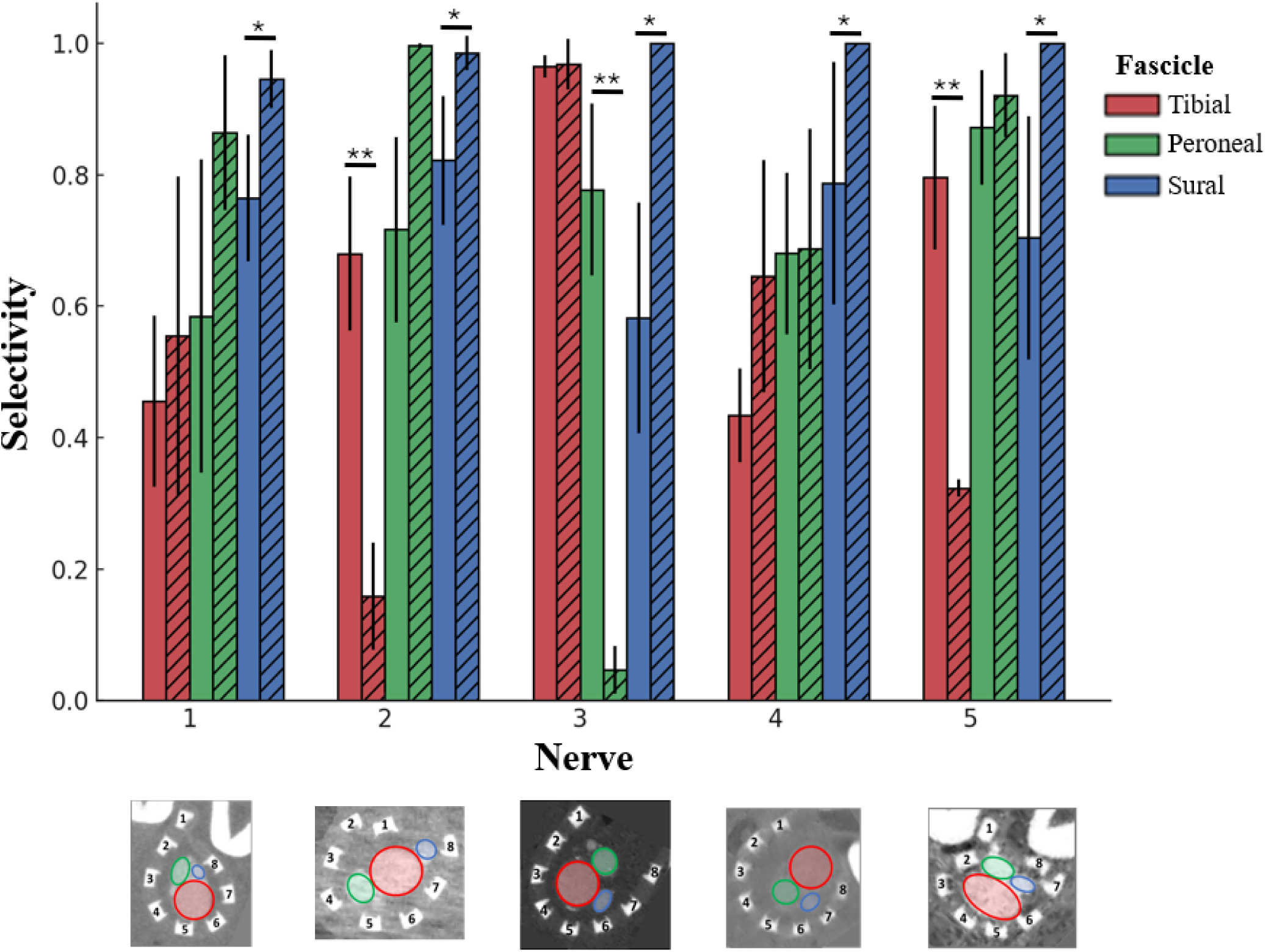
Comparison of *ex vivo* (solid) and simulation (striped) selectivity results. Top selectivity indices for each stimulation configuration were averaged together across the same nerve. Error bars represent 1 SD from the mean of the 5 configuration selectivity indices in each configuration. * (p < 0.05)

### Electrode Agreement Metric

The difference between predicted and actual selectivity based on electrode position was quantified and nominally called the electrode agreement (EA) metric. This was achieved by specifically comparing the highest selectivity values based on electrode position for each nerve trial. The EA metric was defined by the normalized radial distance between the average position of the electrodes that provided the top 10 selectivity indices for the *ex vivo* results and the average position of those respective electrodes for the simulations of the same trial. The radial distance was calculated on an idealized circle with evenly spaced electrodes, not the actual spacing measured from the CT scans. The metric was used to evaluate the identity of the most selective electrodes, comparing simulation to experimental results, not to try to understand the relationship between electrode position and fascicular location. **Figure 15** shows the EA plot for Trial 2 with the electrode agreement (EA) metrics for each fascicle shown color coded in the bottom right. This polar plot was separated into 8 sectors, each representing an electrode on the nerve cuff. Each data point was plotted into the sector representing the electrode responsible for that top 10 selectivity index. The distance away from the origin was defined by the amplitude at which that selectivity index was produced. The subset angle within the sector of each data point was defined the pulse width at which that selectivity index was produced. The opacity of the electrodes data point was defined by the magnitude of the selectivity index (0-1). Finally, the *ex vivo* data was represented by dots, while the simulation data was represented by crosses. The EA metrics for each fascicle in Trial 2 were 0.75 (tibial), 0.5 (peroneal), and 0.75 (sural). An EA metric of 0 indicated that the electrodes were on the direct opposite side of the polar plot and most likely the nerve as well, while a 1 indicated that the electrodes are the same electrode. Therefore, a 0.75 indicated that the predicted and *ex vivo* electrodes that provided the top 10 selectivity indices for a particular nerve and configuration were just 1 electrode apart, or they are nearest neighbors. An EA metric 0.5 indicated two electrodes apart, and so on. To investigate this metric across all nerve trials and configurations, the top 10 selectivies for each configuration within a nerve were averaged together, shown in **Figure 16**. There were not any significant trends across nerves for particular fascicles having a higher electrode agreement score. The peroneal and sural fascicles had an average above 0.50 across all nerves, therefore, the model predicted the most selective electrode for the *ex vivo* within at least 2 electrode positions for every peroneal and sural highly selective activation. With the precision of 2 electrodes on a 8 electrode cuff, the most accurate model predictions for electrode positions that evoke selectivity is on the same half of the nerve. However, this does not discount that higher prediction and selectivity may be obtained from cuffs with higher numbers of electrodes, yielding improved resolution.

**Figure 15.**
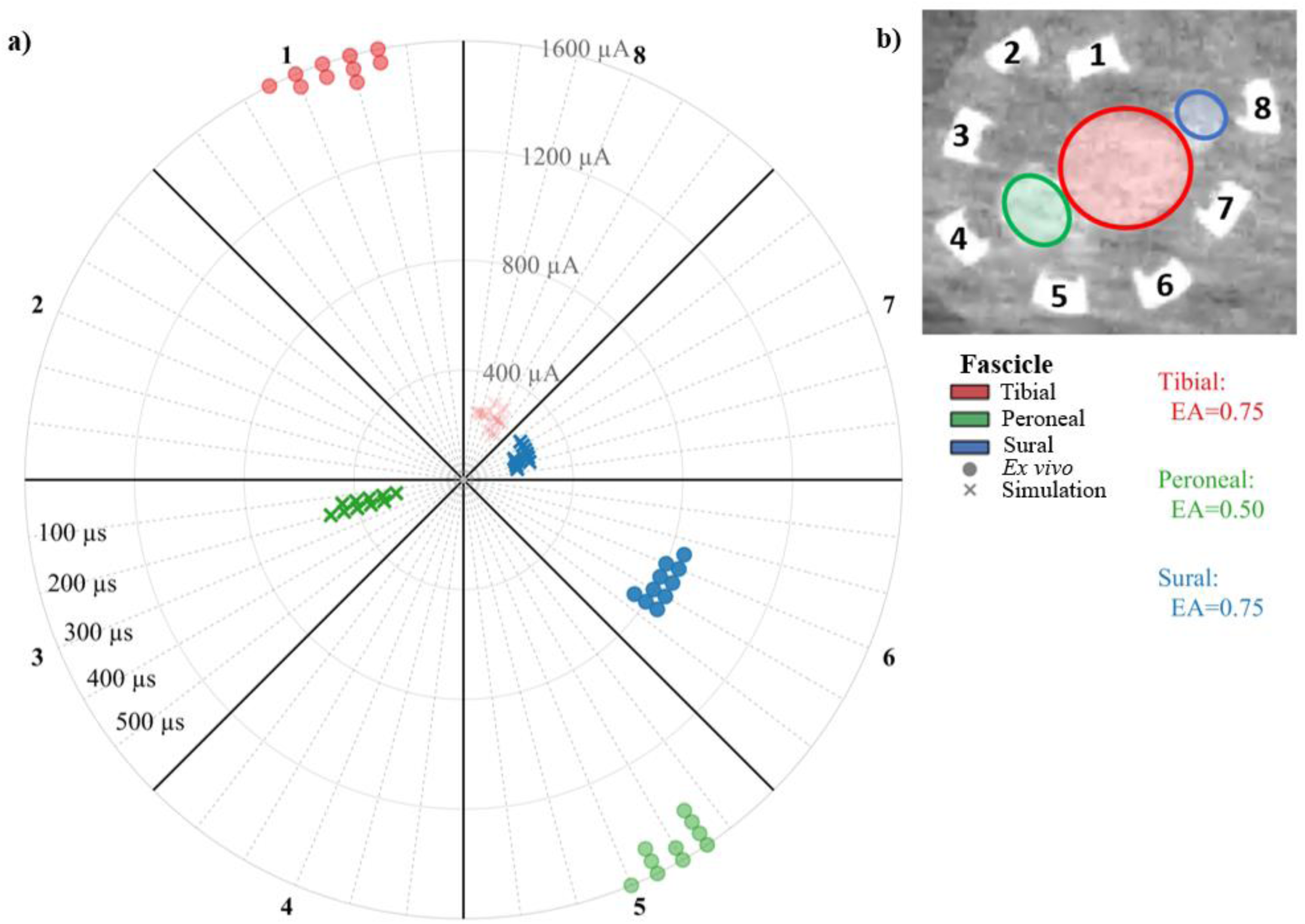
Electrode agreement metric diagram. a) Selectivity data plotted onto polar plot divided into 8 sectors to represent the 8 electrodes, not exact locations, just similar visual representation. The rings are levels of current amplitude for each data point. The radii are levels of pulse widths increasing anticlockwise. Ex vivo data represented by circles. Simulated data represented by crosses. b) Trial 2 nerve that corresponds to the data in the polar plot.

**Figure 16.**
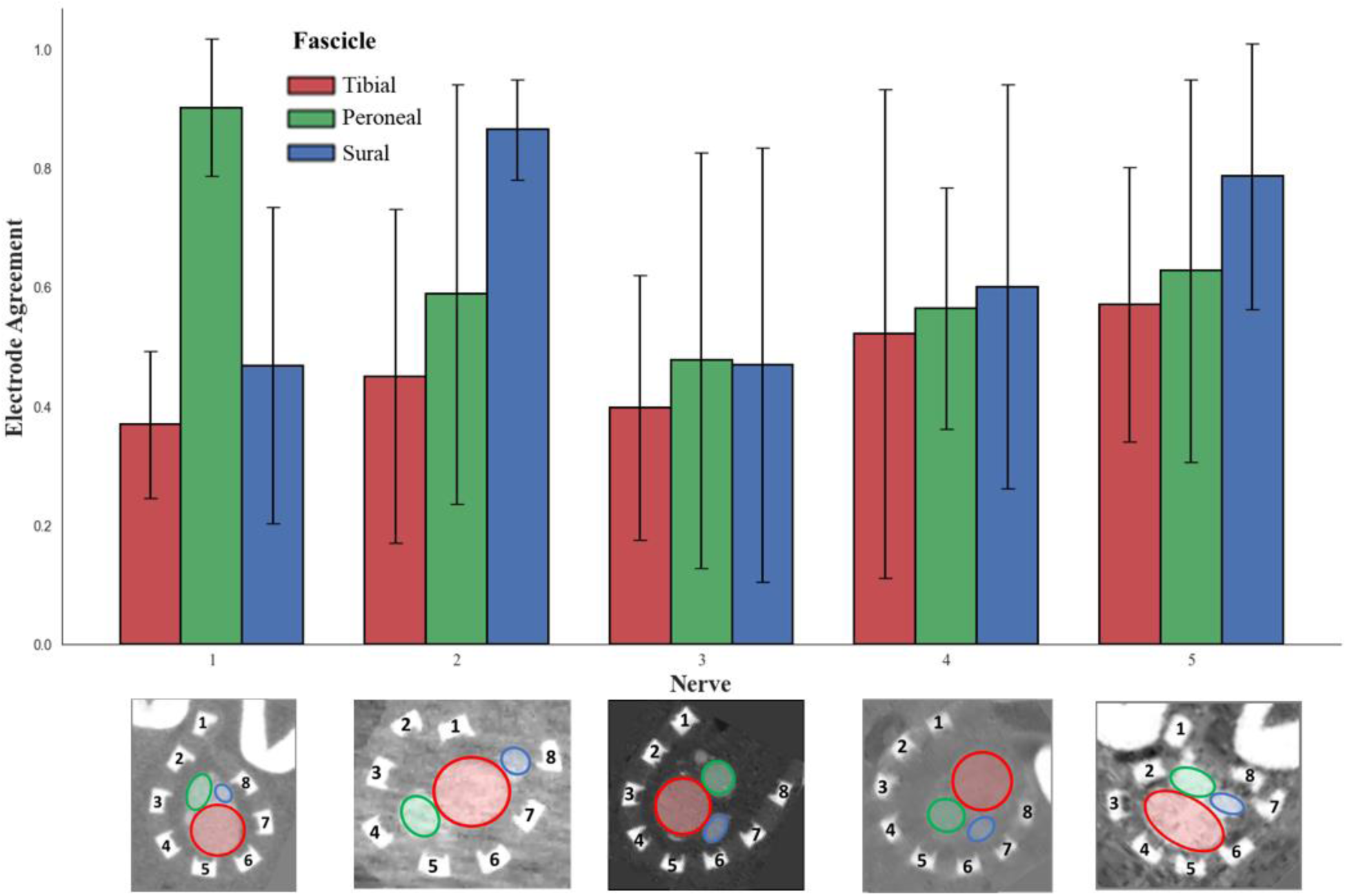
Electrode agreement metrics from the top 10 selectivity indices of each configuration, averaged together for each nerve. Error bars indicate 1 SD.

### Multivariate Regression Analysis

To identify similar trends across the anatomical variation, both *ex-vivo* and simulation selectivity were modelled as a function of stimulation waveform parameters (amplitude, pulse width, injected charge, phase symmetry), electrode configuration (bipolar or tripolar), and average electrode–fascicle distance (average of source and sink electrode distance to fascicle). Multivariate ordinary least squares (OLS) models were fit separately for *ex vivo* and *in silico* data, using only observations with positive selectivity for the fascicle under consideration. Across all fascicles in both the *ex vivo* and *in silico*, the average distance term had the largest coefficients and remained statistically significant after adjustment for all waveform terms (**Table 1**). All variables and coefficients can be found in Supplementary Material. *Ex vivo* average distance coefficients were negative for all fascicles, indicating higher selectivity at shorter distances from electrode pair to fascicle of interest. In the simulation, the average-distance coefficients remained the largest in magnitude but differed in sign from the *ex vivo* slope in the sural and tibial fascicles.

**Table 1.** Effect of Average Electrode-Fascicle Distance on Fascicular Selectivity.

| **Fascicle** | **Model** | **N** | **Coefficient** | **p** |
| --- | --- | --- | --- | --- |
| Tibial | Ex vivo | 14595 | -0.137 | $2.72 \times 10^{-46}$ |
| Tibial | In silico | 16791 | +0.173 | $1.74 \times 10^{-53}$ |
| Peroneal | Ex vivo | 23767 | -0.069 | $2.49 \times 10^{-3}$ |
| Peroneal | In silico | 17483 | -0.025 | $9.58 \times 10^{-58}$ |
| Sural | Ex vivo | 32465 | -0.172 | $1.00 \times 10^{-300}$ |
| Sural | In silico | 30022 | +0.320 | $1.00 \times 10^{-300}$ |

To better understand the differences between the coefficients in the two models, scatterplots were constructed comparing average fascicle-electrode distance to selectivity for each fascicle. Fascicle selectivity patterns aligned to anatomical size. In the tibial fascicle (largest), the simulation under-represented high-selectivity outcomes when the electrode pair was closer to a particular fascicle relative to ex vivo (**Figure 17a**), despite a small positive regression slope for average distance. The peroneal fascicle showed the closest agreement between modalities, with similar coefficient direction and almost complete overlap in the distance–selectivity plane, with just slightly higher in silico selectivity, but not specific to a particular distance. In the sural fascicle (smallest), the simulation predicted a narrower distribution of high selectivity at shorter distances than the ex vivo results produced, and exhibited a positive average-distance coefficient, consistent with selective outcomes appearing beyond just the closest geometries. Occupancy maps (**Figure 17b**) summarized these distributional differences by shading bins populated by only one of the two selectivity values. These findings reinforce the trend that the largest fascicle (tibial) is underpredicted, especially at lower average distances between active electrode and tibial fascicle. Additionally, the smallest fascicle (sural) is systematically overpredicted for selectivity at all distances.

**Figure 17.**
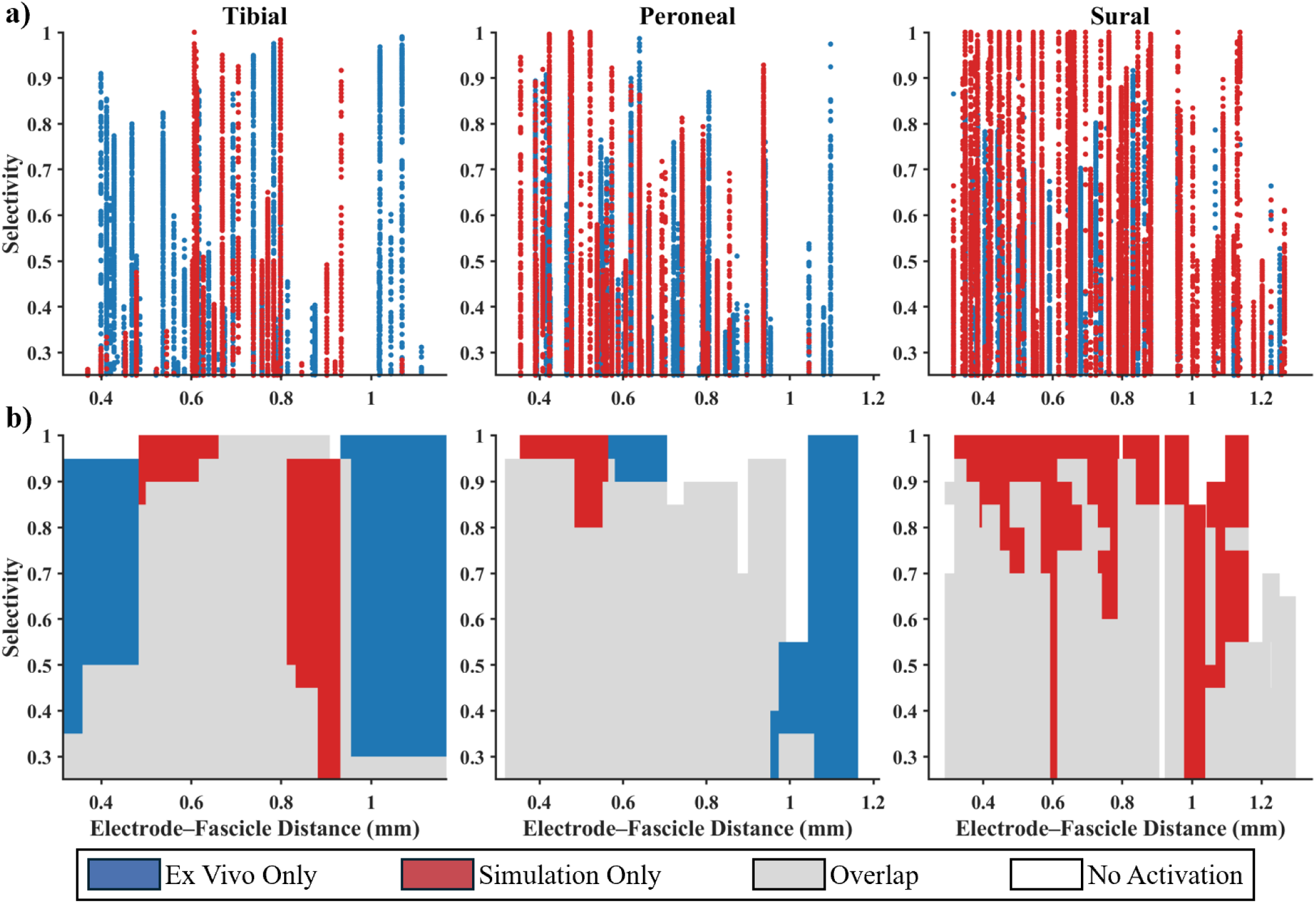
a) Scatterplots of all nerve stimulations that produced > 0.25 selectivity in a particular fascicle, comparing average fascicle-electrode (source and sink) distances to fascicular selectivity. Both *ex vivo* (blue) and simulation (red) selectivity are displayed. b) Occupancy maps that highlight the areas of the scatterplots where there was overlap (grey) between the *ex vivo* and simulation values. The areas where there was no overlap are colored blue (*ex vivo* only) or red (simulation only). Each scatterplot was broken up into grids with rectangular cell size: width at 20% of the fascicle size and height at a 0.05 selectivity increment. Each cell was classified as *ex vivo* only, simulation only, or overlap based on the presence of those respective points in the cell. A minimum threshold of 10 contiguous cells was imposed to highlight the larger clusters of similar selectivity trends.

## 5. Discussion

### Ex vivo selectivity with fully polymeric nerve cuff

The first objective of this study was to manufacture a multipolar fully polymeric nerve cuff to investigate *ex vivo* fascicular spatial selectivity. Across 5 nerve trials, a high degree (>0.65 SI) of selective stimulation was achieved, enabling discreet threshold activation for each of the three fascicles in the rat sciatic nerve. This systematic study assessed a wide range of stimulation parameters, identifying specific parameters in each trial that were able to achieve selectivity. This study shows similar results using the CE cuff compared to prior studies^27^ that have investigated spatially selective stimulation with metallic nerve cuffs. While spatially selectivity was not improved when compared to modern metallic cuff designs, these results support the wider use of fully polymeric nerve interfaces in neural interfacing applications by establishing performance parity.

### MicroCT informed simulated selectivity

A unique advantage of the fully polymeric nerve cuff is that it enabled accurate 3D reconstruction of the nerve bundle and the overlying nerve cuff using microCT for the first time. The Lugol stain typically used for tissue contrast enhancement in microCT^38^ was absorbed by the conductive elastomer (PEDOT:PSS/PU matrix) and allowed precise visualization of the electrode sites on top of the sciatic nerve without the detriment of artefacts inherent to metallic electrodes. Using the 3D reconstruction of both the electrodes and nerve bundle in COMSOL, this study then assessed the ASCENT simulation pipeline for accuracy in predicting the *ex vivo* selective stimulation results with more accurate electrode to fascicle geometric relationships. While the simulation does predict selective activation of each of the three fascicles with agreement across some specific waveforms within discreet nerve trials, when averaged across all nerve trials there was deviation in both the maximum achievable selectivity and electrode combination required to achieve selectivity, between the *in silico* prediction and *ex vivo* experiment. The sural fascicle was overpredicted for high selectivity (nearly SI = 1) for every single nerve, which in the *ex vivo* experiments achieved no greater selectivity when compared to the other two fascicles when averaged across the nerves (average SI ∼ 0.7). On the other hand, the tibial nerve was underestimated for the degree of selective activation, but only by about 15%. The peroneal did not differ significantly except in one nerve in which the fascicle was more distant from any of the overlying electrodes.

Based on the multivariate regression analysis, there is a clear relationship between capacity for selective activation and the size of the fascicle. The smaller sural fascicle was overestimated in the simulation, and the larger tibial fascicle was underestimated in the simulation, while the middle-size peroneal nerve was the most similar between simulation and ex vivo. This consistent size-dependent trend suggests an erroneous assumption in the construction of the fascicles in the simulation. For this study, the size distribution and density of fibres within each fascicle was kept constant, so the primary difference in the development of the model for the fascicles is the construction of the perineurium for each fascicle. There are many studies focused on the importance of modelling the perineurium accurately in peripheral nerve simulation experiments.^35,39,40^ The two main aspects of perineurium in modelling charge conduction are the perineurium thickness and the determination of an accurate equivalent circuit for the perineurial layer.^41^ In this study, the perineurial thickness scales linearly with the fascicle diameter, in line with findings from Raspopovic^42^. The linear relationship between fascicle diameter and perineural thickness used in this study was determined experimentally for the rat vagus nerve.^43^ One limitation of this study was the inability to measure the actual perineurial thickness from the microCT scans or other imaging modalities. To determine the significance of the effect of the perineurial thickness, a portion of the simulations were rerun comparing models with constant thickness for all perineurium,^44^ the published experimentally determined rat sciatic perineurial thicknesses,^45,46^ and thicknesses based on a linear equation that reduces the range of the thicknesses compared to the original range. None of these variations resulted in a model with a higher accuracy for predicting selectivity when compared to the *ex vivo* data. While improvements to the transfer function for the equivalent circuit of the perineural layer were not explored in this study, the impact of the perineural layer on the propagation of the electrical field was visualized to understand the overprediction of selectivity in fascicles with smaller diameters and the underprediction of selectivity in fascicles with larger diameters. The perineurial layer was modelled as a contact impedance with no physical thickness in the COMSOL model. Instead, the thickness of the perineurium was used to calculate the final resistivity of the perineurial layer (ρ_peri_ = 0.00088 S/m) with a sheet resistivity taken from Raspopovic et al.^42^ being a recent update of the original empirical determination of perineural resistivity by Weerasuriya et al.^47^ The attenuation and dispersion of the magnitude of the electric field across the boundary of the perineurium is shown in **Figure 18**. The homogeneity of the magnitude of the electric field is observed in each fascicle. This homogenization of electrical field within the fascicle has been well documented and investigated with respect to selectivity studies in simulated environments and has been attributed to high perineurial thickness and high perineurial resistivity values.^35^ A fascicle size-dependent bias is consistent with these model assumptions that homogenize intrafascicular fields and boundary properties relative to biological preparations. In smaller fascicles, such assumptions can decrease activation thresholds of axons much further away compared to others that are in closer, larger fascicles, which functions to attenuate local recruitment in those larger fascicles. While a lot of effort has been put into determining the precise relationship between the fascicular diameter and the perineurial thickness, the difference between **Figure 18a** and **Figure 18b** demonstrates that there is very little impact on the theoretical electric potentials generated in the COMSOL model. When comparing **Figure 18a** and **Figure 18c** (most commonly used ρ_peri_ = 0.021 S/m in literature),^42^ it is evident that impact of the perineurial resistivity is much higher than the impact of the relationship between perineurial thickness and fascicle diameter. This study provides *ex vivo* data that can be used to interrogate future models of perineurial resistivity.

**Figure 18.**
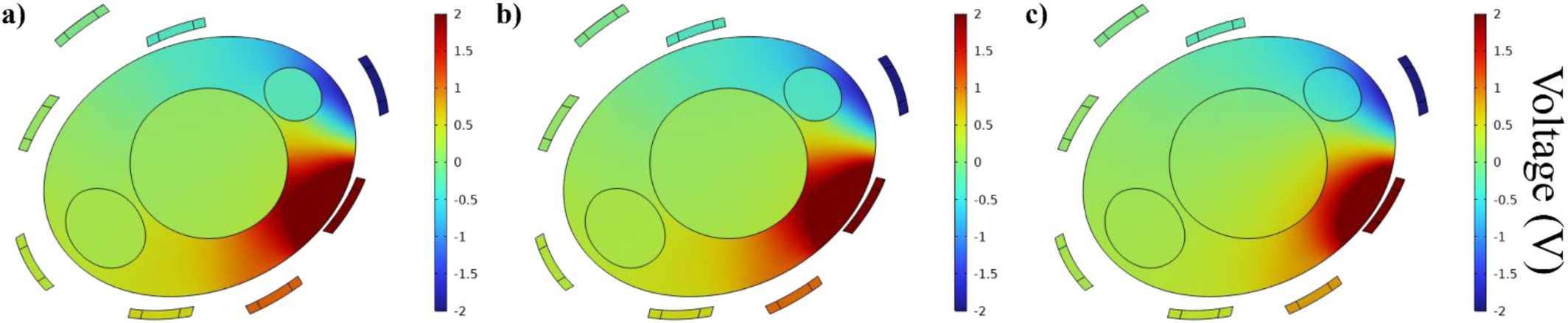
Comparison of quasi-static electric potential (V) solutions when varying perineurial parameters. a) Uses scaled perineurial thickness and perineurial resistivity from Raspopovic et al (default in ASCENT) b) Uses a constant perineurial thickness (10 μm) and default resistivity. c) Uses scaled perineurial thickness and perineurial resistivity from Weerasuriya.

While the perineurium is likely to have an impact on the fascicular selectivity of nerve cuffs in ASCENT and should be modelled as accurately as possible, it is possible that other assumptions in the model may also have an impact. While there are many advantages to using the conductive elastomer-based electrodes, one limitation is that they are not as well characterized in modelling systems compared to metallic electrodes. The charge transfer characteristics of metal electrodes are well understood and operate primarily with double layer capacitance within the biological environment. However, the volumetric conductance of a conductive elastomer incorporates both ion mobility and electron transfer through bipolaron formation within the doped polymer backbone.^48^ This complicates the accurate modelling of the current injection from the active electrodes. Similar to the perineurial changes, incorporation of a more accurate model of charge transfer across the CE electrode boundary may lead to greater similarity between the model and *ex vivo* selectivity data. Another potential limitation in the model is the fact that the solution to the current injection input is a quasi-static approximated solution. A dynamic solution would more accurately account for the capacitive behavior of the electrodes and the time-based parameters like stimulation frequency. This study used a 25 Hz frequency of *ex vivo* stimulation (40 ms between each biphasic pulse). The approximated quasi-static solution does not take into account the frequency to lower the computational demand for an exact solution. While a 40 ms pulse separation should be sufficient for stabilisation after polarisation, and the order of the amplitude sweep was randomised to prevent ramping effects, this is still an assumption that differs from real world implementation and must be considered.

### Stimulation Paradigms and Device Designs for Selectivity Improvements

While almost every fascicle in each nerve was able to achieve at least a selectivity index of 0.45 in the *ex vivo* experiments, the differences in neuroanatomy led to bigger differences in the maximum selectivity achieved compared to the differences in waveform configurations. There are two potential avenues which are proposed for achieving higher selectivity in peripheral nerve stimulation with a fully polymeric device. One approach is to further investigate changes in the stimulation technique, and the other is to implement changes in the device design and interface with the target tissue.

In this study, nearest neighbour bipolar and tripolar stimulation was employed across both even and uneven biphasic waveforms. Tripolar configurations gave a slightly higher selectivity in one fascicle, which aligns with literature by localizing the electric field more than bipolar configurations. Higher selectivity could potentially be achieved with more advanced stimulation paradigms like temporal interface or intermittent inferential current stimulation.^49,50^ This technique would not require any significant changes to the device design as it relies on the constructive interference of sinusoidal currents from two distinct bipolar pairs. The hotspot of suprathreshold extracellular voltages is contained between the pairs, with the exact location determined based on the ratio of the amplitudes of the sinusoidal waves, so circumferentially spaced electrodes are well suited to target deep fibers without activating superficial ones. The selectivity could then be increased further reducing the size of the electrodes in the circumferential spacing and increasing their density. Any stimulation technique would likely benefit from further miniaturization of these fully polymeric electrode designs. Miniaturization of electrodes heavily relies on higher precision and control in micromachining fabrication methods, which are enabled in this study by lasers with femtosecond pulse widths and sub 10-micron spot sizes.

Device design aspects could also improve selectivity, including the development of more reliable closure mechanisms that ensure complete and conformal wrapping of the entire nerve diameter. It was clearly demonstrated in this study that conventional suture methods can result in varied levels of electrode contact with the tissues, but approaches in which heavy or tight pre-formed cuffs constrain the nerve raise concerns with respect to long-term nerve health. Some promising approaches for improved closure without compromising nerve health and that can be adapted to polymer devices include a ratcheted belt loop approach^51^ or incorporation of photochemical-based tissue adhesion.^52^ Fully polymeric arrays enable a wide variety of options for chemical modifications that can target adhesion between devices and adjacent tissues.

Finally, selectivity could also be improved with changes to the way the device interfaces with the target tissue. With laser based micromachining, rapid iteration of fabrication is enabled with the capacity to precisely control the layout and form factor of the electrode array. As such custom devices can be designed for specific peripheral nerve anatomy. For example, penetrating, interfascicular electrodes can be designed for implantation at specific locations where the nerves branch and the fascicles are more distinct under the epineurium. This would permit the electrodes to be placed close to the target fibers while minimizing damage to the nerve by only going between fascicles. Alternatively, regenerative peripheral nerve interfaces (RPNIs), in cases where amputation is required, can be utilised to create interfaces where regenerated and branched nerve fibers are more isolated and accessible. RPNIs are surgical constructs that provide implanted free muscle and/or skin grafts into which transected peripheral nerves regrow. These constructs both contain the nerve to a known location while also allowing them to rebranch out into the de-innervated grafts. As such electrode arrays can be specifically designed to conform to the grafts, increasing the number of independently addressable fibers. Ultimately, this approach may provide an opportunity for bidirectional neural interfacing with high channel counts for the restoration of sensory perception and control of limb prostheses.

## Conclusion

This study demonstrates the manufacture and functionality of a fully polymeric transverse multipolar electrode array for selective fascicular activation in a rat sciatic nerve. It directly compares the *in silico* simulation and predicted fiber activation to experimentally determined fascicular activations from an *ex vivo* rat sciatic nerve study. It was demonstrated that a high degree of fascicular selectivity with the fully polymeric nerve cuff in the rat sciatic nerve can be achieved. However, while the model also predicts that fascicular selectivity can be achieved, when uniquely informed by microCT, it does not accurately predict the selective activation of a particular fascicle given the location of an active pair of electrodes and a specific stimulation waveform. While the ASCENT pipeline can inform comparative electrode designs to achieve higher degrees of selectivity, more accurate *in silico* models of charge transfer mechanisms may be required for both perineurium impacts and non-metallic electrode charge injection. While varying the perineurium thickness was not sufficient alone, improving the charge transfer function for this cellular component may provide greater modelling accuracy. Specific to this study, mirroring the mixed ionic and electronic charge transfer of CE-based electrodes may improve similarities between the ASCENT pipeline and *ex vivo* peripheral nerve stimulation studies.

## Experimental Section/Methods

### Nerve Cuff Manufacture

Fabrication of the nerve cuffs was based on the work done by Cuttaz et al.^14^ and is depicted in **Figure 19**. Polydimethylsiloxane (PDMS) (Sylgard 184, Dow Chemical Company, GER) was mixed at a 10:1 ratio of monomer to curing agent. While the PDMS was degassed, a release layer of polystrenesulfonic acid (PSSA) was spin-coated (Photo Resist Spinner, Electronic Micro Systems, EMS 6000, UK, 600 rpm for 10 s and 1500 rpm for 45 s) onto a glass slide and left in an oven to cure at 60°C for 15 minutes. After complete degassing, the PDMS was then spin-coated on top of the PSSA release layer at 600 rpm for 10 s and 2000 rpm for 45 s. The glass slide was left in an oven at 50 °C for 20 min while the PDMS cured. At this point, the PDMS remained tacky, and a CE sheet was placed on the PDMS on the glass slide. For this work, 25 wt% CE sheets were fabricated using specifications from Cuttaz et al.^13^

**Figure 19.**
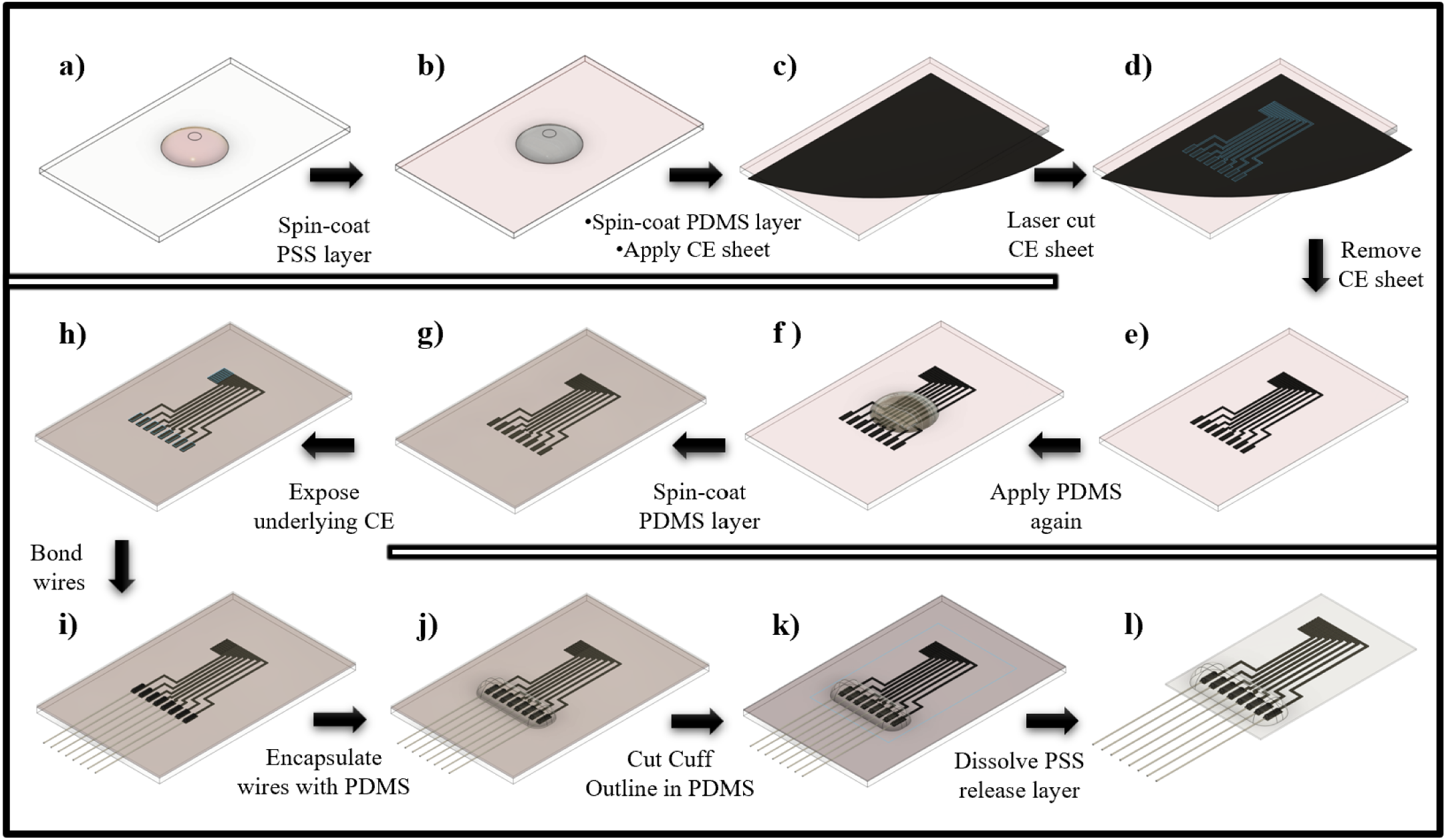
a) 2 mL of PSS was spin-coated onto the glass slide. b) After 15 minutes on heat, 1 mL of PDMS was spin-coated on top of cured PSS layer. c) Once the first PDMS layers cures for about 20 minutes and becomes sticky, a CE sheet was manually applied on top. d) The laser pattern was cut into the CE sheet on top of the glass slide. e) The CE sheet was manually peeled away from the underlying PDMS layer, leaving behind the laser cut patterned electrodes. f) A second 1 mL of PDMS was added on top of the CE electrode pattern and spin-coated. g) Second layer of PDMS was left on heat to cure overnight. h) The PDMS over the active sites and wire bonding was removed by laser ablation. i) Silver wires were bonded to the exposed wire-bonding sites. j) The wires and wire-bonding sites were encapsulated with PDMS and glass slide was left on heat overnight for PDMS to fully cure. k) Rectangular shape of nerve cuff was manually cut around the electrode pattern. l) Nerve cuff is removed from glass slide after a water bath dissolved the PSS release layer. Adapted from Cuttaz et al.^[15]^

After smooth rolling the CE sheet on the PDMS, the glass slide was aligned and fixed in place under a laser, and the CE was laser cut (Coherent Monaco 515 nm, Optec Lasea Group, Power: 80%, Freq: 333 kHz, Speed: 800 mm/s, Loop Count = 10, Layer Count Delay = 20 ms) into wire bonding sites, electrode active sites, and the tracks between them. Then the CE sheet was manually peeled off the PDMS layer, leaving the electrode array on the PDMS. Then, a second layer of PDMS was spin-coated on top of the CE layer and the whole slide was left to cure overnight at 45°C, resulting in a CE patterned electrode array in between two insulation layers of cured PDMS. The sites for bonding wires and the active sites of the electrodes were then exposed. The larger wire bonding sites were exposed manually with fine forceps. One refinement of the manufacturing process established by Cuttaz et al.^14^ for fully polymeric nerve cuffs was undertaken: the PDMS over the active sites was removed by laser ablation (Coherent Monaco 515 nm, Optec Lasea Group, Power: 45%, Freq: 333 kHz, Speed: 800 mm/s, Loop Count = 10, Layer Count Delay = 40 ms), as opposed to manual removal of the PDMS used previously. The active sites were too small to be manually exposed, and the laser ablation technique enabled high precision exposure of active and bonding sites. Any debris from PDMS ablation was then removed with scotch tape. Following the laser ablation, the PDMS surrounding the electrode array was cut into a 30 x 11 mm rectangle, leaving the CE electrode array in the middle of the rectangle. To remove the PMDS-encapsulated electrode array, the glass slide was then submerged in deionized water to dissolve the PSSA. Forceps were used to peel the PDMS array off the glass slide. Silver wires (125 µm Ø, polytetrafluoroethylene insulated, Advent Research Materials, UK) were bonded to the wire bonding sites of the electrode array. The bonding area with the silver wires was then encapsulated with silicone (MED 4-4220, NuSil Technology, LLC, USA) and left to cure overnight. Finally, PDMS pads were added to the top of the electrode array using the Nusil silicone as an adhesive to aid in handling of the electrode array during the placement onto the nerve.

### Electrochemical Performance of Nerve Cuffs

Each nerve cuff device was characterized by electrochemical impedance spectroscopy (EIS) and cyclic voltammetry (CV) to assess device performance and to ensure consistency in the cuff manufacturing process. Both tests were run using a commercial potentiostat (PGSTAT101, Metrohm Autolab, NLD). EIS and CV measurements were taken at 2 timepoints: immediately post-manufacture and when located on the *ex vivo* nerve. For each testing point, the same 3-electrode setup was used: a platinum wire (0.5 mm Ø) counter electrode, a leakless Ag|AgCl reference electrode, and each of the 8 channels for the working electrode. The post-manufacture testing was performed in Dulbecco’s phosphate buffer solution (DPBS) while the *ex vivo* testing was performed in a modified Krebs-Henseleit buffer (oxygenated by gas dispersion tube using carbogen (95% O_2_, 5% CO_2_), composed of NaCl (113mM), KCl (4.8mM), CaCl_2_•H_2_O (2.5 mM), KH_2_PO_4_ (1.2 mM), MgSO_4_ (1.2mM), NaHCO_3_ (25mM), and Dextrose (5.55 mM), Sigma). EIS was measured using a 30 mV sinusoidal waveform at frequencies ranging from 1 Hz to 100 kHz. CV was measured with a voltage sweep from -0.6 V to 0.8 V for 6 cycles to allow for stabilization. Impedance magnitude (at 1 kHz) and phase shift waveform, from EIS, alongside charge storage capacity (CSC), from CV, were compared across devices and across manufacturing points to ensure manufacturing consistency and electrochemical performance. To ensure that the laser ablation of PDMS did not damage the underlying active sites, CSC values were obtained in the same manner as above for electrodes that had been exposed by laser ablation vs manual removal. The size of the active sites in these test electrodes (n=5) was scaled up by 100% from the final size of the active sites in the nerve cuff to ensure that manual removal was possible.

### Electrode Geometry

To assess spatial selectivity of a nerve bundle with a nerve cuff, a transverse electrode array was designed (Fusion 360, Autodesk, Version 2.0.13881) and laser cut into the CE during nerve cuff manufacture. The transverse array contained one ring of 8 electrodes (0.3 x 2 mm) spaced evenly, with an inter-electrode gap of 0.15 mm. The design can be seen in **Figure 1**. All sizing was based on the average size of a female rat (Sprague-Dawley) sciatic nerve (0.9-1.1 mm Ø at rat weight of 200-230 g).

### Sciatic Nerve Dissection

Sciatic nerves were harvested from female Sprague-Dawley rats weighing 200 – 230g (Charles River, UK). All procedures were carried out under a project license approved under by the Home Office Animals in Science Regulation Unit and in accordance with the Animals (Scientific Procedures) Act 1986 (PPL: PA98EF5E9). Rats were anaesthetised with isoflurane (5% at 2.5 L/min for ∼5 minutes) prior to humane culling. Both sciatic nerves were then harvested from the origin at the L4-S3 roots, down to the distal ends of the 3 largest branches of the sciatic nerve: the tibial nerve (TN), the peroneal nerve (PN), and the sural nerve (SN). To begin, the skin on the posterior aspect of the rat hind limb was incised to expose underlying biceps femoris and superficial gluteus. The fascial layer between those two muscles was incised to expose the sciatic nerve. The nerve was then freed from surrounding tissue and followed up to the point of exit from the sciatic notch in the posterior pelvis. Thin motor branches that innervate the semitendinosus, semimembranosus, and biceps femoris were sacrificed as the sciatic nerve was freed toward the sciatic notch. Distally, the sciatic nerve was dissected down to the branching points of the TN, PN, and SN in the fatty popliteal fascia. At this point, the distal end of each branch was located (superficially at the lateral malleolus for SN, just proximal to the tarsal tunnel for TN, and the posterior aspect of the fibular head for the PN). Each branch was dissected proximally from these distal points, all the way back to the popliteal fossa where they branched from the sciatic nerve. With all of the branches freed from surrounding tissue, the sciatic nerve was transected at the sciatic notch. The whole nerve (seen in **Figure 20**) was placed in a chilled, oxygenated modified Krebs-Henseleit Buffer solution. Then, the nerve was cleaned under a microscope to remove excess fat before being transported to the *ex vivo* nerve chamber. Both sciatic nerves in each animal were extracted using the same protocol.

**Figure 20.**
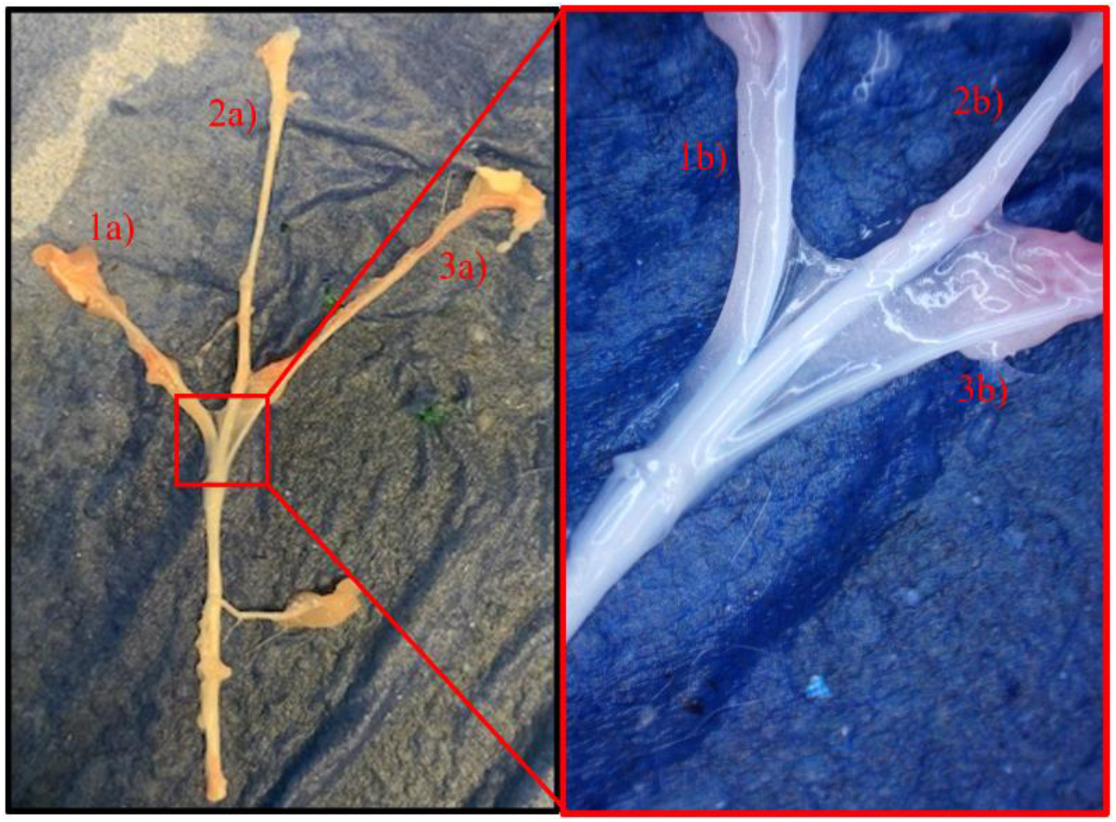
Excised rat sciatic nerve showing the branching just proximal to the popliteal fossa. 1a) Macroscopic view of peroneal nerve. 2a) Macroscopic view of tibial nerve. 3a) Macroscopic view of sural nerve. 1b) Microscopic view of peroneal nerve. 2b) Microscopic view of tibial nerve. 3b) Microscopic view of sural nerve.

### Ex Vivo Nerve Chamber Setup

After the nerve was extracted, it was cleaned of remaining fascia or muscle using microsurgical forceps and scissors (Fine Surgical Tools, InterFocus Ltd, UK). Then, the open end of each nerve was sutured closed to prevent fluid ingress into the fascicles. Finally, the nerve was placed into a custom-designed nerve chamber made from acrylic, similar to one used by Cuttaz et al.^14^, but increased in chamber size to accommodate neural recording from three independent branches. To put the nerve into the chamber, both sides of the nerve chamber were first filled with an oxygenated modified Krebs-Henseleit (mKH) buffer (37 °C). The proximal end of the sciatic nerve was then fed through the 1.2 mm hole in the wall between the two chambers to separate the stimulation and recording chambers. The feedthrough hole was then blocked with silicone-based grease to prevent fluid leakage, then the buffer in the recording chamber (with the 3 branches in it) was replaced by mineral oil (Sigma) to electrically isolate the custom-made bipolar silver hook electrodes used to record neural activity. Once the nerve was fixed in place in both chambers with stainless steel insect pins, the nerve cuff was wrapped around the nerve. The cuff was pulled around the nerve with the PDMS reinforcement pads on the tail of the cuff. Once wrapped around the nerve, the cuff was clamped with a disposable microvascular clamp (TKM-1, Synovis MCA, USA) to secure the cuff fit during stimulation. In the stimulation chamber, the mKH buffer was constantly drained from the chamber using a peristaltic pump (Pump P-1, Pharmacia Biotech, Sweden) and then pumped back into a bottle where the buffer was reoxygenated and then fed back into the nerve chamber. In the recording chamber filled with mineral oil, each branch was placed onto a separate custom-built bipolar Ag, AgCl-coated hook electrode. The two hooks on each bipolar recording electrode were fixed 5 mm apart to ensure consistency across each fascicular recording. Each bipolar electrode provided an independent input into a differential gain amplifier (DP-304A, Warner Instruments, USA, Gain: 1000, Low-Pass: 5 kHz, High-Pass: 300 Hz). A separate Humbug (Digitimer, USA) was used to remove 50-60 Hz noise from each signal. Finally, each signal was digitized by a data acquisition interface (Micro3 1401, Cambridge Electronic Design Ltd., UK) and processed and saved using Spike2 software (Version 9.16, Cambridge Electronic Design Ltd., UK).

### Stimulation Protocol

In this study, multiple waveform parameters were explored to achieve maximum selectivity. Stimulation was controlled by a stimulus generator (Multichannel Systems 4008, 8-channel, Harvard Bioscience, MA, USA) All waveforms were biphasic, but the number of poles was either two (bipolar) or three (tripolar), and the symmetry of the phases was either even or uneven. An amplitude and pulse width sweep were performed for each stimulation configuration. All phases were charge balanced by keeping the ratio of the amplitude and pulse widths equal across phases. Because the orientation of the nerve fascicles to the overlying electrodes is not known *a priori*, the stimulation protocol was applied to all sides of the nerve, through each electrode. Only nearest neighbor electrode couplings were investigated here, that resulted in 8 different electrode positions to test for each waveform parameter variation. The pulse width sweep extended from 50 µs to 300 µs. For each pulse width and electrode position, a range of input currents (50 µA to 1600 µA, 50 µA increments) was used to determine a recruitment curve for each fascicle within the sciatic nerve. The order of the input currents was randomized to avoid potential temporal ramping bias in the nerve response. Each input current was repeated 5 times non-sequentially and the repeat neural responses were then averaged to reduce noise.

### Data Processing

Each stimulation pulse was synced with a trigger pulse that initiated data acquisition 0.2 ms before the stimulation rising edge until 0.5 ms after the stimulation rising edge. After averaging over the 5 repeats, the neural responses were clipped to isolate the stimulation artifact and neural response. The clipped neural response data was then smoothed with a moving average to ensure that the peaks would be located with the peak finding function built into MATLAB. The minimum and maximum of the neural response were then obtained to calculate the peak-to-peak voltage (*V*_*pp*_) of each generated compound nerve action potential (CNAP). The maximum CNAP for each fascicle per nerve trial was then obtained from all stimulation parameter combinations and used to normalize each fascicular activation respectively.

### Selectivity Analysis

To calculate fascicular selectivity for each configuration within a single waveform parameter study, **Equation 1** was employed, assuming each of the three fascicles is the target nerve for the respective selectivity calculation.^27^

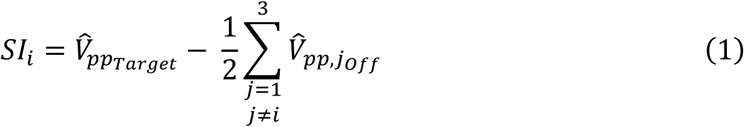

In this equation, selectivity indices (SI) for each fascicle are obtained by subtracting the average of the normalized CNAPs of the off target fascicles (*V̂*_*ppOff*_) from the normalized CNAP in the target fascicle (*V̂*_*ppTarget*_). A selectivity index of 0 indicates equal recruitment of all three fascicles. A positive value of the selectivity index indicates greater recruitment of a single fascicle compared to the other two. A value of 1 would indicate complete activation of a single fascicle and no activation of either of the other two fascicles, demonstrating optimal fascicular selectivity. Finally, a negative value of selectivity indicates a greater recruitment of the other two fascicles compared to the target fascicle. Because only positive values of selectivity indicate selective recruitment of single fascicles, only positive values of selectivity are considered in the results. To best visualize the impact of waveform parameters, electrode locations, and stimulation amplitude on selectivity, RGB heatmaps were produced in which single pixels represent the activations of each of the fascicles at a particular amplitude and a particular variation of a waveform parameter that is being investigated. The normalized activations were mapped from the range 0 to 1 onto a range 0 to 255 to easily map as red, green, and blue colors in the heatmaps. The overlay of each of the colors in the heatmap is color representation of selectivity.

### CT Scan

Following the completion of the stimulation protocol, the nerve was fixed in alginate (2:1 mixing ratio, Industrial Plasters Ltd, UK) that was set using a buffered iodine solution to ensure contrast staining for the microCT. The buffered iodine solution was prepared by mixing equal parts Lugol’s solution (18% w/v, Sigma-Aldrich, UK) and Sorsenson’s Buffer (3:1 ratio of 266 mM Na_2_HPO_4_ and 266 mM KH_2_PO_4_). The buffered Lugol (B-Lugol) solution was used to stabilize the pH and reduce the shrinkage of the tissue during staining.^38^ After 5 minutes, the alginate had hardened around the nerve cuff and nerve, and the whole block of alginate was removed from the nerve chamber and left to soak in Lugol solution for 60 hours to improve the contrast staining of the fascicles in the sciatic nerve bundle. Before the microCT scan, the sample was removed from B-Lugol solution, patted dry, and wrapped in cling film to maintain moisture content of the alginate during the scan. The samples were then imaged using a microCT scanner (Xradia 510 Versa, Zeiss, GER, 0.4x objective, 80 kV, 7 A, 6 s exposure). The orientation of the electrodes and the underlying fascicles was determined for each nerve and the distances between each electrode and each fascicle were measured to include in the multivariate regression analysis. Additionally, the exact fascicular structure in each nerve was fed back into the ASCENT pipeline to determine if the simulation results would more closely match the *ex vivo* results if the exact fascicular structure is known *a priori*.

### ASCENT model

The ASCENT pipeline (version 1.3.1) was used to simulate the *ex vivo* experiments presented in this study.^20^ The pipeline was in line with the latest methodology employed by the developers of the simulation pipeline in their validated preclinical animal model studies.^53^ Briefly, COMSOL v5.6 (COMSOL Inc., Cambridge, UK) was used to construct finite element models (FEMs) of each nerve, each nerve cuff, and the surrounding medium within the *ex vivo* stimulation chamber. These FEMs were used to solve the electric potentials in each nerve in response to current-driven stimulation delivered through the electrodes wrapped around the nerve. These extracellular potentials were then applied to cable models of mammalian myelinated and unmyelinated fibers in NEURON v7.6.^54^ The cuff was placed in the middle of the nerve length (12.5 mm), and action potentials were detected at 90% along the length of the nerve (at location z = 11.25 mm). An action potential was detected when the transmembrane potential crossed -30 mV with a rising edge. Activation thresholds were then determined with a binary search strategy. These activation thresholds were then used to determine which fibers activated given a specific stimulation amplitude. The percentage of fibers that activated in each fascicle was then used as the normalized activation of each fascicle, and fascicular selectivity in simulation was determined with the same selectivity index formula in *Selectivity Analysis*. There were a few key distinctions in these simulation studies compared to those in the validated rat model studies, reported by Musselman et al.^53^ Unique nerve morphology and trial-specific cuff electrode geometry obtained from the CT scan were constructed in COMSOL within the ASCENT pipeline. Custom ellipsoidal part primitives were generated for each trial to ensure accurate modelling of the fit of the cuff electrode array around each nerve. The boundaries of the media were assigned to ground to replicate the grounding electrode at the edge of the nerve chamber. Rather than platinum, the primary conductive material was conductive elastomer with a conductivity of 660 S/m. The surrounding media were assigned to be modified Krebs-Henseleit Buffer solution with an electrical conductivity of 2 S/m to mimic the *ex vivo* nerve stimulation chamber. A list of all material assignments with their assigned conductivity values can be found in **Table 2**. Lastly, stimulation configuration files were generated for each unique waveform to match the *ex vivo* stimulation paradigms.

**Table 2.**
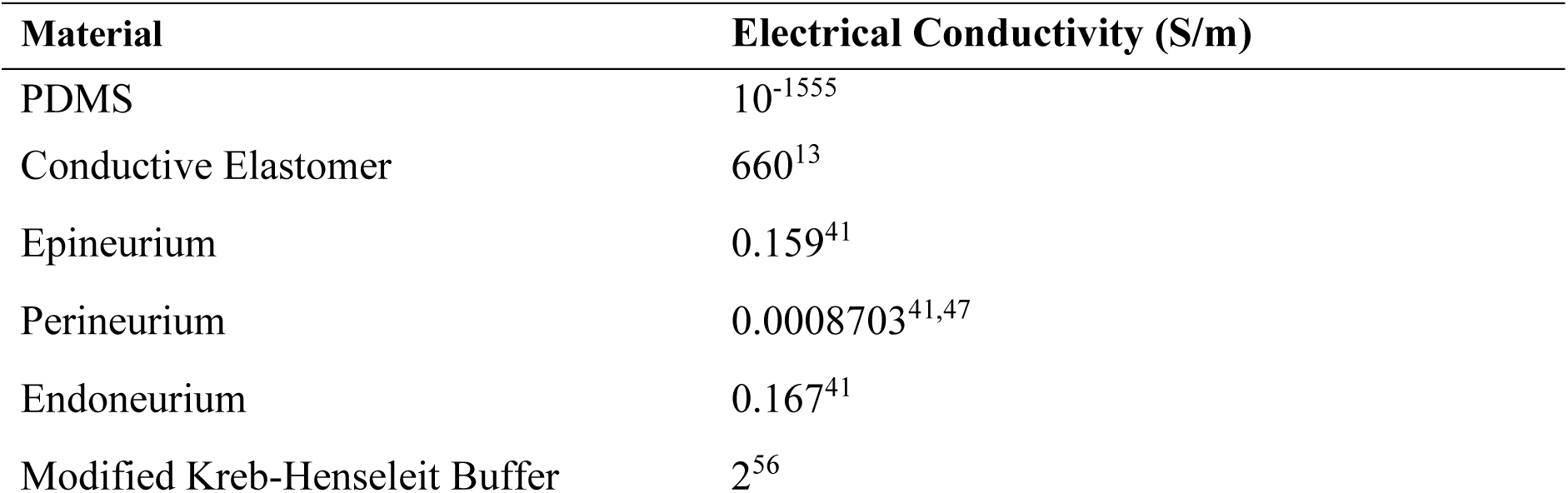
Material assignments and conductivities in the COMSOL FEMs.

## Supporting information

Heatmaps for all nerves

## Supporting Information and Data Availability

The authors declare that the data supporting the findings of this study are available within the paper, its supplementary information files, and a Github repository created for this project. (https://github.com/zkb17/Neuromodulation_Peripheral_Nerve_Polymeric_Cuff)

## Author Contribution Statement

Z.K.B. led the data curation, analysis, investigation, methodology, visualization, and manuscript drafting, with equal contributions to conceptualization and software. Z.N. contributed equally to analysis, investigation, methodology, and visualization, with additional support for data curation and review. E.A.C., I.B.M., C.A.R.C., A.R., J.G., A.Ro., and T.C. provided supporting contributions across conceptualization, methodology, investigation, supervision, resources, and manuscript review, while R.A.G. provided lead supervision, funding acquisition, project administration, and equal contributions to review.

## Ethics Declaration

All procedures were carried out under a project license approved under by the Home Office Animals in Science Regulation Unit and in accordance with the Animals (Scientific Procedures) Act 1986 (PPL: PA98EF5E9).

