## Supplementary material for "Neuromodulation of a peripheral nerve using fully polymeric cuff electrodes: Understanding predictability of selective stimulation": Heatmaps for all nerves

### *Activation Heatmaps*

Figures are organized by nerve first (Nerve 1–5). Within each nerve, configurations are ordered as 1–5. For every configuration, the ex vivo figure is presented immediately followed by the simulation figure.

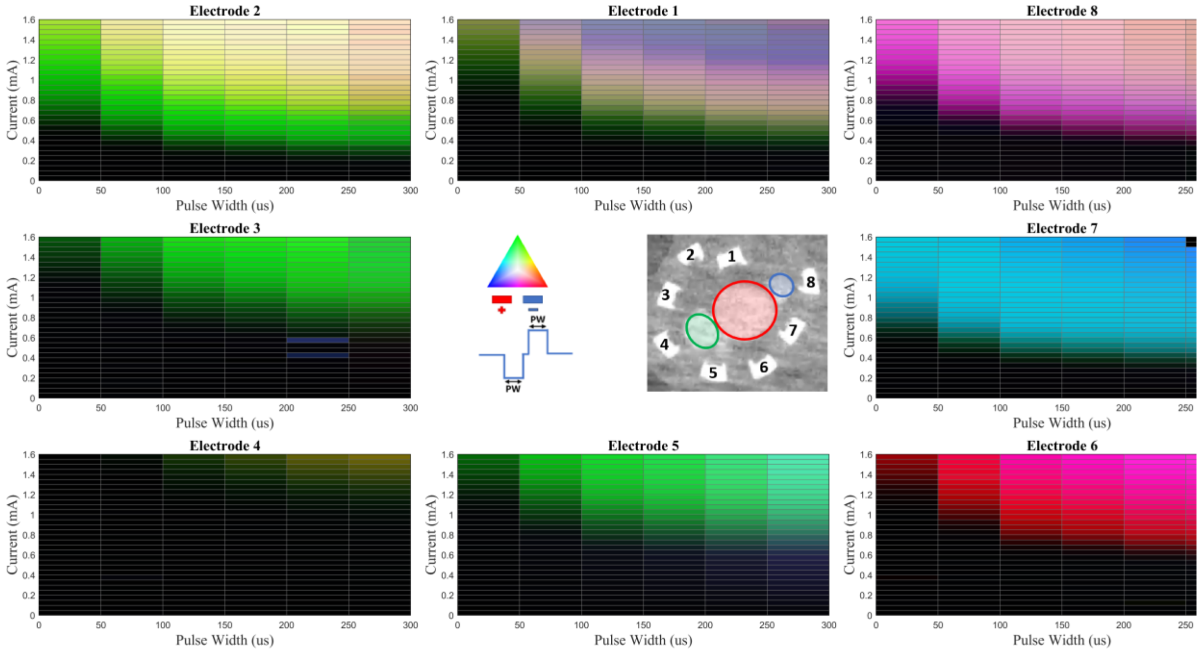


**Figure S1.** Nerve 1 – Config 1 (Ex Vivo)

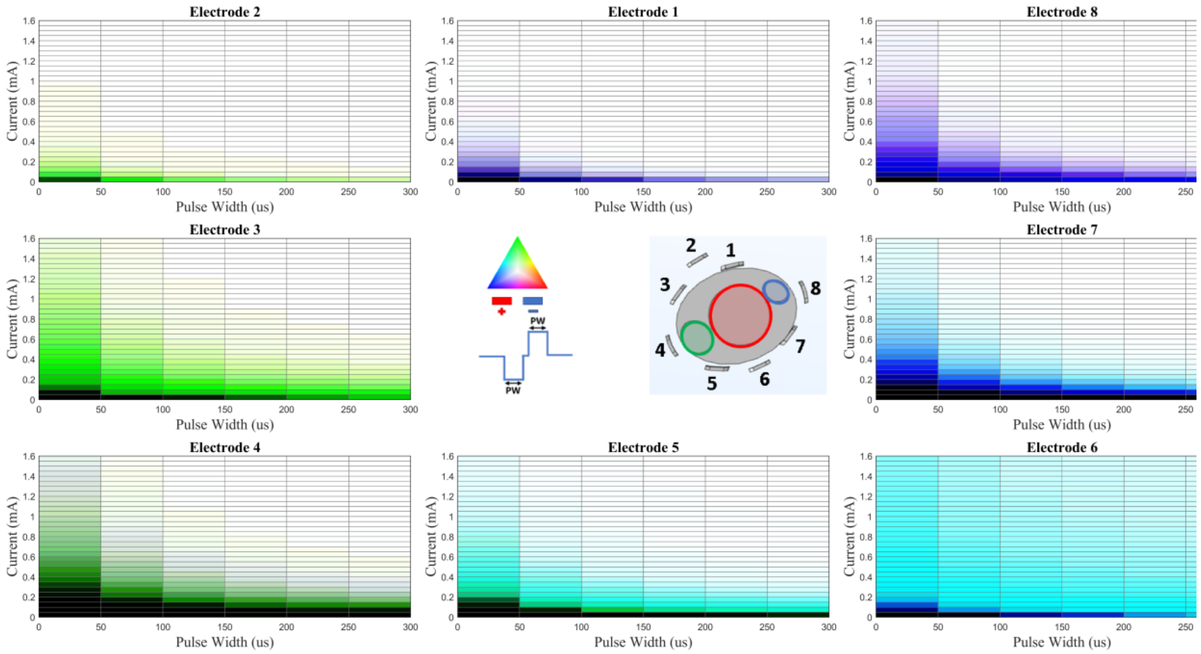


**Figure S2.** Nerve 1 – Config 1 (Simulation)

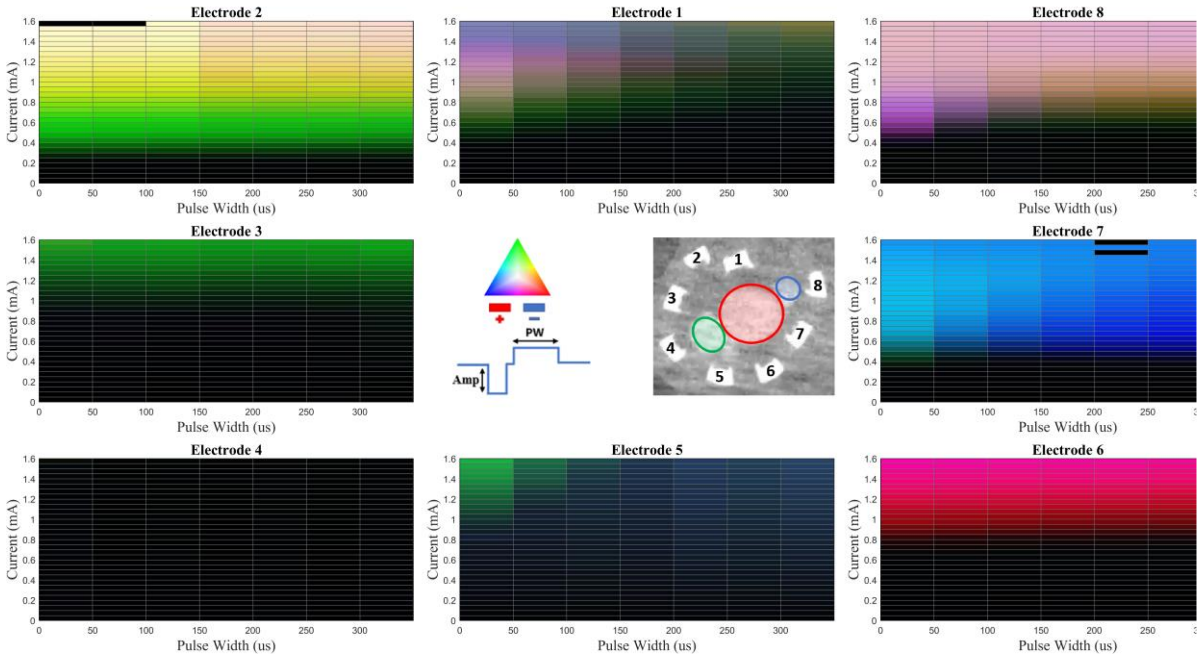

**Figure S3.** Nerve 1 – Config 2 (Ex Vivo)

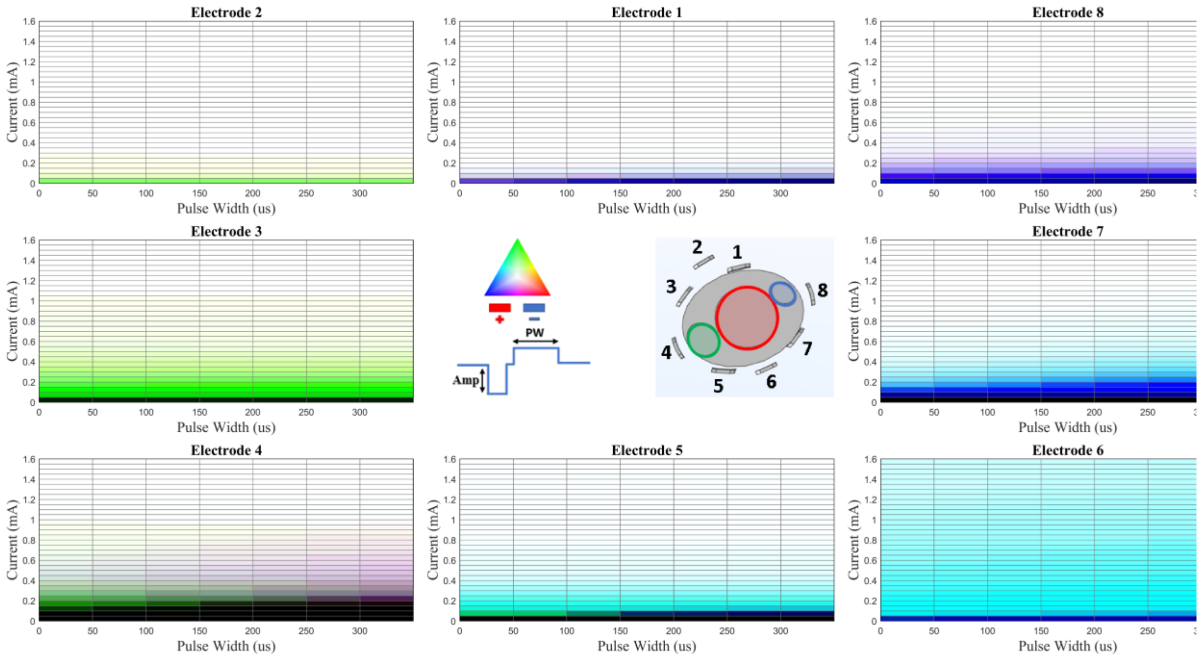

**Figure S4.** Nerve 1 – Config 2 (Simulation)

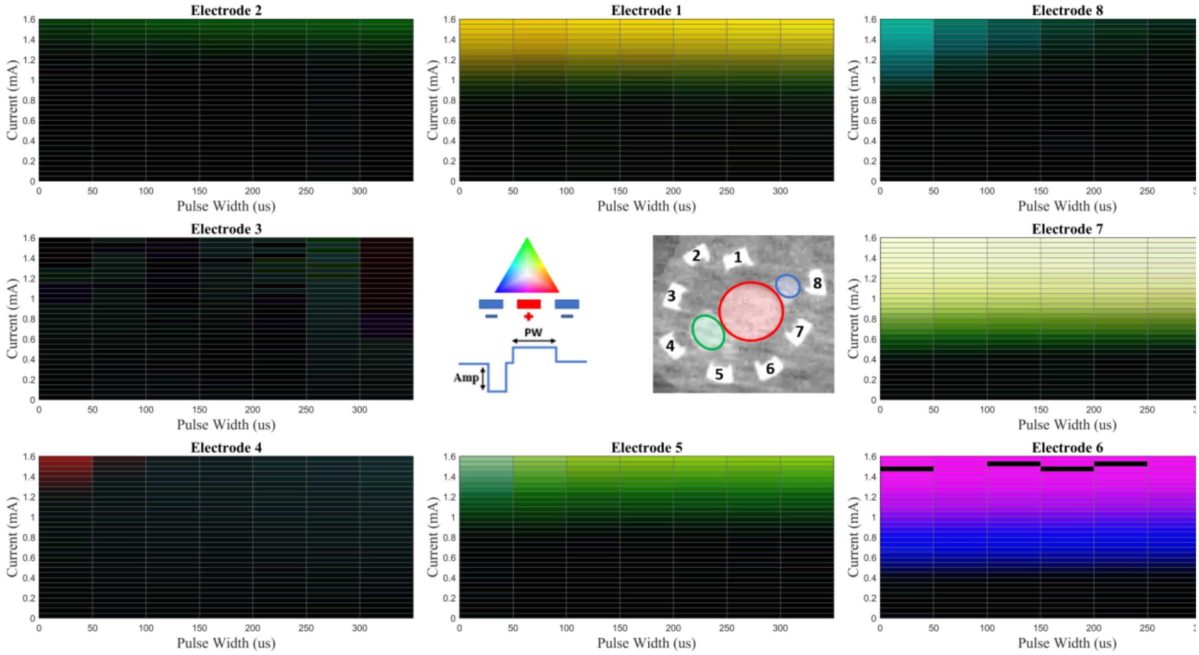


**Figure S5.** Nerve 1 – Config 3 (Ex Vivo)

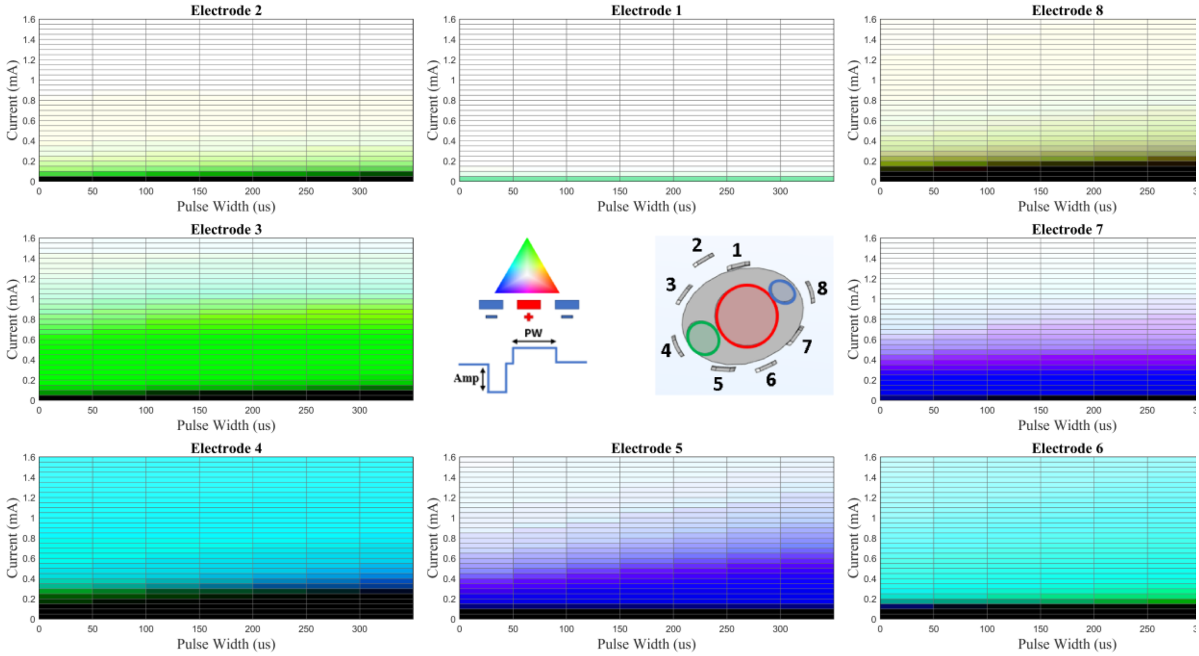


**Figure S6.** Nerve 1 – Config 3 (Simulation)

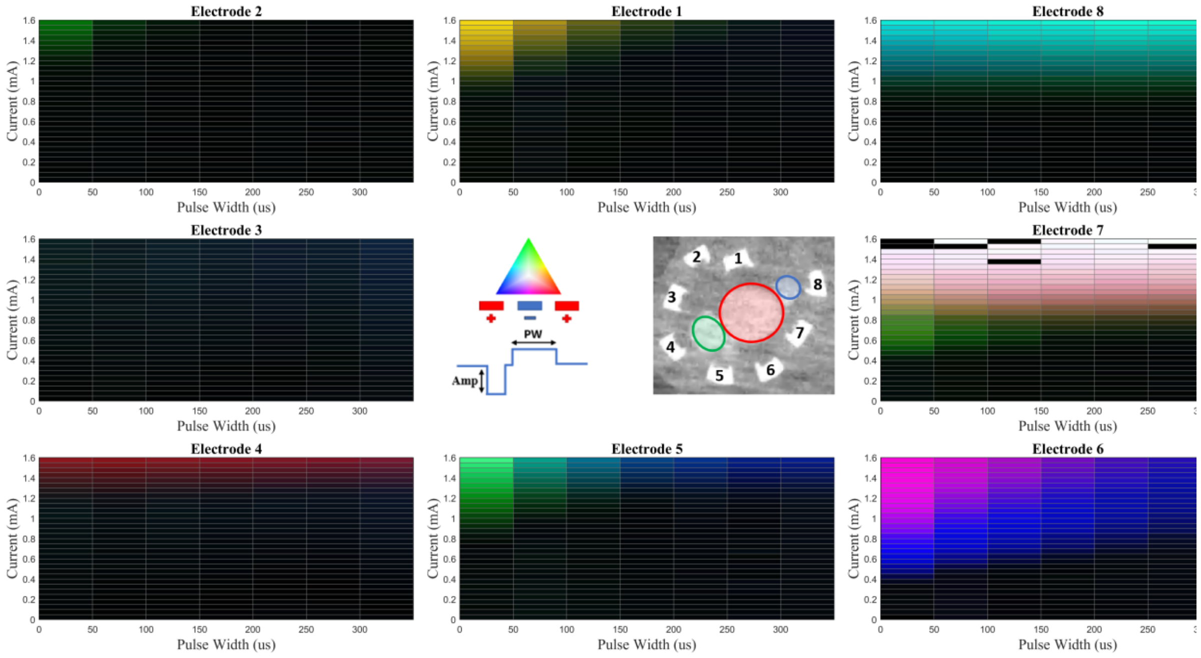


**Figure S7.** Nerve 1 – Config 4 (Ex Vivo)

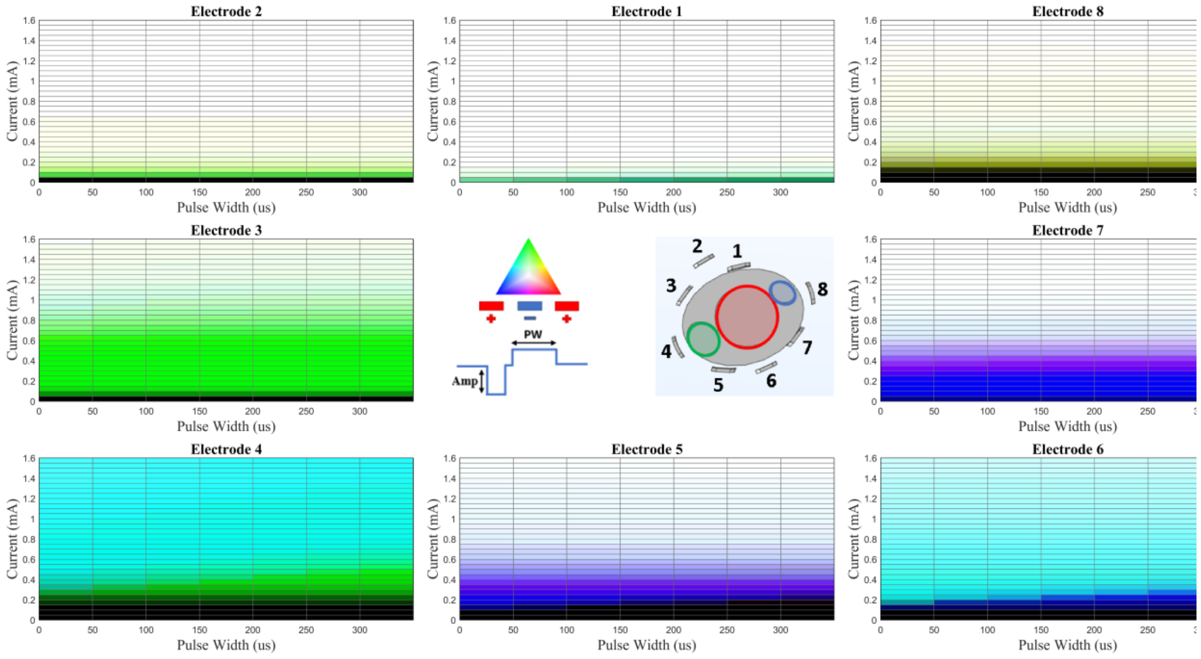


**Figure S8.** Nerve 1 – Config 4 (Simulation)

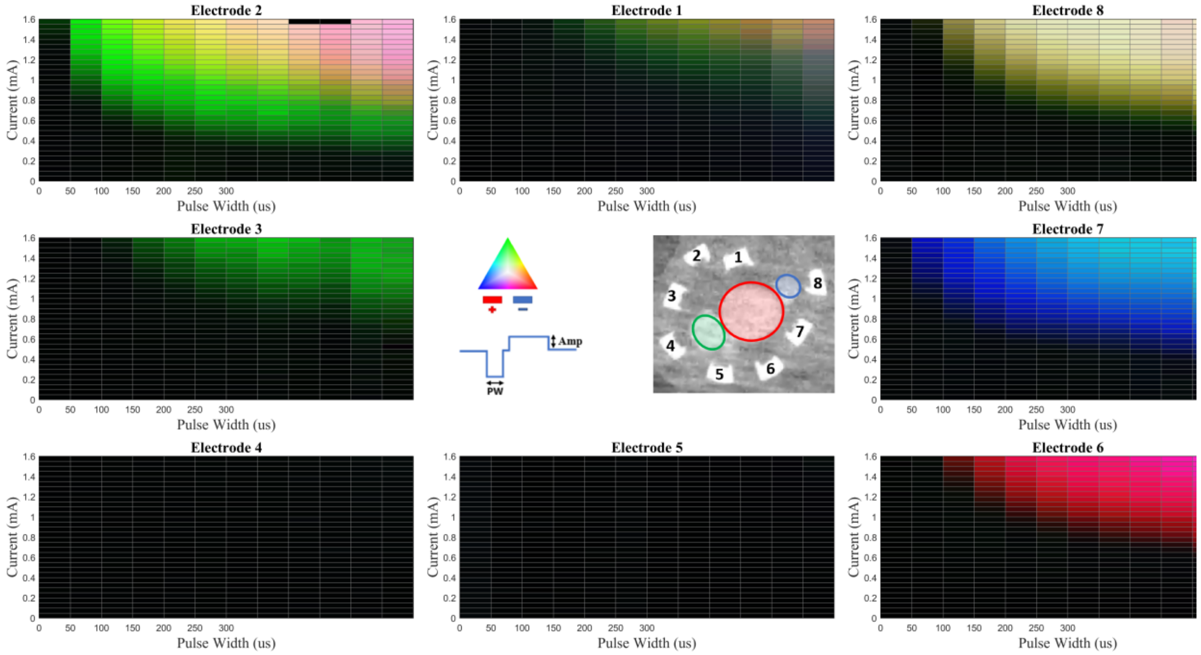


**Figure S9.** Nerve 1 – Config 5 (Ex Vivo)

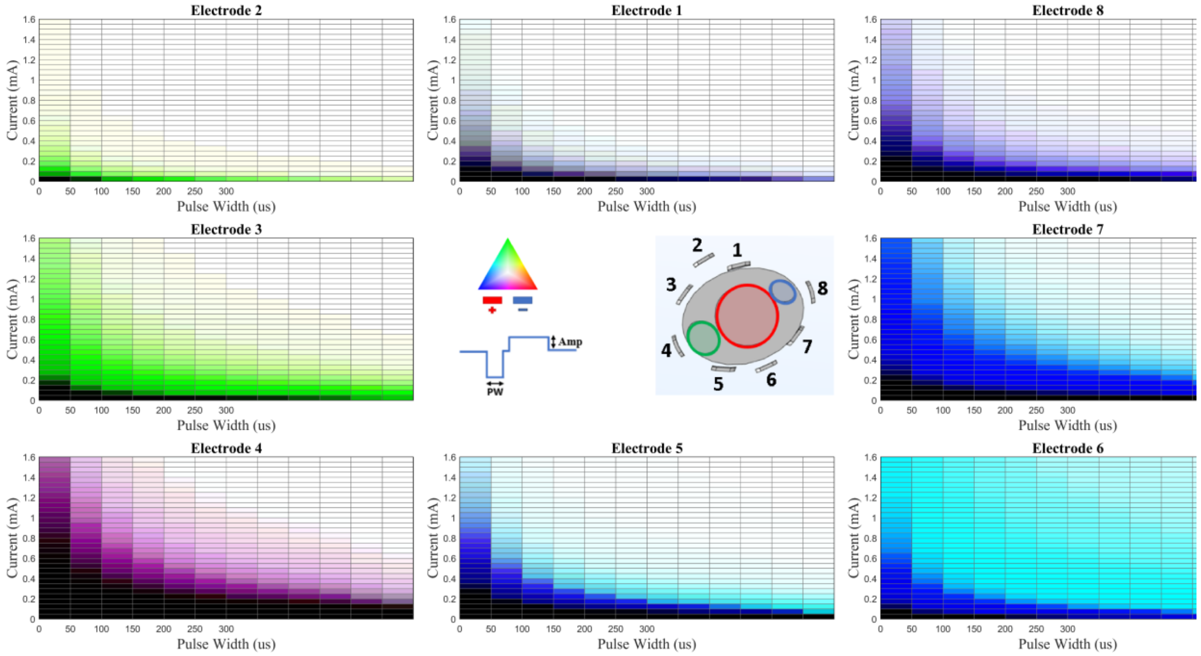


**Figure S10.** Nerve 1 – Config 5 (Simulation)

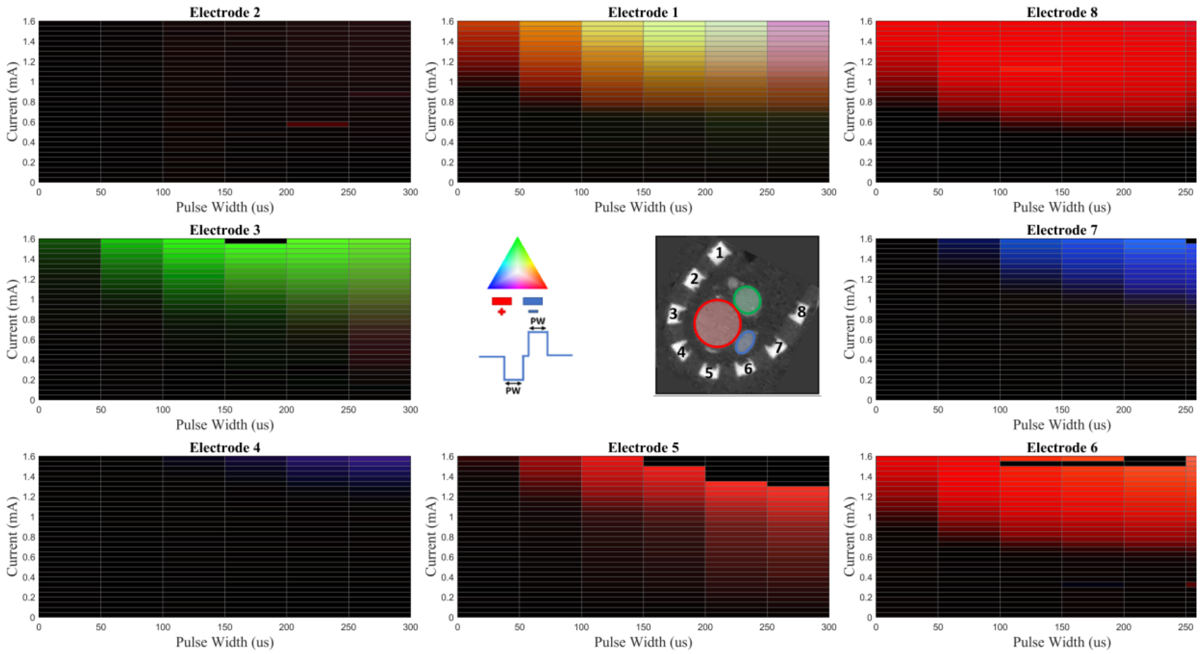


**Figure S11.** Nerve 2 – Config 1 (Ex Vivo)

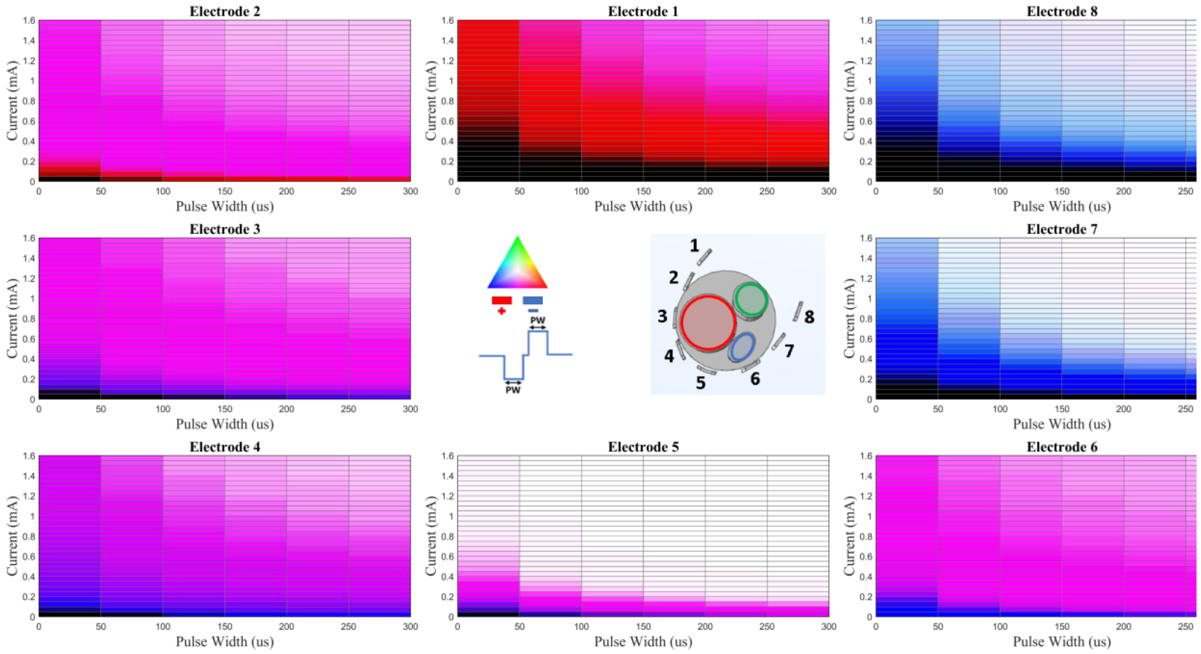


**Figure S12.** Nerve 2 – Config 1 (Simulation)

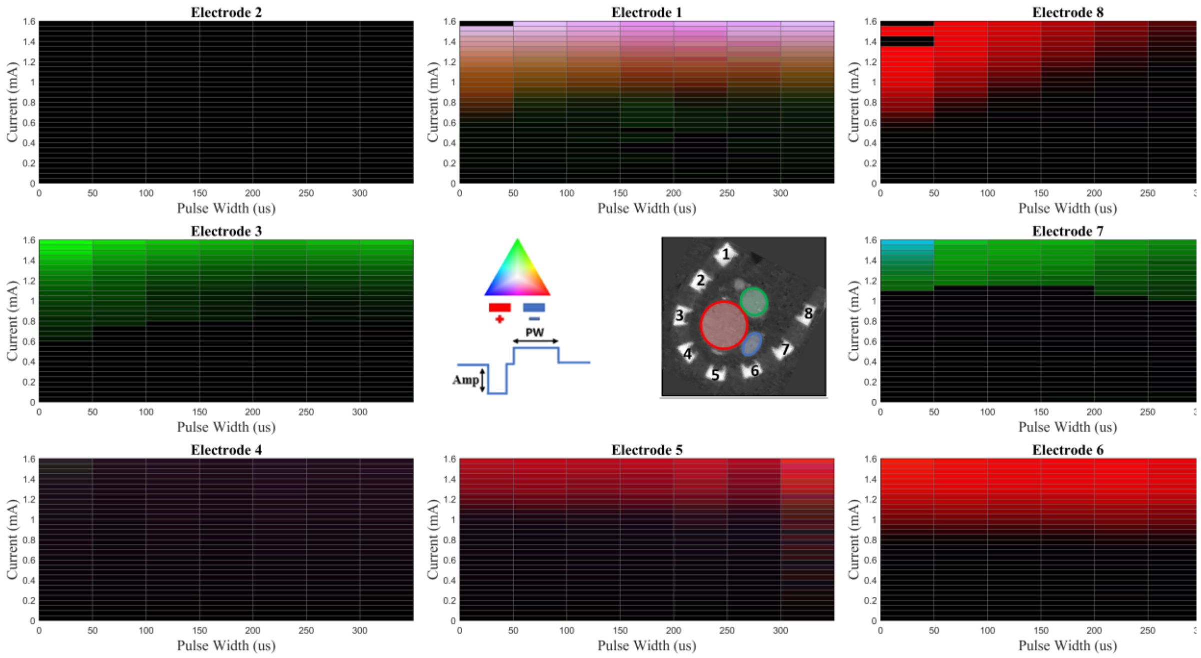


**Figure S13.** Nerve 2 – Config 2 (Ex Vivo)

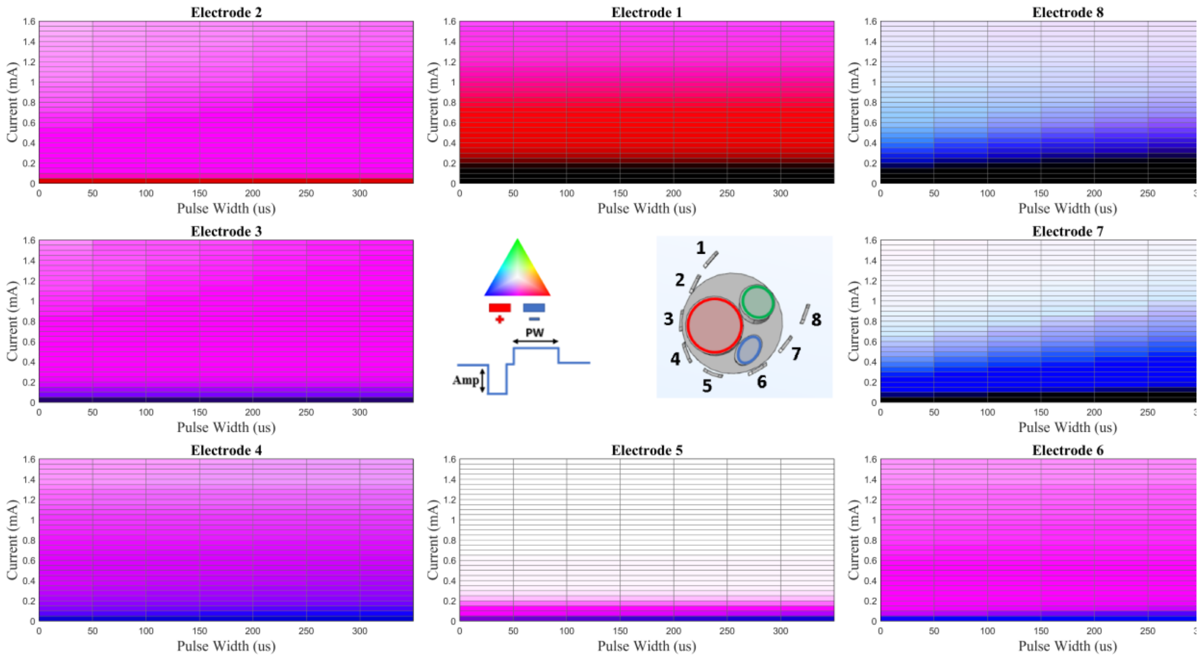


**Figure S14.** Nerve 2 – Config 2 (Simulation)

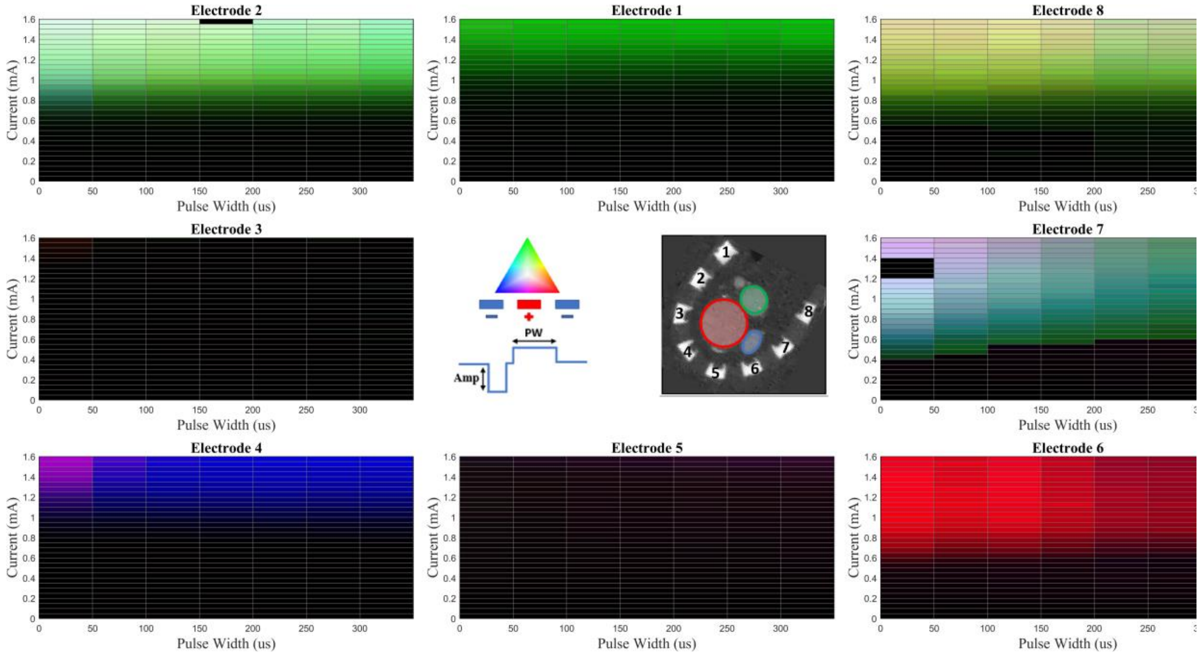


**Figure S15.** Nerve 2 – Config 3 (Ex Vivo)

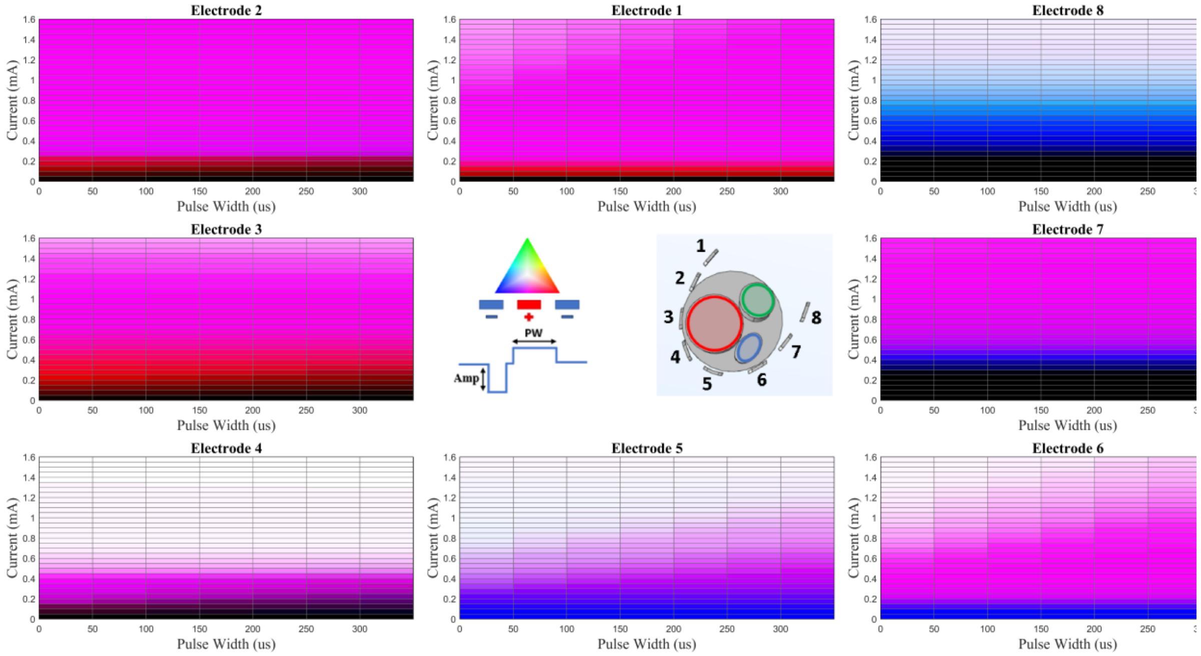


**Figure S16.** Nerve 2 – Config 3 (Simulation)

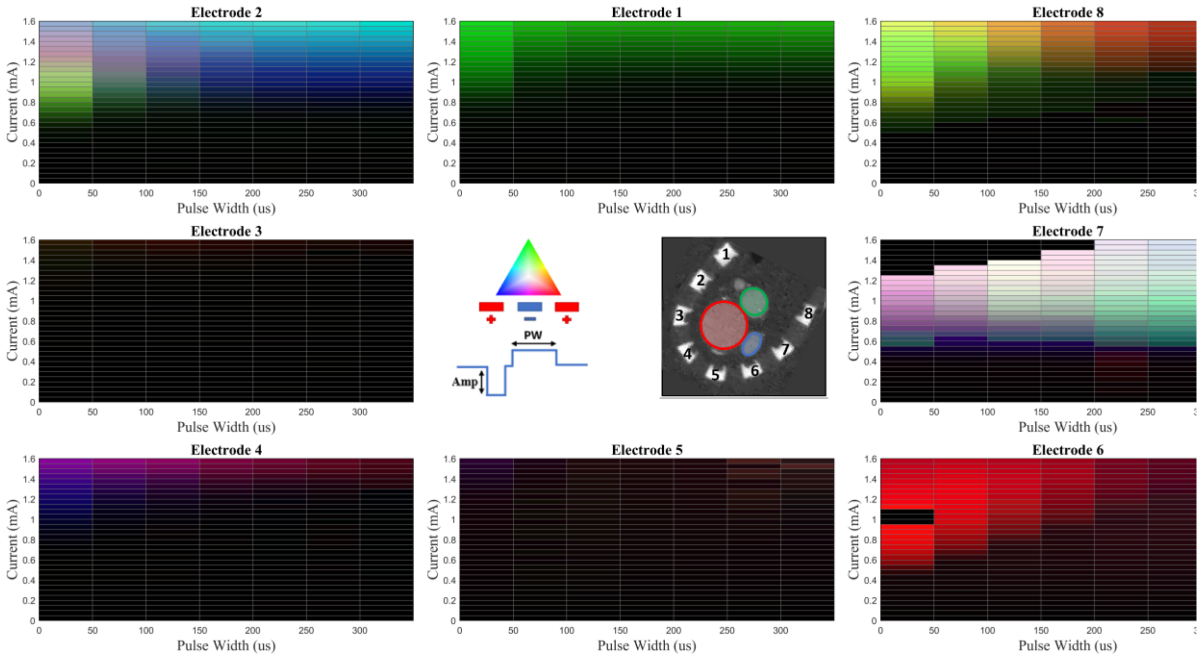


**Figure S17.** Nerve 2 – Config 4 (Ex Vivo)

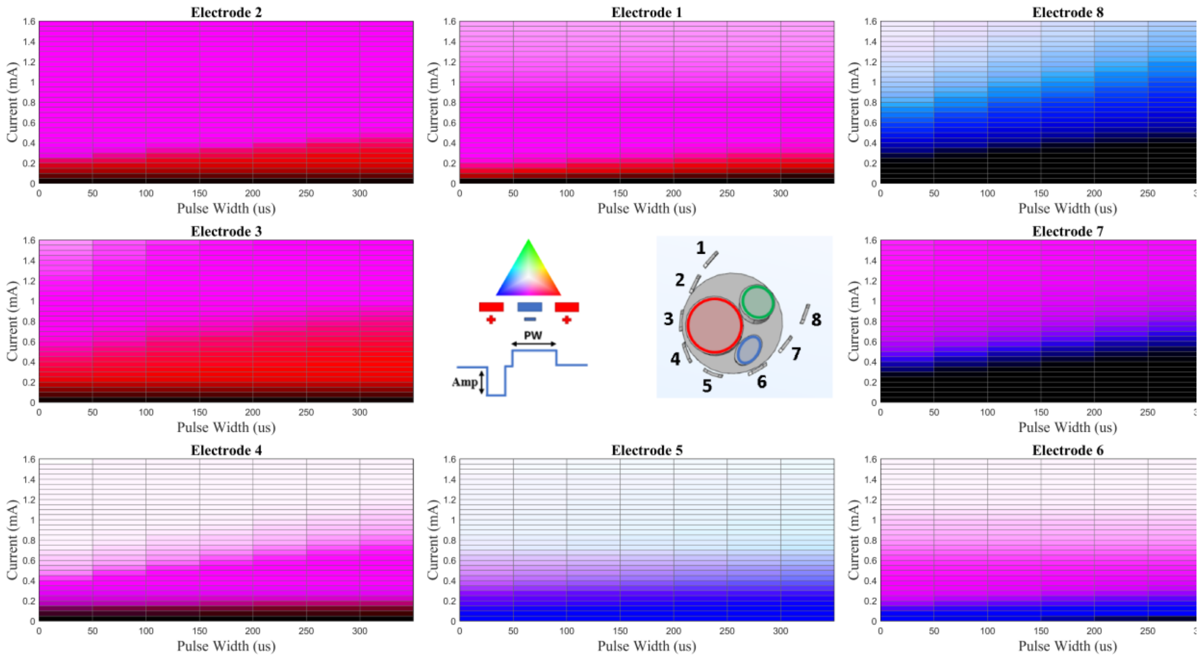


**Figure S18.** Nerve 2 – Config 4 (Simulation)

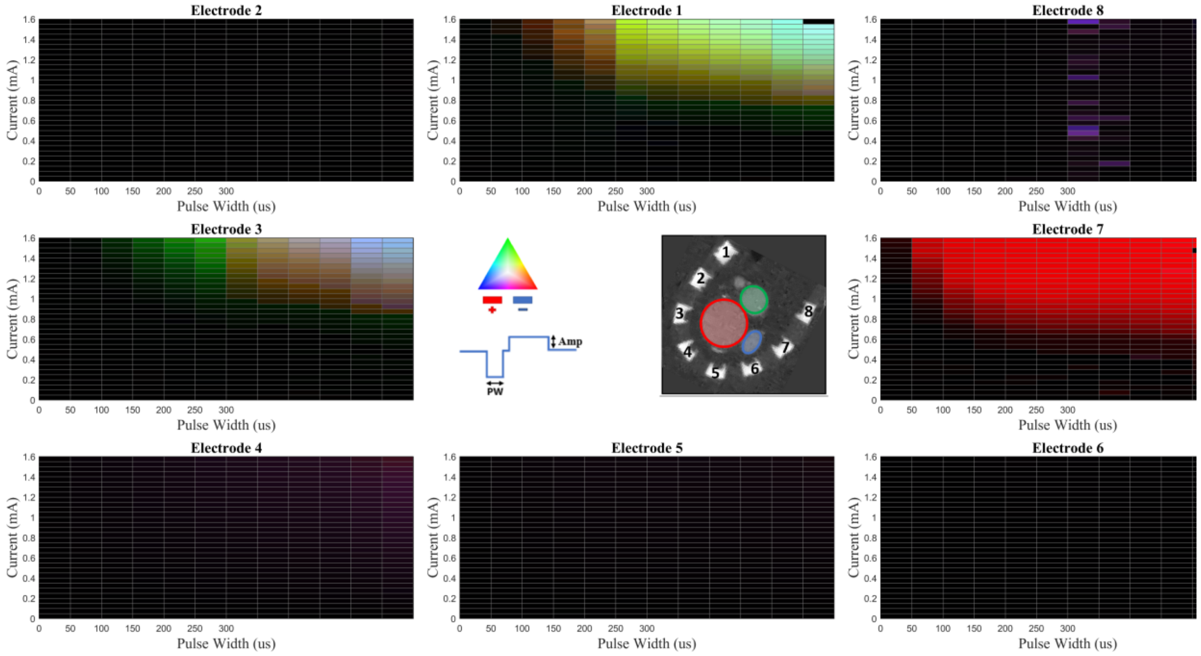


**Figure S19.** Nerve 2 – Config 5 (Ex Vivo)

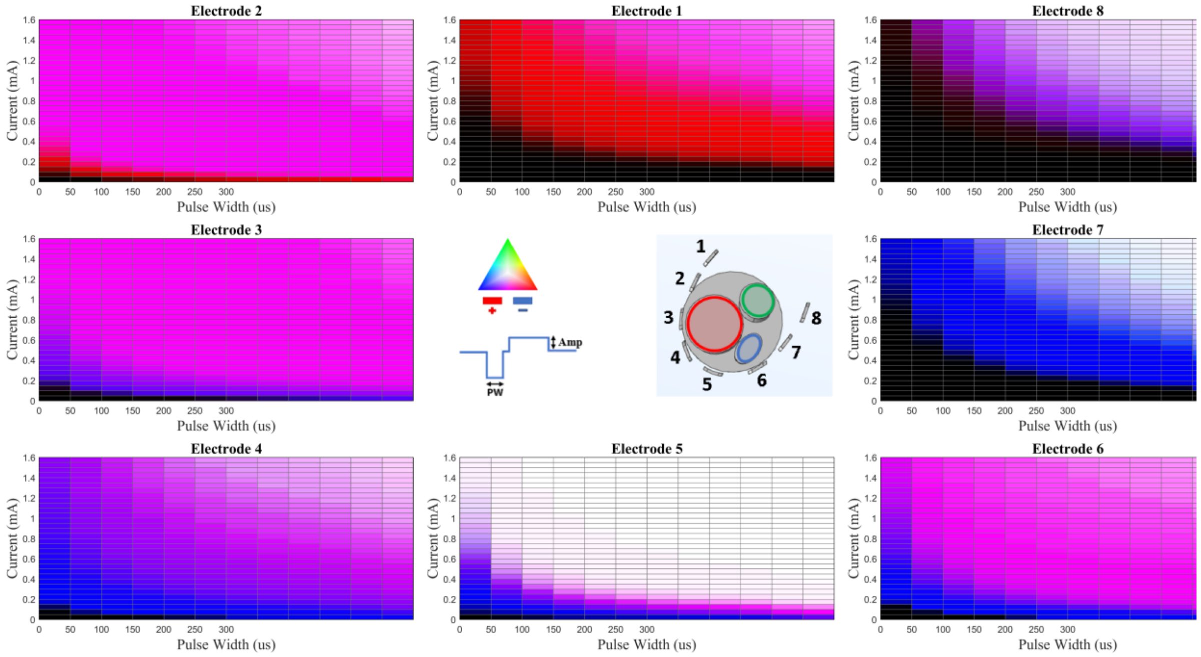


**Figure S20.** Nerve 2 – Config 5 (Simulation)

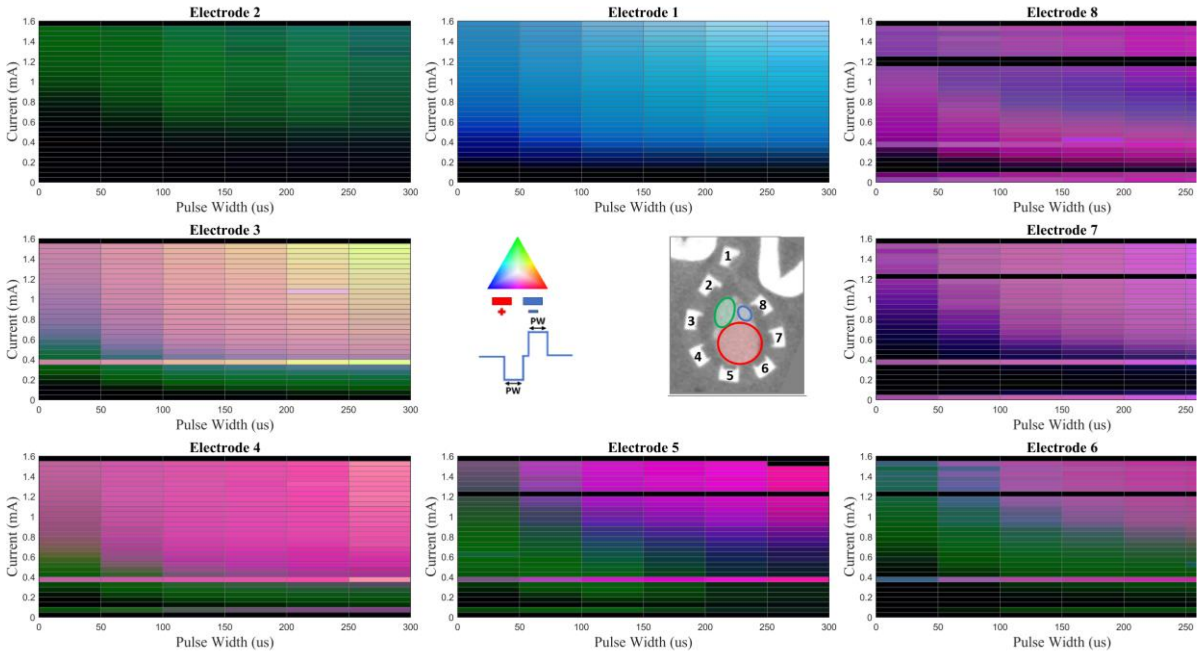


**Figure S21.** Nerve 3 – Config 1 (Ex Vivo)

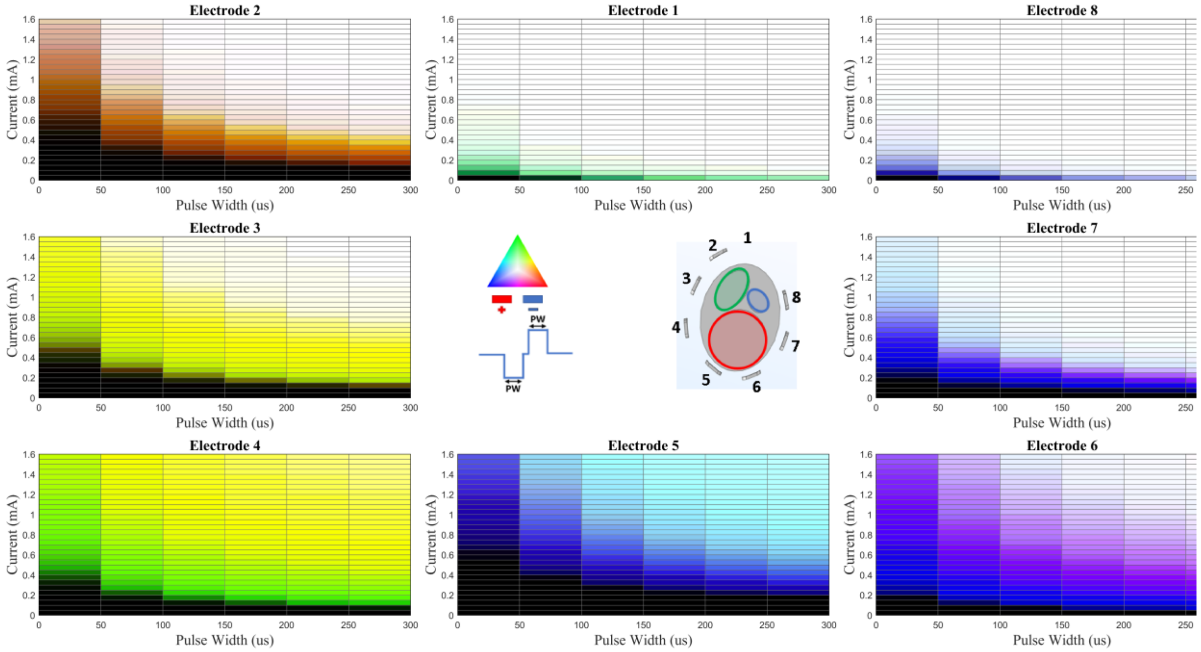


**Figure S22.** Nerve 3 – Config 1 (Simulation)

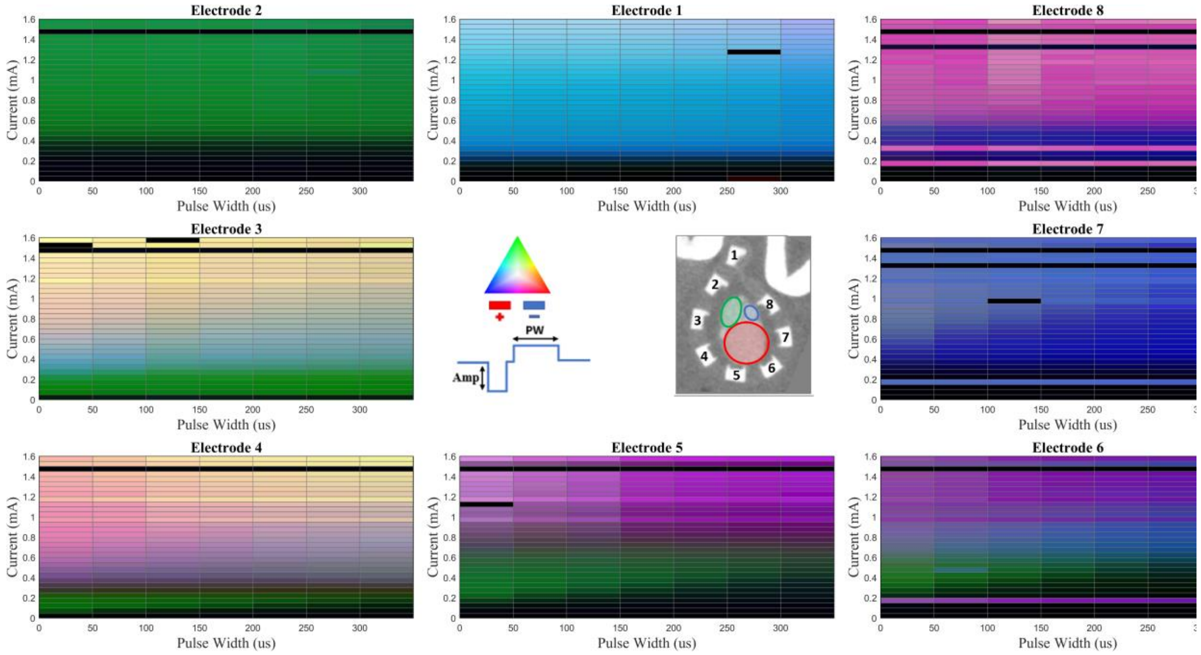


**Figure S23.** Nerve 3 – Config 2 (Ex Vivo)

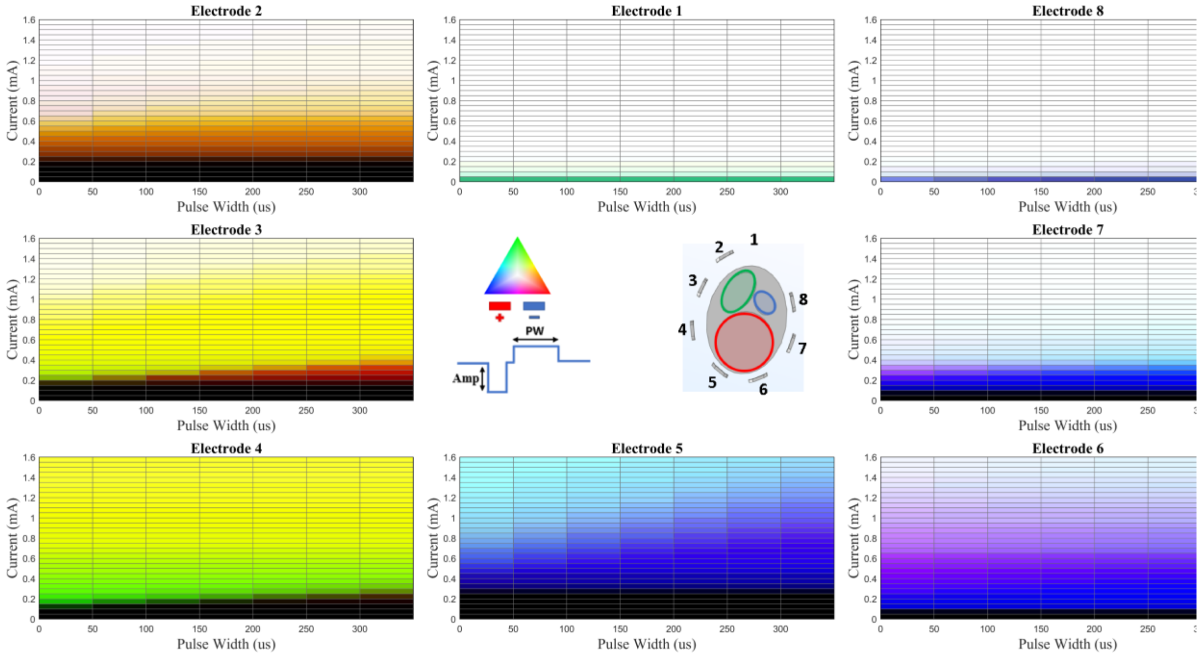


**Figure S24.** Nerve 3 – Config 2 (Simulation)

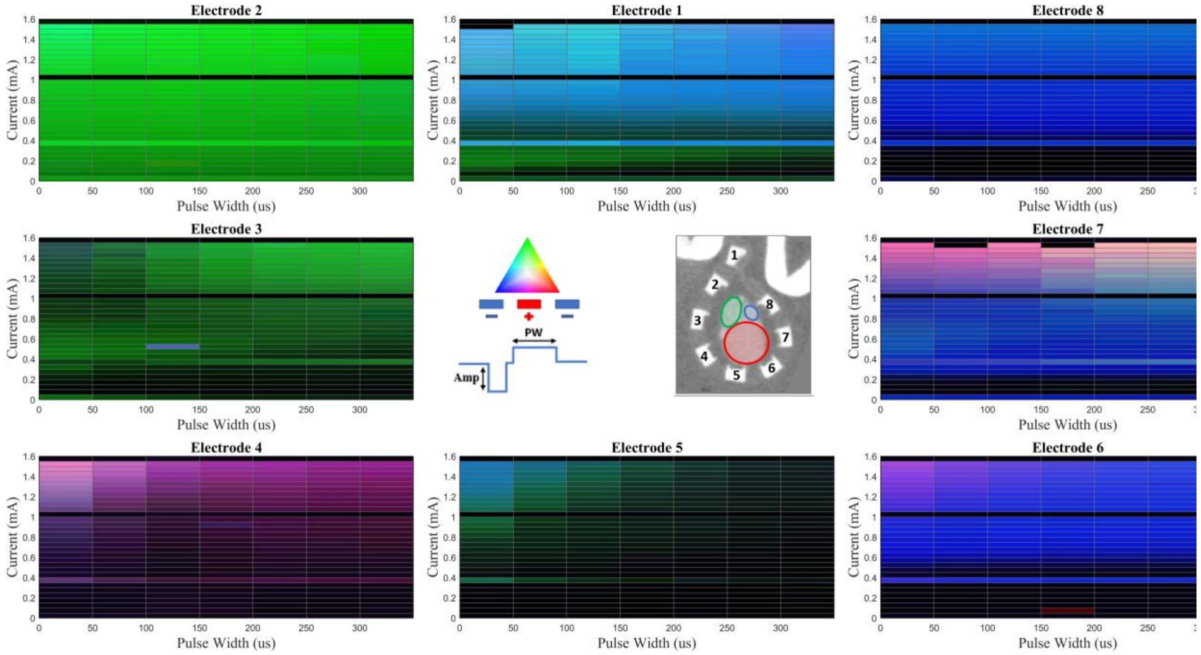


**Figure S25.** Nerve 3 – Config 3 (Ex Vivo)

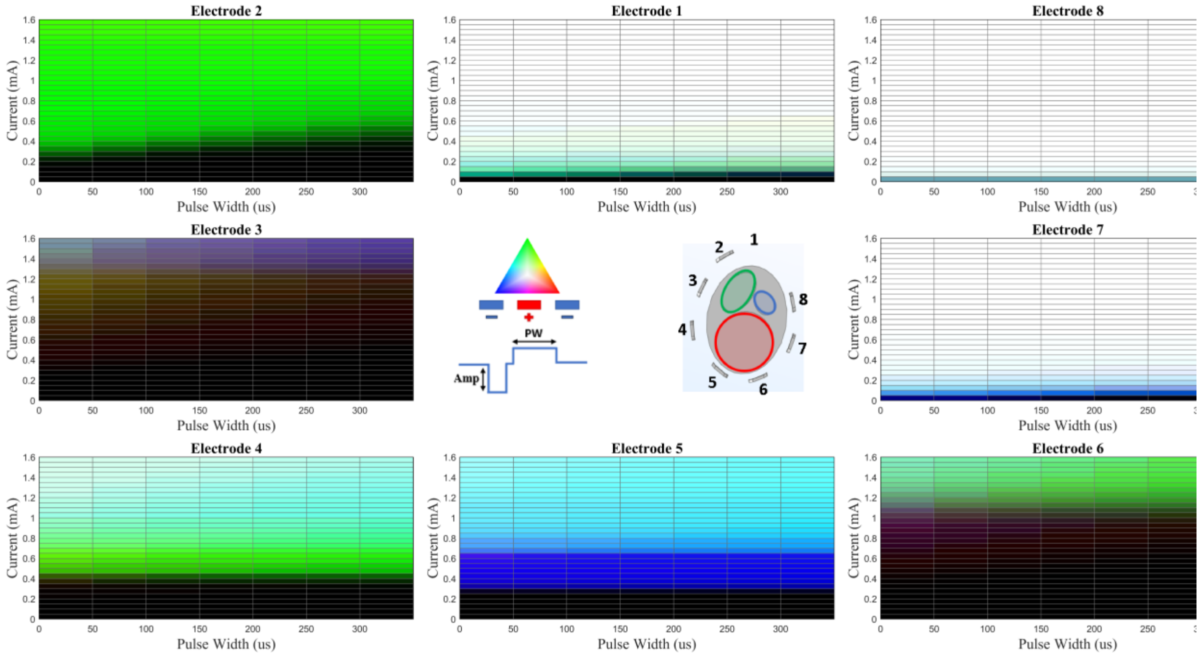


**Figure S26.** Nerve 3 – Config 3 (Simulation)

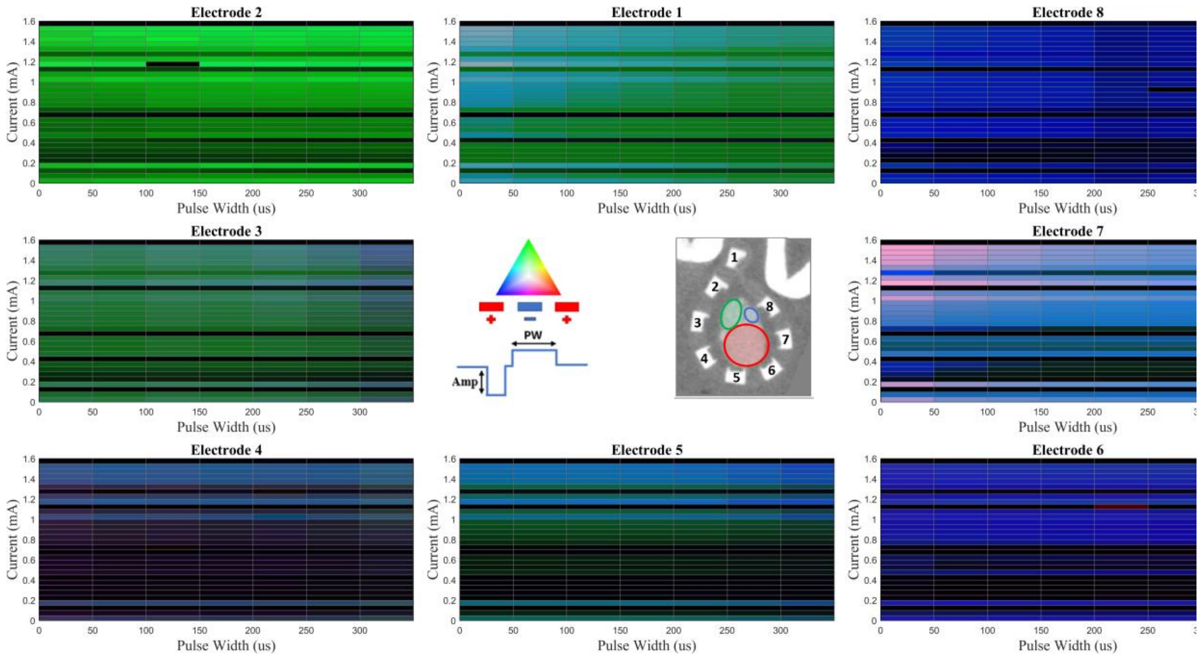


**Figure S27.** Nerve 3 – Config 4 (Ex Vivo)

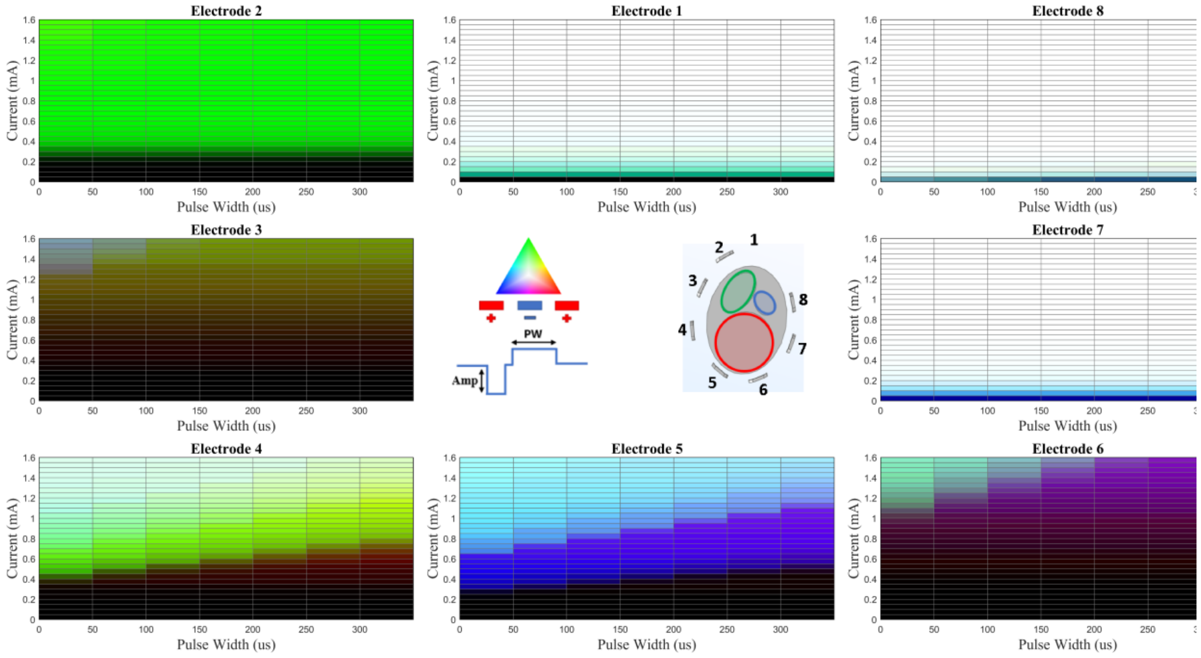


**Figure S28.** Nerve 3 – Config 4 (Simulation)

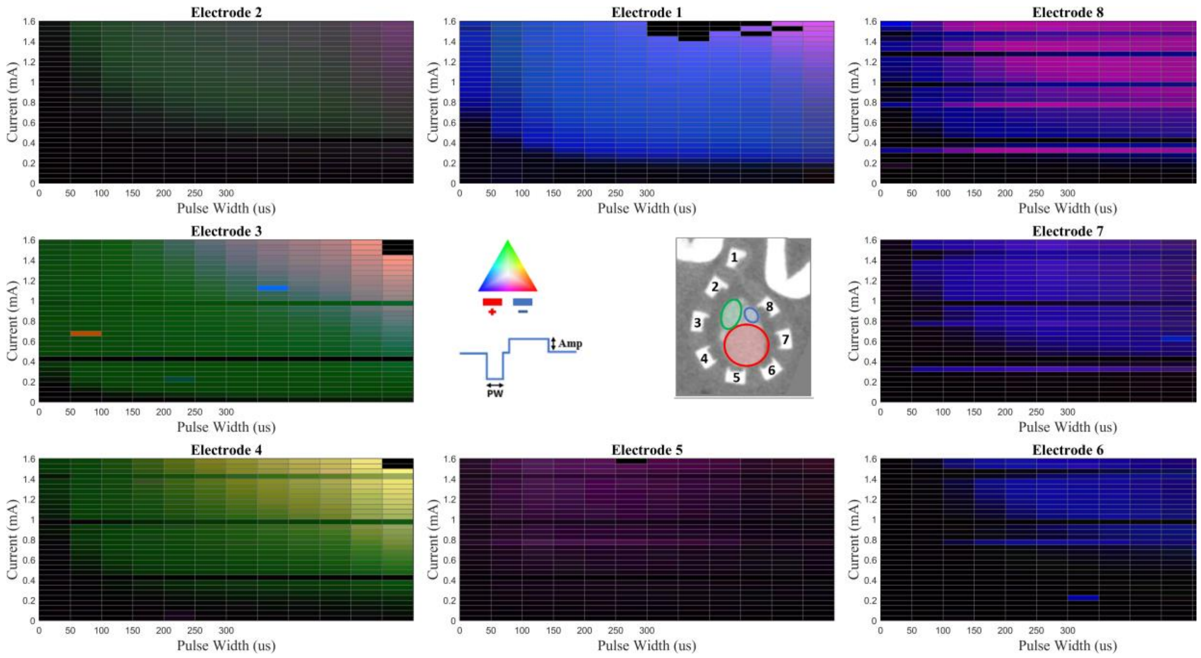


**Figure S29.** Nerve 3 – Config 5 (Ex Vivo)

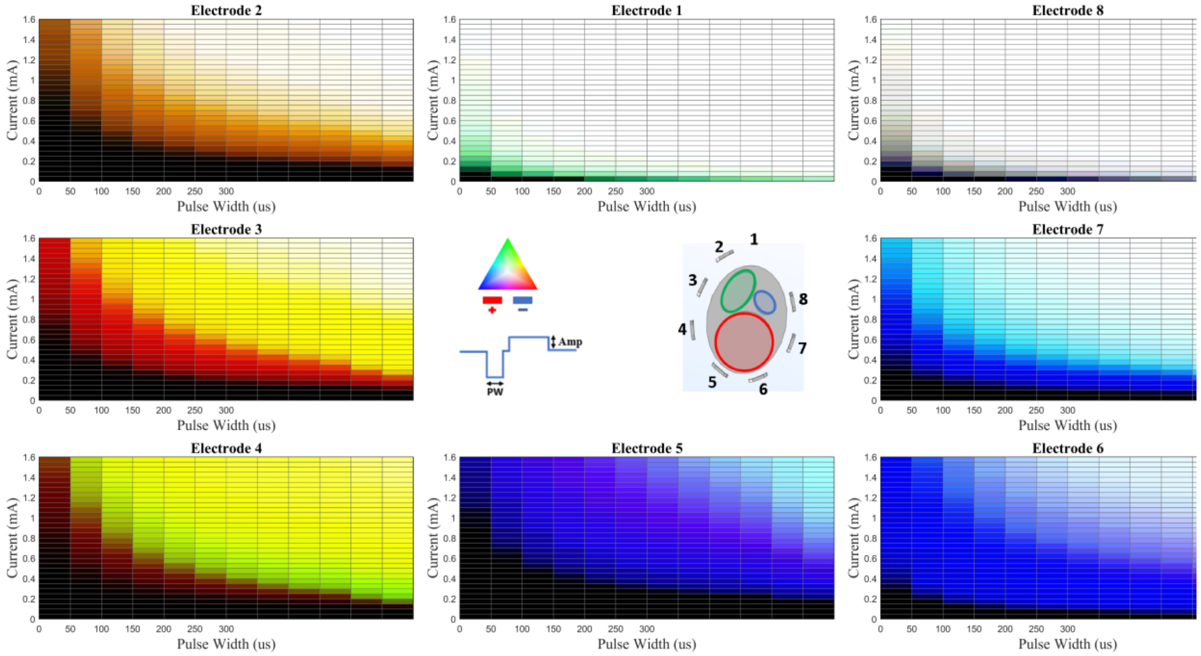


**Figure S30.** Nerve 3 – Config 5 (Simulation)

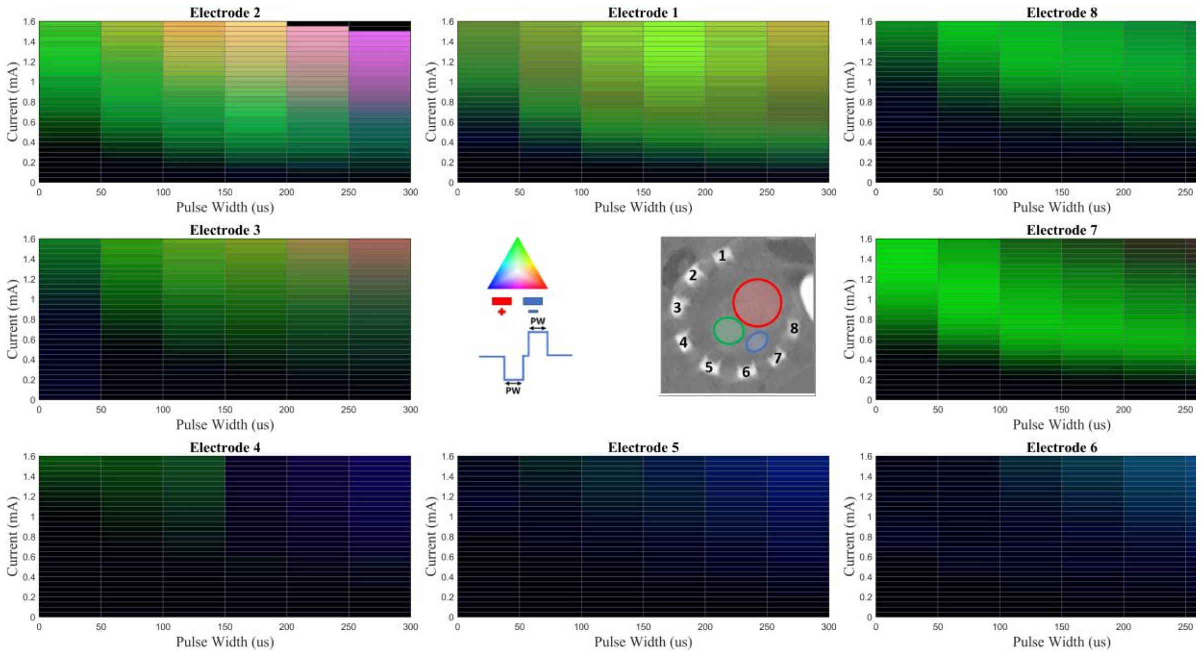


**Figure S31.** Nerve 4 – Config 1 (Ex Vivo)

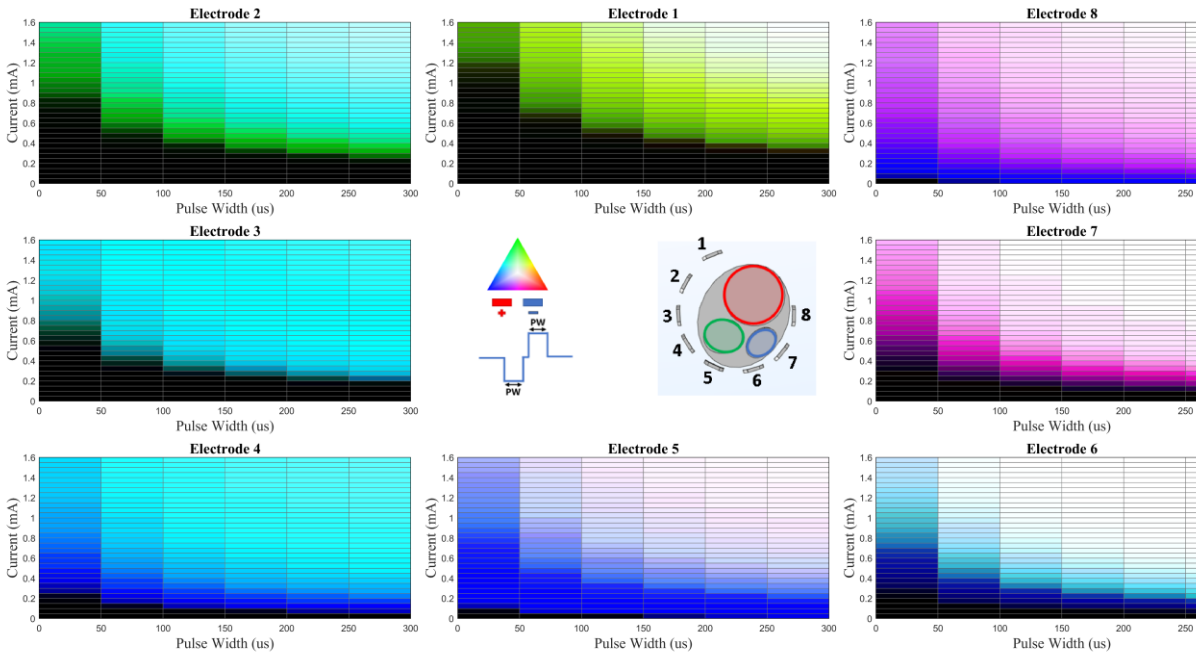


**Figure S32.** Nerve 4 – Config 1 (Simulation)

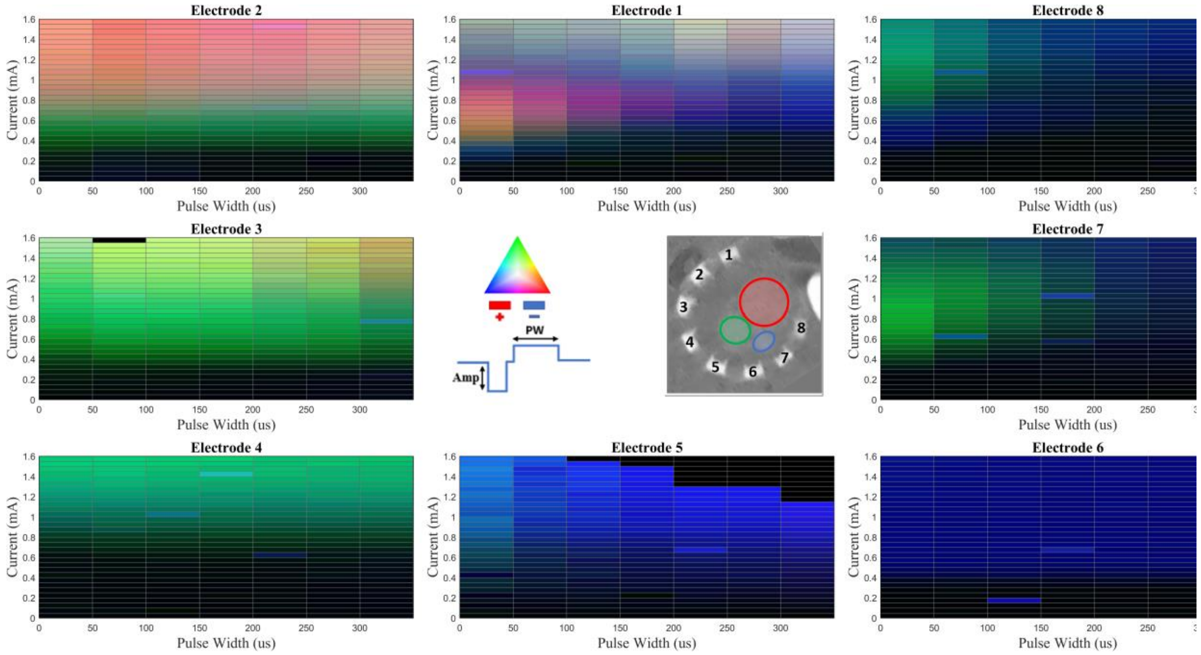

**Figure S33.** Nerve 4 – Config 2 (Ex Vivo)

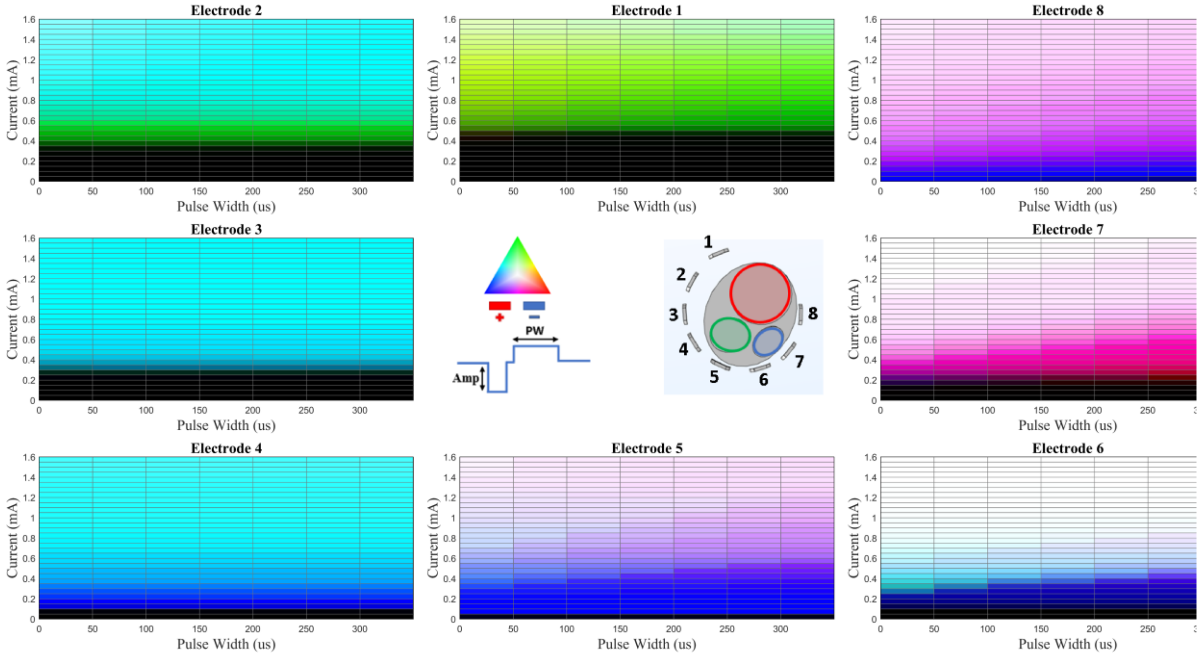

**Figure S34.** Nerve 4 – Config 2 (Simulation)

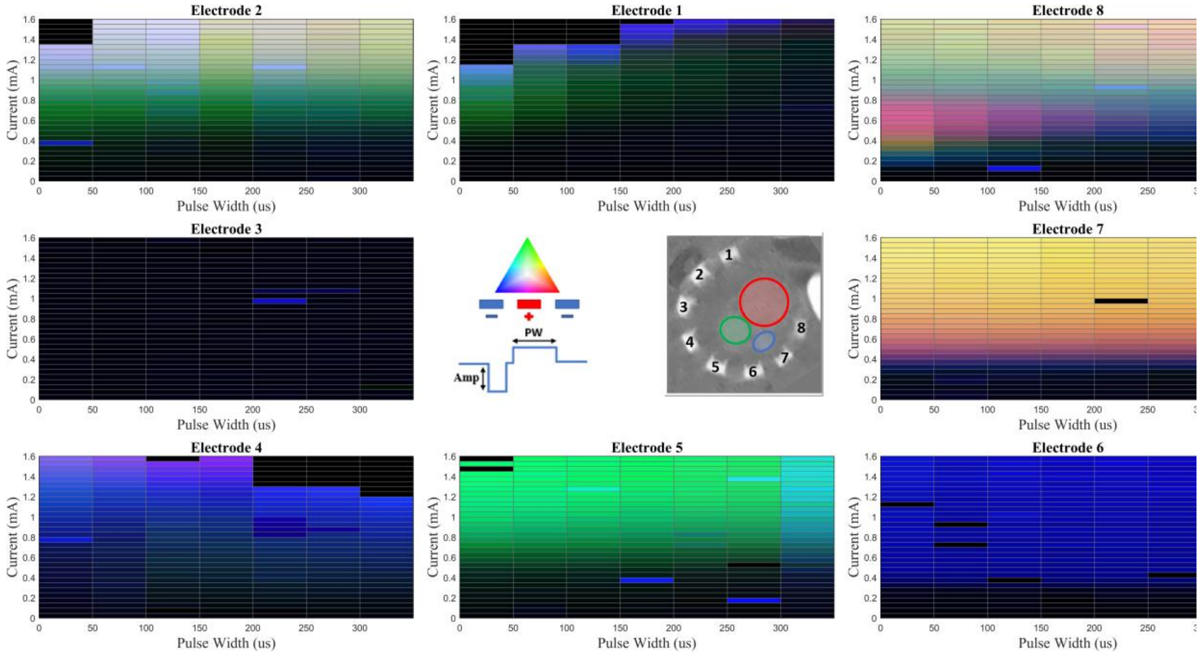


**Figure S35.** Nerve 4 – Config 3 (Ex Vivo)

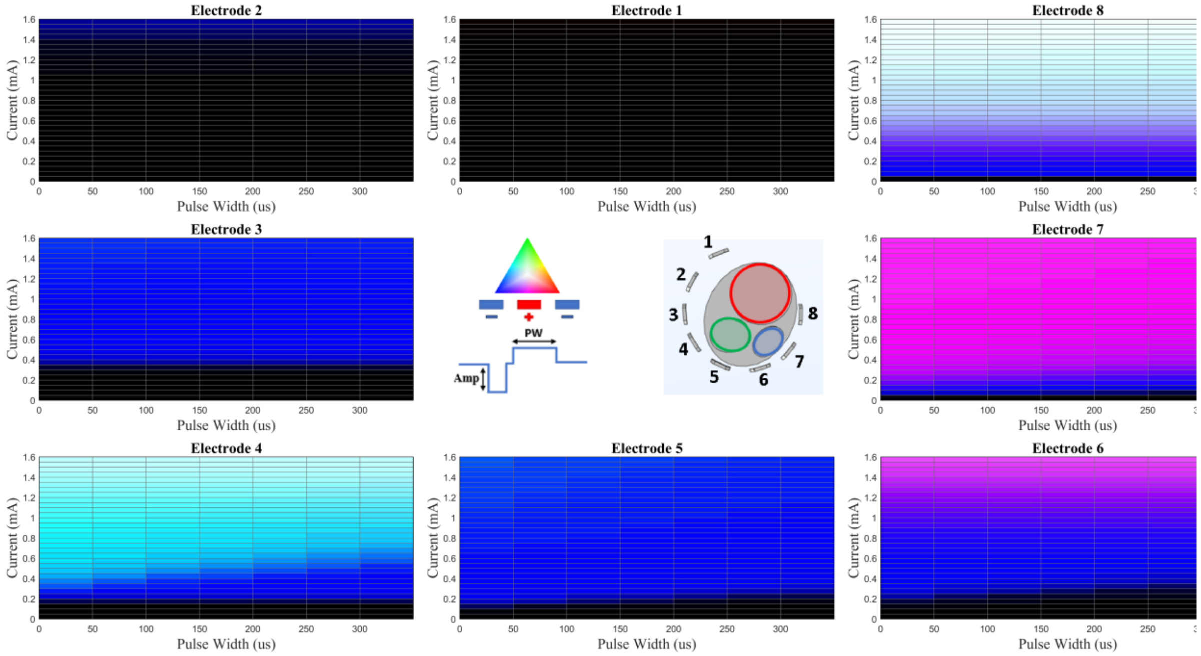


**Figure S36.** Nerve 4 – Config 3 (Simulation)

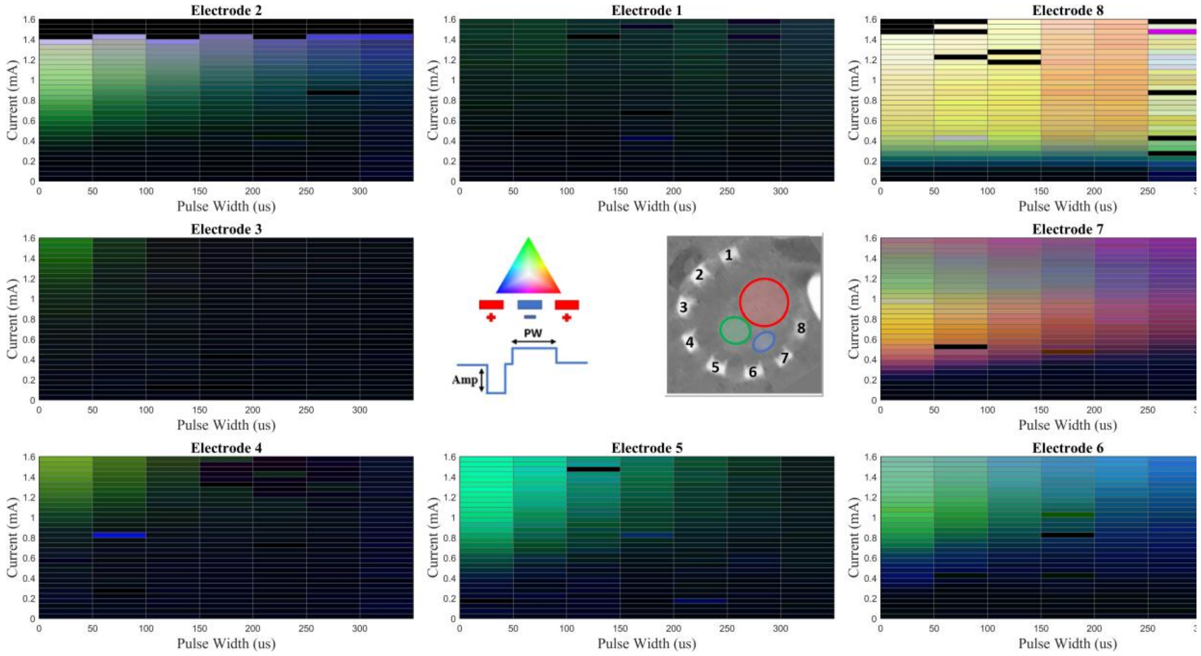


**Figure S37.** Nerve 4 – Config 4 (Ex Vivo)

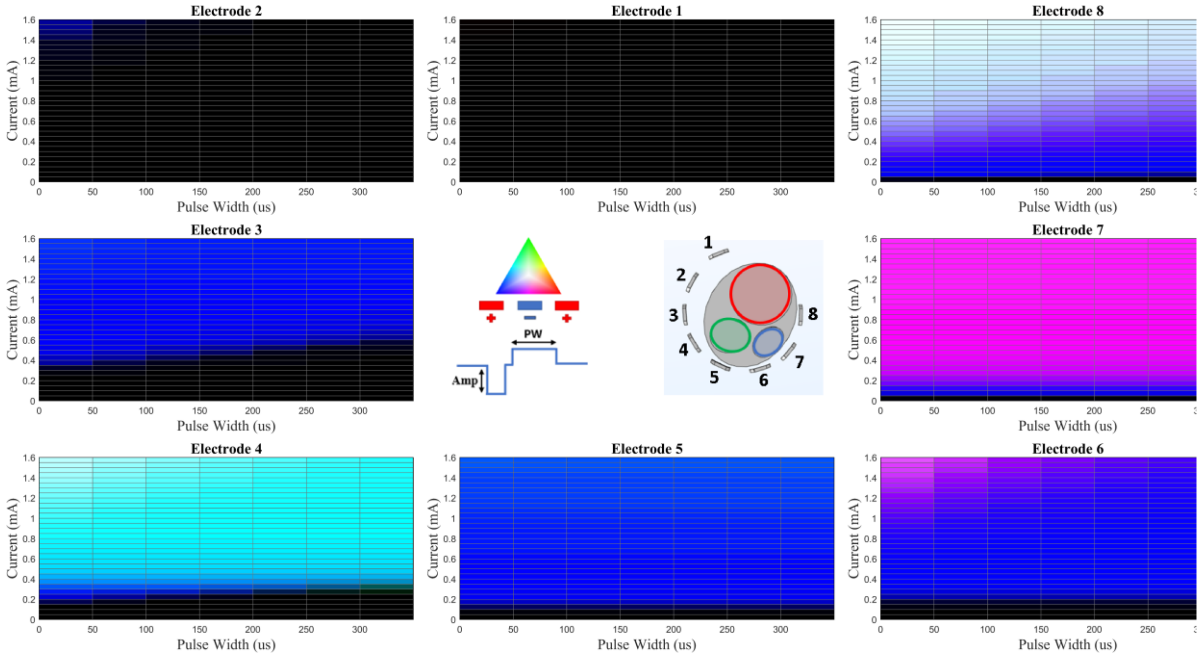


**Figure S38.** Nerve 4 – Config 4 (Simulation)

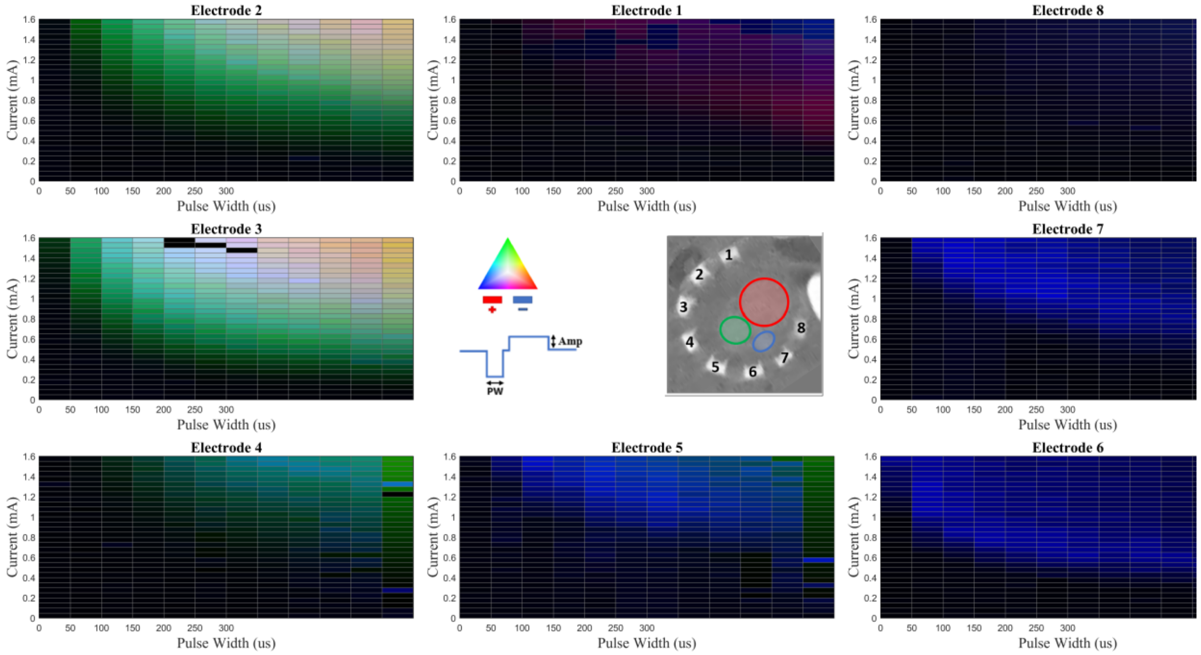

**Figure S39.** Nerve 4 – Config 5 (Ex Vivo)

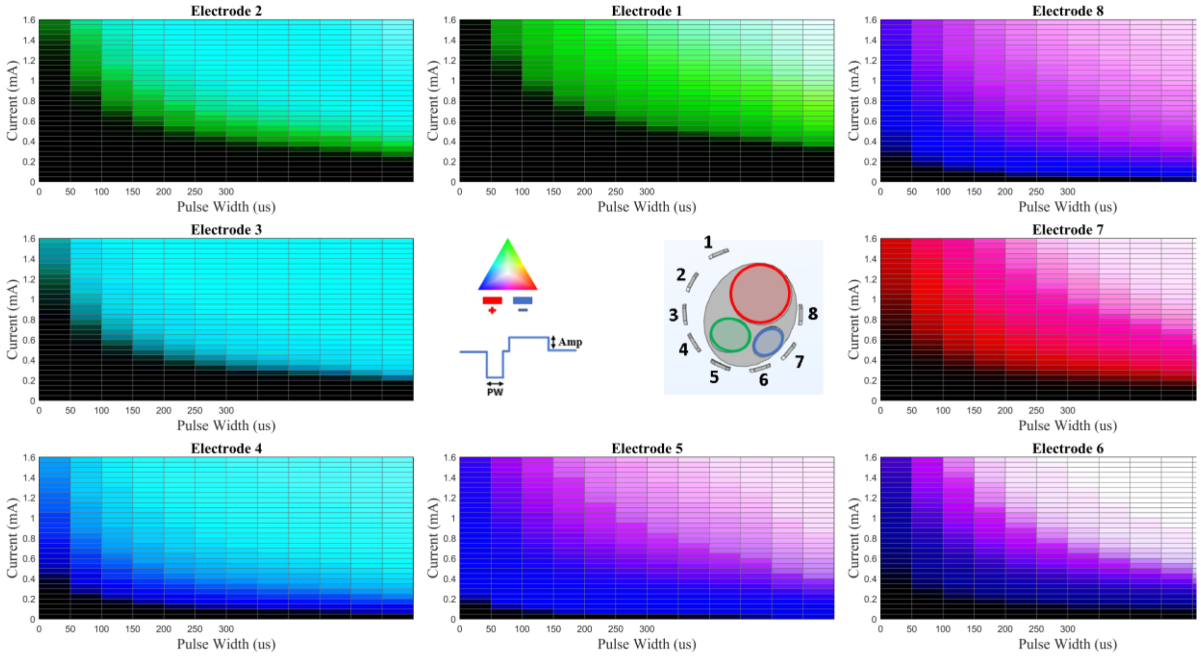

**Figure S40.** Nerve 4 – Config 5 (Simulation)

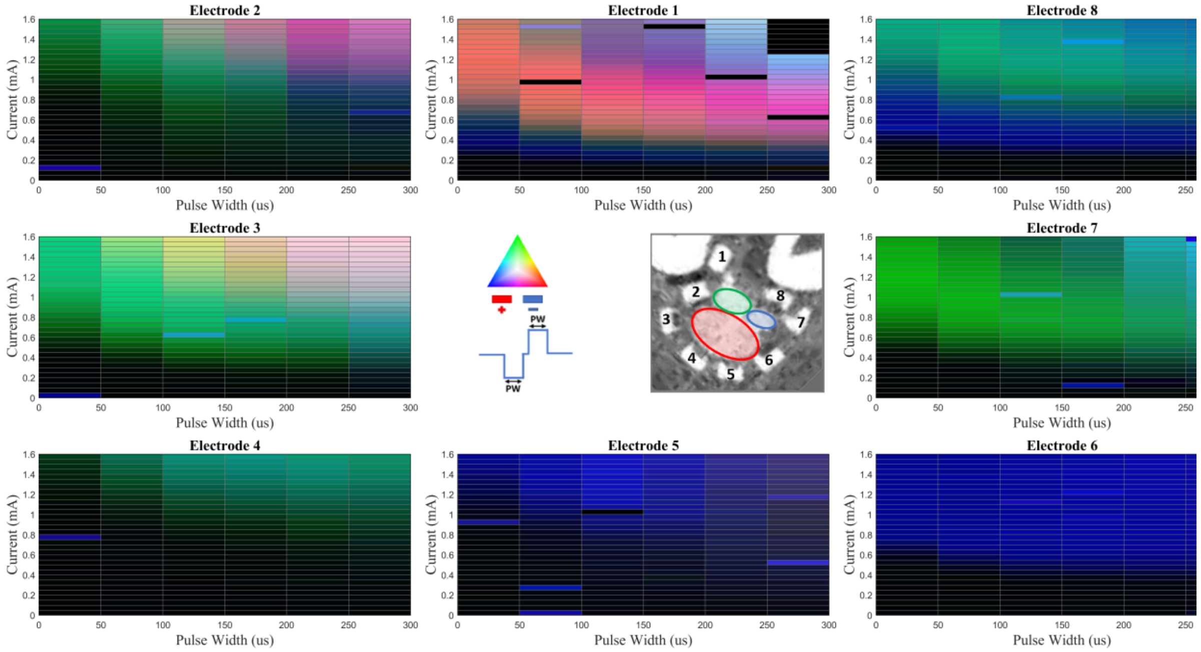


**Figure S41.** Nerve 5 – Config 1 (Ex Vivo)

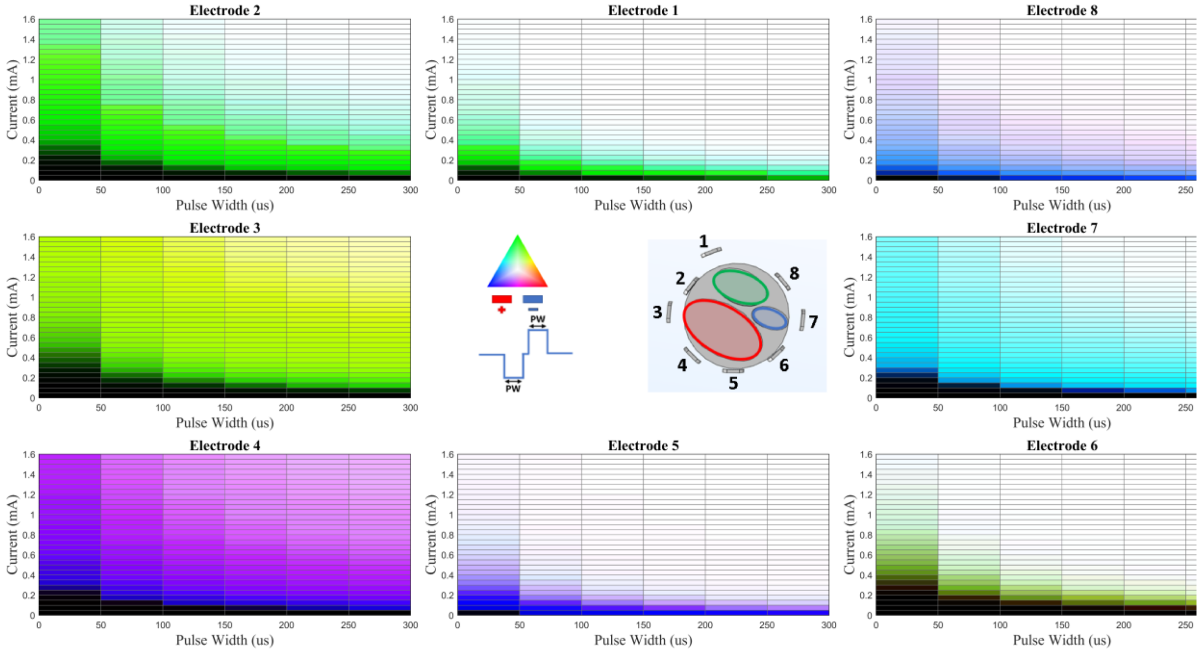


**Figure S42.** Nerve 5 – Config 1 (Simulation)

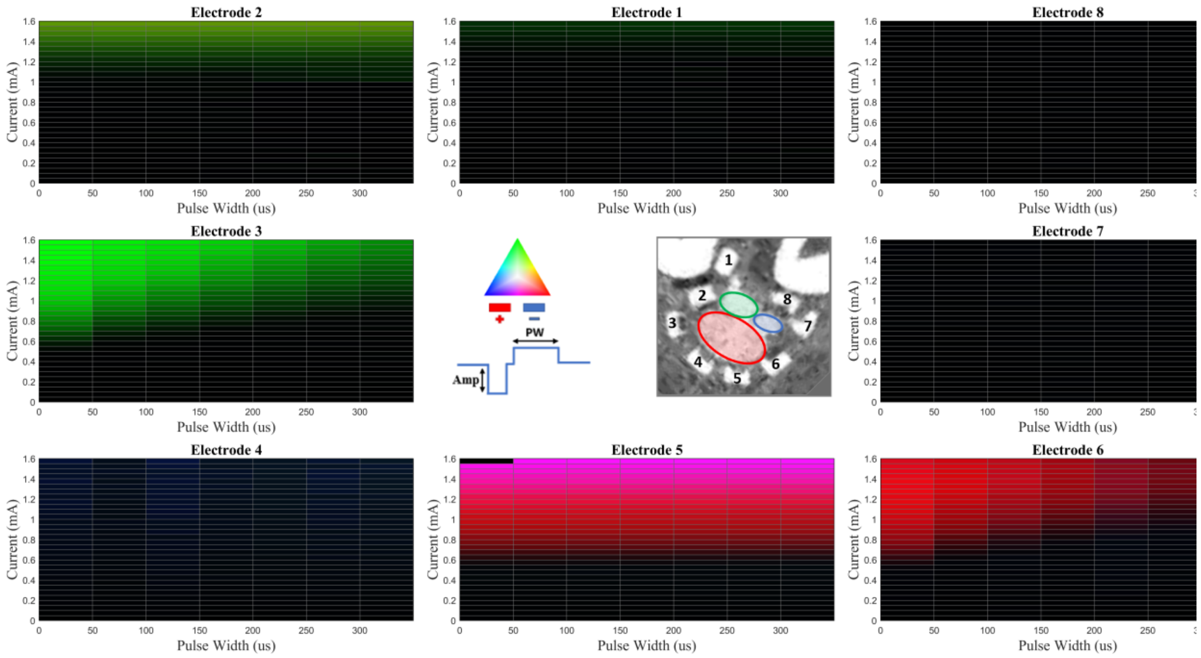


**Figure S43.** Nerve 5 – Config 2 (Ex Vivo)

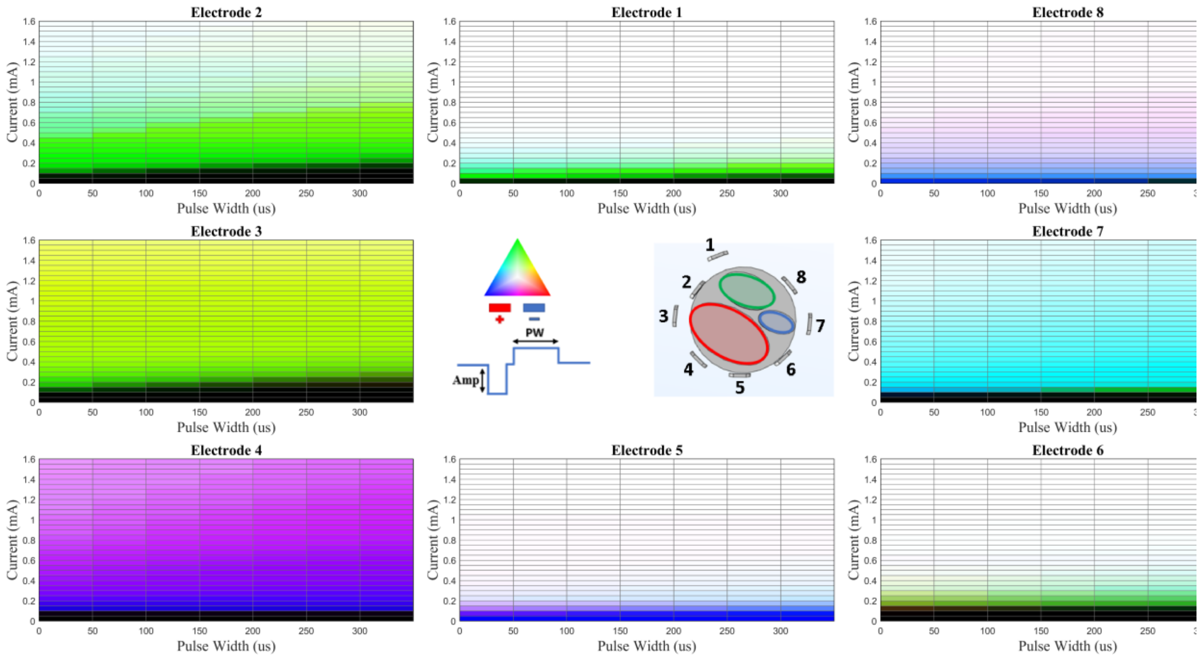


**Figure S44.** Nerve 5 – Config 2 (Simulation)

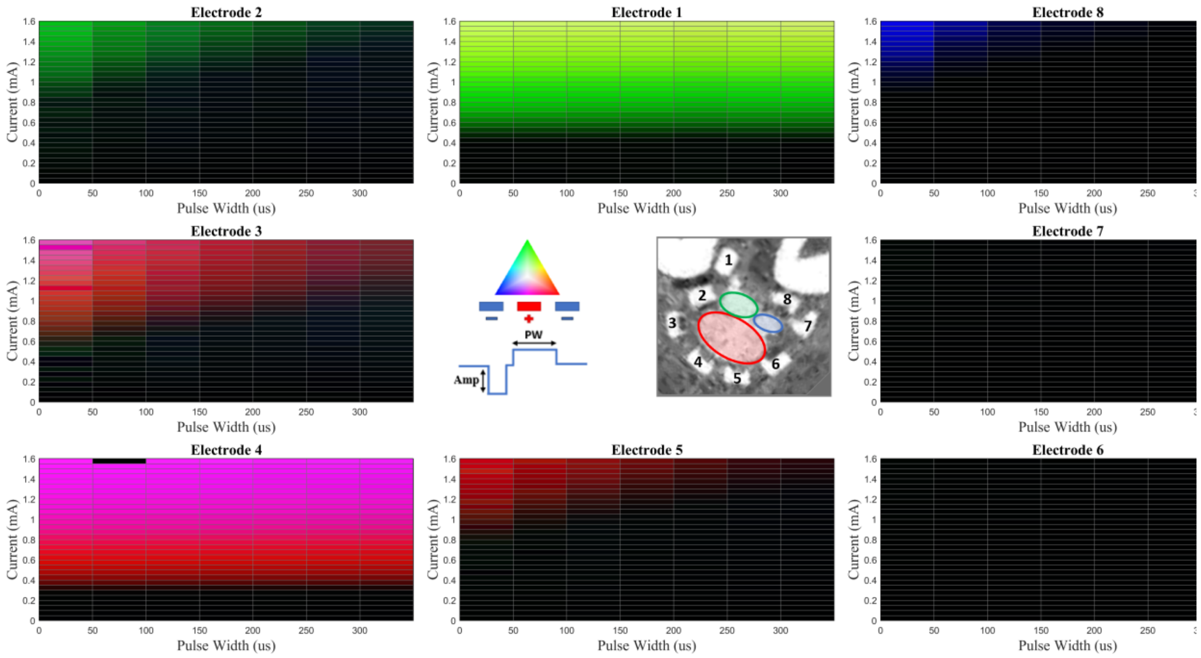


**Figure S45.** Nerve 5 – Config 3 (Ex Vivo)

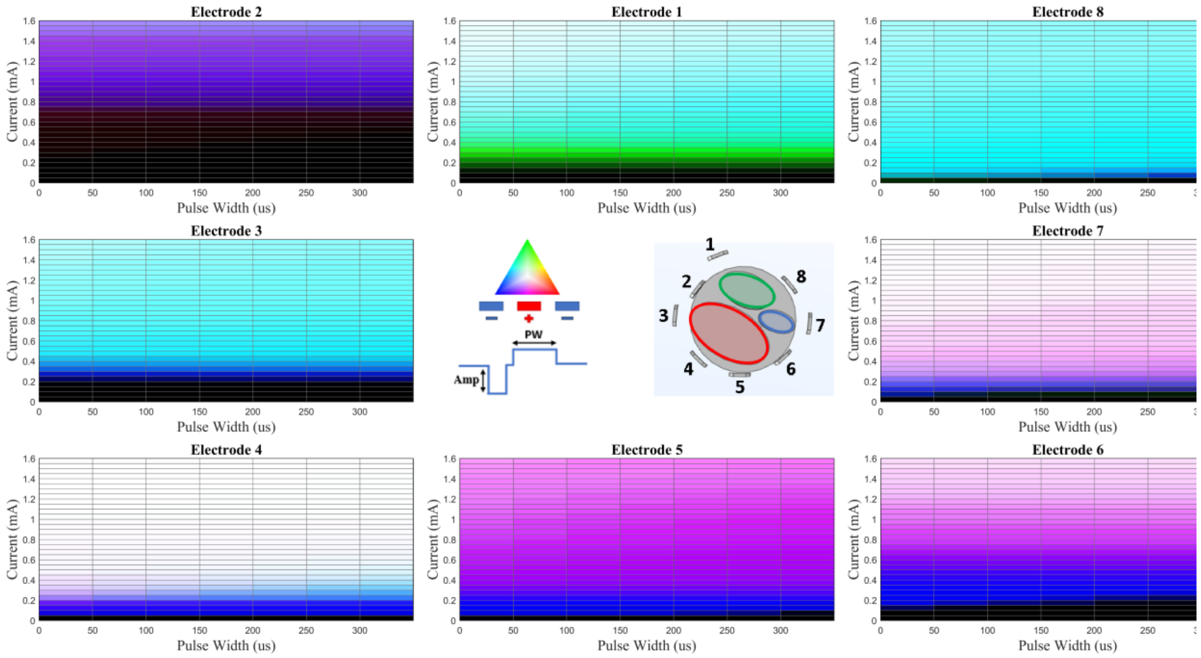


**Figure S46.** Nerve 5 – Config 3 (Simulation)

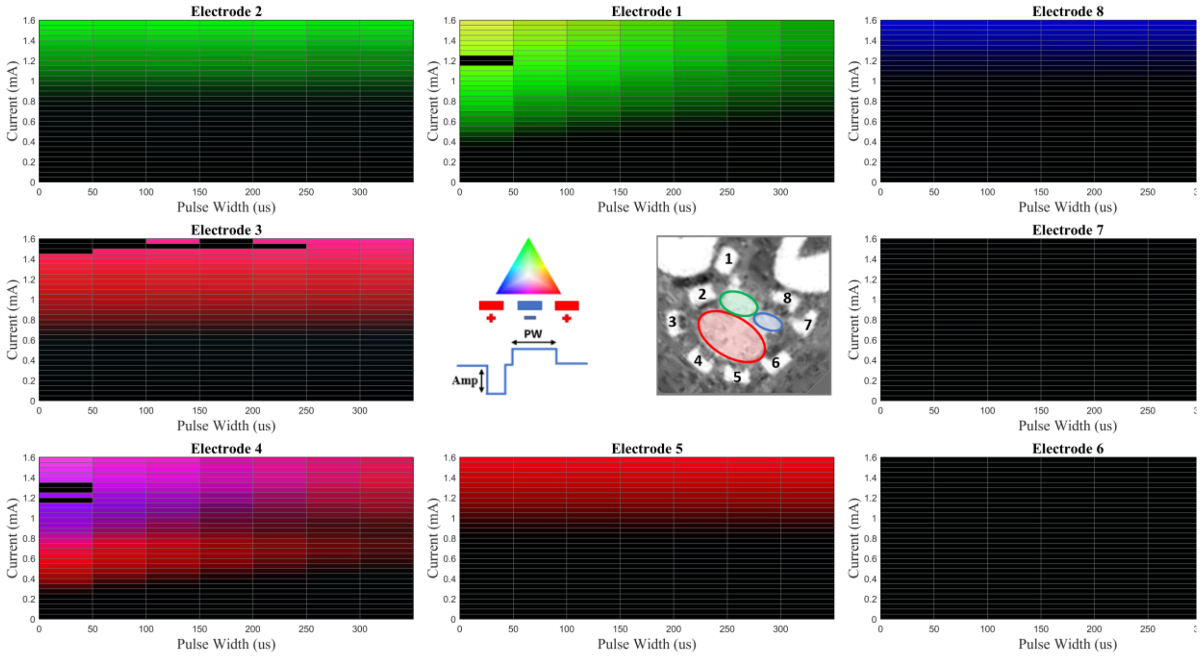


**Figure S47.** Nerve 5 – Config 4 (Ex Vivo)

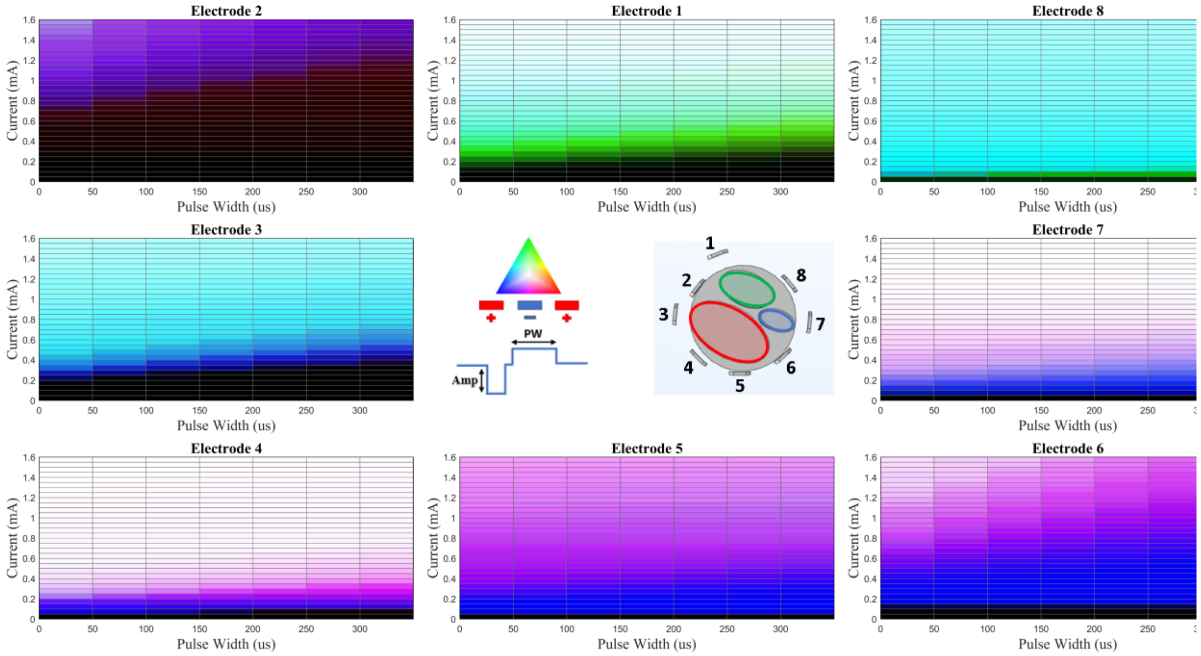


**Figure S48.** Nerve 5 – Config 4 (Simulation)

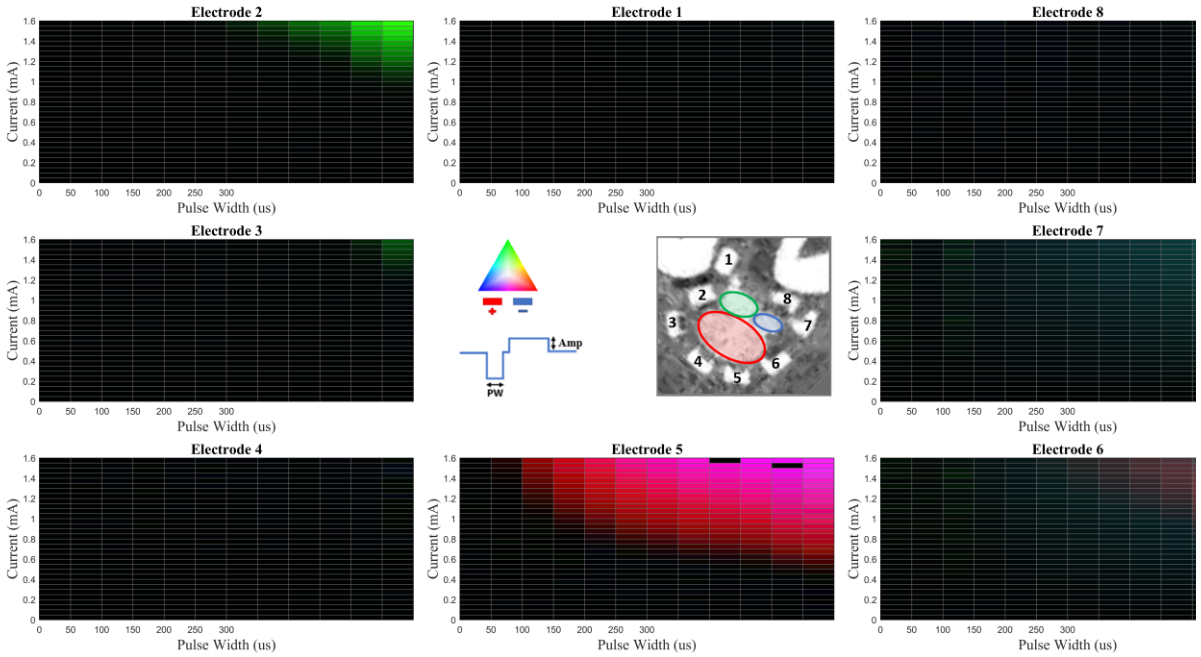

**Figure S49.** Nerve 5 – Config 5 (Ex Vivo)

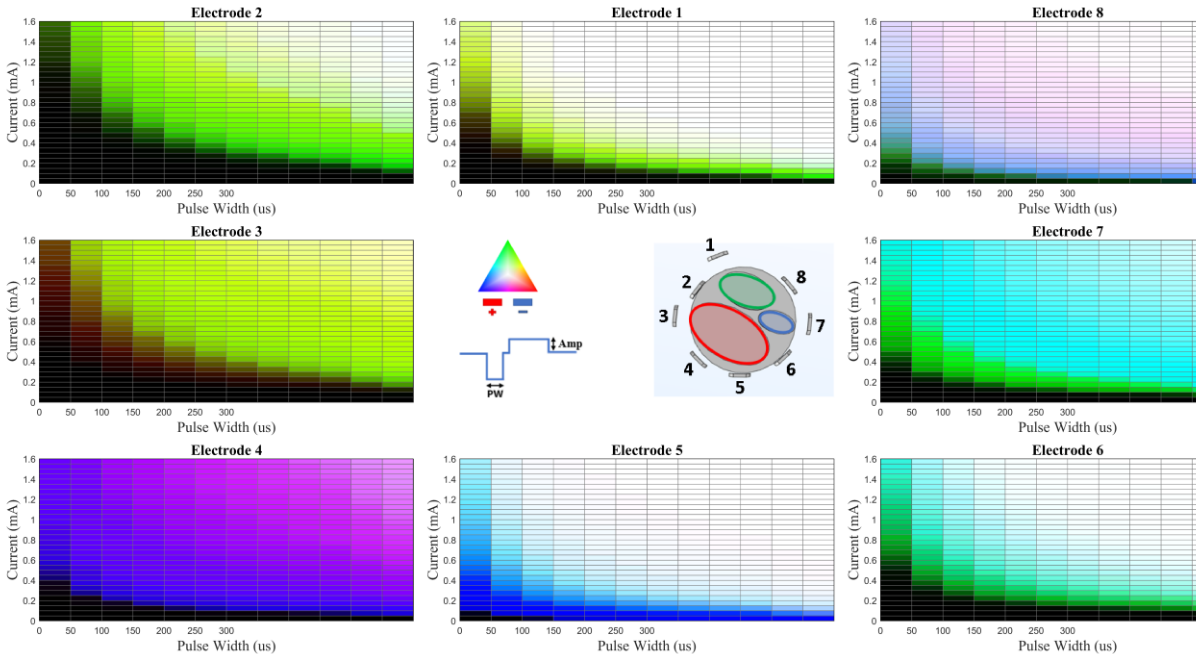

**Figure S50.** Nerve 5 – Config 5 (Simulation)

*Regression Analysis Coefficients*

**Table S1.** Amplitude Phase 1

| Fascicle | Model | N | Coefficient | p |
|---|---|---|---|---|
| Overall | ExVivo | 70827 | 0.0001292 | <0.001 |
| Overall | Simulation | 64296 | -6.376e-05 | <0.001 |
| Peroneal | ExVivo | 23767 | 9.624e-05 | <0.001 |
| Peroneal | Simulation | 17483 | 2.206e-05 | 0.016 |
| Sural | ExVivo | 32465 | 0.000139 | <0.001 |
| Sural | Simulation | 30022 | -0.0001412 | <0.001 |
| Tibial | ExVivo | 14595 | 0.0001095 | <0.001 |
| Tibial | Simulation | 16791 | 4.64e-05 | <0.001 |

**Table S2.** Amplitude Phase 2

| Fascicle | Model | N | Coefficient | p |
|---|---|---|---|---|
| Overall | ExVivo | 70827 | 6.969e-05 | <0.001 |
| Overall | Simulation | 64296 | -5.662e-05 | <0.001 |
| Peroneal | ExVivo | 23767 | 0.0001234 | <0.001 |
| Peroneal | Simulation | 17483 | -7.867e-05 | <0.001 |
| Sural | ExVivo | 32465 | 1.958e-05 | <0.001 |
| Sural | Simulation | 30022 | -3.471e-05 | <0.001 |
| Tibial | ExVivo | 14595 | 9.532e-05 | <0.001 |
| Tibial | Simulation | 16791 | -5.407e-05 | <0.001 |

**Table S3.** Nerve ID

| Fascicle | Model | N | Coefficient | p |
|---|---|---|---|---|
| Overall | ExVivo | 70827 | -0.01681 | <0.001 |
| Overall | Simulation | 64296 | 0.01793 | <0.001 |
| Peroneal | ExVivo | 23767 | -0.01535 | <0.001 |
| Peroneal | Simulation | 17483 | 0.01758 | <0.001 |
| Sural | ExVivo | 32465 | -0.03248 | <0.001 |
| Sural | Simulation | 30022 | 0.03773 | <0.001 |
| Tibial | ExVivo | 14595 | 0.04673 | <0.001 |
| Tibial | Simulation | 16791 | -0.006423 | <0.001 |

**Table S4.** Average Electrode–Fascicle Distance

| Fascicle | Model | N | Coefficient | p |
|---|---|---|---|---|
| Overall | ExVivo | 70827 | -0.1221 | <0.001 |
| Overall | Simulation | 64296 | 0.1229 | <0.001 |
| Peroneal | ExVivo | 23767 | -0.06913 | <0.001 |
| Peroneal | Simulation | 17483 | 0.02507 | 0.024 |
| Sural | ExVivo | 32465 | -0.1721 | <0.001 |
| Sural | Simulation | 30022 | 0.3197 | <0.001 |
| Tibial | ExVivo | 14595 | -0.1372 | <0.001 |
| Tibial | Simulation | 16791 | 0.1737 | <0.001 |

**Table S5.** Charge Injection

| Fascicle | Model | N | Coefficient | p |
|---|---|---|---|---|
| Overall | ExVivo | 70827 | -1.127e-07 | <0.001 |
| Overall | Simulation | 64296 | -5.496e-07 | <0.001 |
| Peroneal | ExVivo | 23767 | -1.683e-07 | <0.001 |
| Peroneal | Simulation | 17483 | -4.426e-07 | <0.001 |

| | | | | |
|---|---|---|---|---|
| Sural | ExVivo | 32465 | -1.751e-07 | <0.001 |
| Sural | Simulation | 30022 | -7.783e-07 | <0.001 |
| Tibial | ExVivo | 14595 | 2.77e-07 | <0.001 |
| Tibial | Simulation | 16791 | -4.895e-07 | <0.001 |

**Table S6.** Pulse Width Phase 1

| Fascicle | Model | N | Coefficient | p |
|---|---|---|---|---|
| Overall | ExVivo | 70827 | 0.0002483 | <0.001 |
| Overall | Simulation | 64296 | -8.609e-05 | 0.003 |
| Peroneal | ExVivo | 23767 | 0.0001851 | <0.001 |
| Peroneal | Simulation | 17483 | 0.0002443 | <0.001 |
| Sural | ExVivo | 32465 | 0.0003284 | <0.001 |
| Sural | Simulation | 30022 | -0.0003339 | <0.001 |
| Tibial | ExVivo | 14595 | 1.401e-05 | 0.753 |
| Tibial | Simulation | 16791 | 0.000216 | <0.001 |

**Table S7.** Pulse Width Phase 2

| Fascicle | Model | N | Coefficient | p |
|---|---|---|---|---|
| Overall | ExVivo | 70827 | -9.935e-06 | 0.234 |
| Overall | Simulation | 64296 | -5.817e-05 | <0.001 |
| Peroneal | ExVivo | 23767 | 6.792e-05 | <0.001 |
| Peroneal | Simulation | 17483 | -0.0001895 | <0.001 |
| Sural | ExVivo | 32465 | -7.35e-06 | 0.518 |
| Sural | Simulation | 30022 | 3.304e-05 | 0.114 |
| Tibial | ExVivo | 14595 | 4.379e-05 | 0.043 |
| Tibial | Simulation | 16791 | 6.725e-06 | 0.748 |

**Table S8.** Sink Electrode ID

| Fascicle | Model | N | Coefficient | p |
|---|---|---|---|---|
| Overall | ExVivo | 70827 | 0.0006598 | 0.070 |
| Overall | Simulation | 64296 | 0.01441 | <0.001 |
| Peroneal | ExVivo | 23767 | -0.003777 | <0.001 |
| Peroneal | Simulation | 17483 | 0.01443 | <0.001 |
| Sural | ExVivo | 32465 | 0.001409 | 0.004 |
| Sural | Simulation | 30022 | 0.02965 | <0.001 |
| Tibial | ExVivo | 14595 | 0.002518 | 0.002 |
| Tibial | Simulation | 16791 | 0.0112 | <0.001 |

**Table S9.** Source Electrode ID

| Fascicle | Model | N | Coefficient | p |
|---|---|---|---|---|
| Overall | ExVivo | 70827 | 0.007555 | <0.001 |
| Overall | Simulation | 64296 | -0.04481 | <0.001 |
| Peroneal | ExVivo | 23767 | 0.01047 | <0.001 |
| Peroneal | Simulation | 17483 | -0.0866 | <0.001 |
| Sural | ExVivo | 32465 | 0.008749 | <0.001 |
| Sural | Simulation | 30022 | -0.009134 | 0.032 |
| Tibial | ExVivo | 14595 | 0.03601 | <0.001 |
| Tibial | Simulation | 16791 | -0.05626 | <0.001 |

**Table S10.** Waveform Configuration ID

| Fascicle | Model | N | Coefficient | p |
|---|---|---|---|---|
| Overall | ExVivo | 70827 | -0.00417 | <0.001 |
| Overall | Simulation | 64296 | 0.0008498 | 0.272 |

| | | | | |
|---|---|---|---|---|
| Peroneal | ExVivo | 23767 | -0.01062 | <0.001 |
| Peroneal | Simulation | 17483 | 0.001421 | 0.303 |
| Sural | ExVivo | 32465 | -0.0009426 | 0.196 |
| Sural | Simulation | 30022 | 0.0047 | <0.001 |
| Tibial | ExVivo | 14595 | -0.004804 | <0.001 |
| Tibial | Simulation | 16791 | -0.007566 | <0.001 |
